# Ex utero development of mouse embryos from fertilization to mid-gestation

**DOI:** 10.64898/2026.11.17.767265

**Authors:** Gulben Gurhan, Alperen Yilmaz, Dmitry Lokshtanov, Mohammad Ghosheh, Francesco Roncato, Juana De La O, Lihua Wang, Xiushuai Yang, Castle Raley, Inbal Serfaty, Nizar Khatib, Reli Rachel Kakun, Nassrala Zawahri, Suhail Elias, Sergey Viukov, Itay Maza, Bernardo Oldak, Noa Novershtern, Alejandro Aguilera-Castrejon, Jacob H. Hanna

## Abstract

Early mammalian development has largely been studied as two periods divided by implantation: a pre-implantation period, when the embryo transits freely through the maternal reproductive tract, and a post-implantation period, when it develops within the uterine tissues. Although robust culture methods for mammalian pre-implantation^1,2^ or post-implantation^3–9^ embryos have been established, the continuous transition from the pre-implantation zygote to organogenesis has remained recalcitrant to faithful capture in vitro in mammalian species^10–15^. Here we establish **PR**E-implantation-**I**nitiated **Ex Ut**ero culture (**PRI-ExUt**), a sequential platform supporting continuous development of natural mouse embryos from the zygote to the E9.5 organogenesis stage, spanning nearly half of gestation without maternal interaction, embedding within exogenous extracellular matrices or co-culture with endometrial organoids. Cultured zygotes developed in vitro into blastocysts, formed egg cylinders, completed gastrulation, and progressed through neurulation, somitogenesis, axial turning and neural tube closure, generating embryonic and extra-embryonic tissues with morphology and single-cell transcriptional identity highly similar to those of stage-matched natural embryos developed in utero. We harnessed this platform as a functional benchmark for mouse stem-cell-derived blastoids and found that supplementation with an in vitro-derived OCT4+GATA6+NANOG− primitive endoderm-like population substantially extended their developmental potential, enabling bona fide mouse blastoids to reach mid-gastrulation egg-cylinder stages, contingent upon application of PRI-ExUt approach. Collectively, our findings demonstrate that mouse embryos can self-organize faithfully from fertilization to mid-gestation independently of maternal input and establish a continuous experimental window into the first half of mouse embryogenesis. By supporting continuous ex utero development from the zygote to E9.5 in the mouse (corresponding to human Carnegie stages 12–13, 26–32 days post fertilization), this platform offers a flexible framework for evaluating the developmental potential of mouse pre-implantation embryos and embryo models, and informs ongoing international ethical discussions and legislative efforts to revisit the limits placed on human embryo in vitro culture beyond day 14.

## Introduction

The transformation of a fertilized mammalian egg into a patterned embryo requires a succession of morphogenetic and lineage decisions that occur during an interval of profound experimental inaccessibility^16,17^. In mice, the fertilized zygote gives rise to a blastocyst after the first 3 days of embryogenesis^9^. The late blastocyst implants into the uterus about 4 days post-fertilization (d.p.f.) and reorganizes itself into an egg-cylinder-shaped structure, which then undergoes gastrulation between days 6 and 8, followed by neurulation and organogenesis^18–22^. These events define the architecture of the mammalian body, yet they are highly dependent on maternal uterine environment and tissues^23,24^, making them difficult to observe, perturb and interrogate over continuous developmental time, even in rodents.

^26–29^Pre–implantation culture enables zygotes to develop to blastocysts in vitro^2^, whereas post-implantation whole-embryo culture^3,8,10,25^ supports growth of natural embryos isolated after implantation, through organogenesis and fetal development^3,5^. However, the continuous transition from the pre-implantation zygote/blastocyst to organogenesis stages has remained recalcitrant to faithful capture in vitro in mammalian species. Ex utero embryogenesis, when starting from post-implantation mouse embryos, has revealed an extensive self-organizing capacity and intrinsic developmental timing in mammalian embryos and post-implantation Stem-Cell-Based Embryo Models (SCBEMs), suggesting that core programs of post-implantation embryogenesis can proceed outside maternal tissues for extended developmental intervals^3,5,25–28^. A gap nonetheless remains in capturing authentic in vitro development from pre-implantation to post-implantation stages, specifically, zygotic embryos competent to further develop adequately through to organogenesis entirely ex utero. Therefore, a remaining central question in developmental biology is <u>whether the maternal environment during implantation</u>, which provides physical, endocrine, immunological and metabolic support^23,29–31^, <u>is strictly required</u> for mammalian development to proceed faithfully towards organogenesis, or whether the embryo itself encodes intrinsic morphogenetic programs sufficient to support autonomous development in the complete <u>absence of maternal uterine</u> <u>interaction from the onset of embryogenesis.</u>

Although recent elegant in vitro culture systems have enabled mouse and human blastocysts to reach early post-implantation-like stages up to the onset of gastrulation, these protocols have failed to faithfully recapitulate defining features of bona fide of peri gastrulation stages or sustain faithful development beyond early gastrulation^10–12,15,32,33^. Even techniques that involve surgical excision of the mural trophoblast compartment or embedding into implantation-like promoting matrices have not so far extended the in vitro developmental potential of mouse blastocysts, but rather partially reduced in vitro delays in developmental timings observed in vitro in such protocols^14,15^. It is possible that the use of ingredients such as knockout serum replacement (KSR) or embedding in high-concentration Matrigel, both shown to severely impair natural mouse embryo growth ex utero^3^, and/or by not utilizing specialized whole embryo roller culture systems, shown to boost mouse embryo developmental potential through organogenesis^3^, profoundly limited the embryo integrity and developmental progression achieved when culturing mouse blastocysts in such settings^10–15,34^. Utilizing endometrial organoids^35,36^ facilitated sophisticated modelling of implantation related events of human blastocysts in an engineered system, however such post-implanted human blastocyst showed only early signs of post-implantation development, likely because once the blastocysts implanted deeply within the endometrial organoids, they became mechanically restricted from further expansion by the resident organoid cells^37,38^. In parallel, SCBEMs cultured ex utero have begun to recapitulate selected aspects of pre-implantation or post-implantation development^25,26,28,39–54^. However, SCBEMs that transcriptionally and morphologically resemble mouse blastocysts (i.e. blastoids) have shown restricted capacity to progress faithfully into post-gastrulation developmental stages^41,49,55,56^. Recent studies devised alternative methods to generate bona fide mouse blastoids and applied them to a variety of experimental settings. Notably, however, when aiming specifically to support progression to post-implantation stages, the authors could not use the novel approaches and culture conditions they developed to generate their bona fide blastoids^57–60^, instead, they modified their newly described protocols toward conditions highly similar to those previously shown to generate ETX/SEM-like structures^25,61^. The latter type of SCBEMs (ETX/SEM), are generated from pre-implantation starting cells, can progress into post-implantation gastrulating structures without first passing through a blastocyst-like stage or structure. Moreover, protocols that generate blastoids using alternative totipotent-like cells^55–58,60,62^, do not model the zygote-to-blastocyst transition, as they do not generate blastomeres or early/late morulae during these regimens. Thus, these approaches remain discontinuous through the zygote to post-implantation/gastrulation transitions. Non-human primate (NHP) blastocysts have also been cultured further in vitro, but the resulting embryos uniformly displayed abnormal morphogenesis. While these groundbreaking achievements can still be helpful for studying certain aspects of the developing NHP embryos, the overall developmental fidelity is difficult to assess rigorously because access to stage-matched post-implantation primate embryos in utero remains limited^63–66^.

Given these limitations in human and NHP models, it has become more evident that the mouse represents the only mammalian system in which natural embryos, advanced post-implantation ex utero embryo culture, stem-cell-derived embryo models, and single-cell reference datasets^67,68^ provide a highly calibrated reference for accurately assessing the extent of developmental fidelity of cultured embryos in any experimental setting. Thus, defining the limits, timing, efficiency and reliability of zygote-to-organogenesis culture in the mouse can provide a biological foundation for interpreting, improving and ethically justifying analogous approaches in other mammalian species, including human embryos and embryo models. This benchmark is particularly relevant given that human pre-implantation embryos are the only accessible living material for studying early human embryogenesis, and progression to gastrulation or organogenesis cannot be evaluated with the same experimental depth against stage-matched natural embryos because such embryos are of limited availability in humans owing to justified ethical and technical restrictions.

## Results

### Endowing cultured mouse zygotes with ability to proceed towards gastrulation ex utero

Given the above current state of the art, in this study we set out to test the feasibility of establishing a culture system capable of supporting faithful and uninterrupted embryo development from pre-implantation zygotic stage to organogenesis, which was eventually named upon concluding optimizations described below as **PR**E-implantation-**I**nitiated **Ex Ut**ero culture **(PRI-ExUt)** (**Extended Data Fig. 1**). Throughout this study, we sought to authentically capture each critical developmental step between zygote and organogenesis (developmental steps i–iv below). We performed a focused iterative screen for the most suitable base media, supplements, serum sources, hormonal cues, substrate coatings, and three-dimensional hydrogel embedding to promote adequate progression from (i) zygote to late blastocyst, (ii) late blastocyst attachment and development to egg cylinder formation (referred to as In Vitro Implantation-like (IVI-like) step), (iii) gastrulation completion and then (iv) advancing through organogenesis. When relevant, we evaluated previously described protocols that achieved limited success at specific developmental stages and/or tested whether components used in those studies could improve the corresponding stages we aimed to optimally capture in vitro. Mouse zygotes were collected at E0.5 and cultured in a variety of conditions. Basal media used for culturing naive mouse ESCs were detrimental to pre-implantation embryo development (**Extended Data Fig. 2A**), while continuous single-step medium (Continuous Single Culture-NX Complete (CSCM-NXC)) allowed robust growth until the late blastocyst stage E4.5 (**Fig. 1A-B**). Under the latter selected condition for further experimentation, at E3.5 embryos formed blastocysts with a visible blastocoel and inner cell mass, and by E4.5 displayed round expanded blastocyst morphology with appropriate segregation between epiblast, primitive endoderm and trophectoderm lineages, as assessed by immunostaining (**Extended Data Fig. 2B-E**). KSOM medium is also widely used for culturing mouse embryos from the zygote to E3.5–E4.5, but it was less efficient in promoting hatching at E4.5 than CSCM-NXC, and thus CSCM-NXC was chosen for further experimentation (**Extended Data Fig. 2F)**.

**Figure 1.**
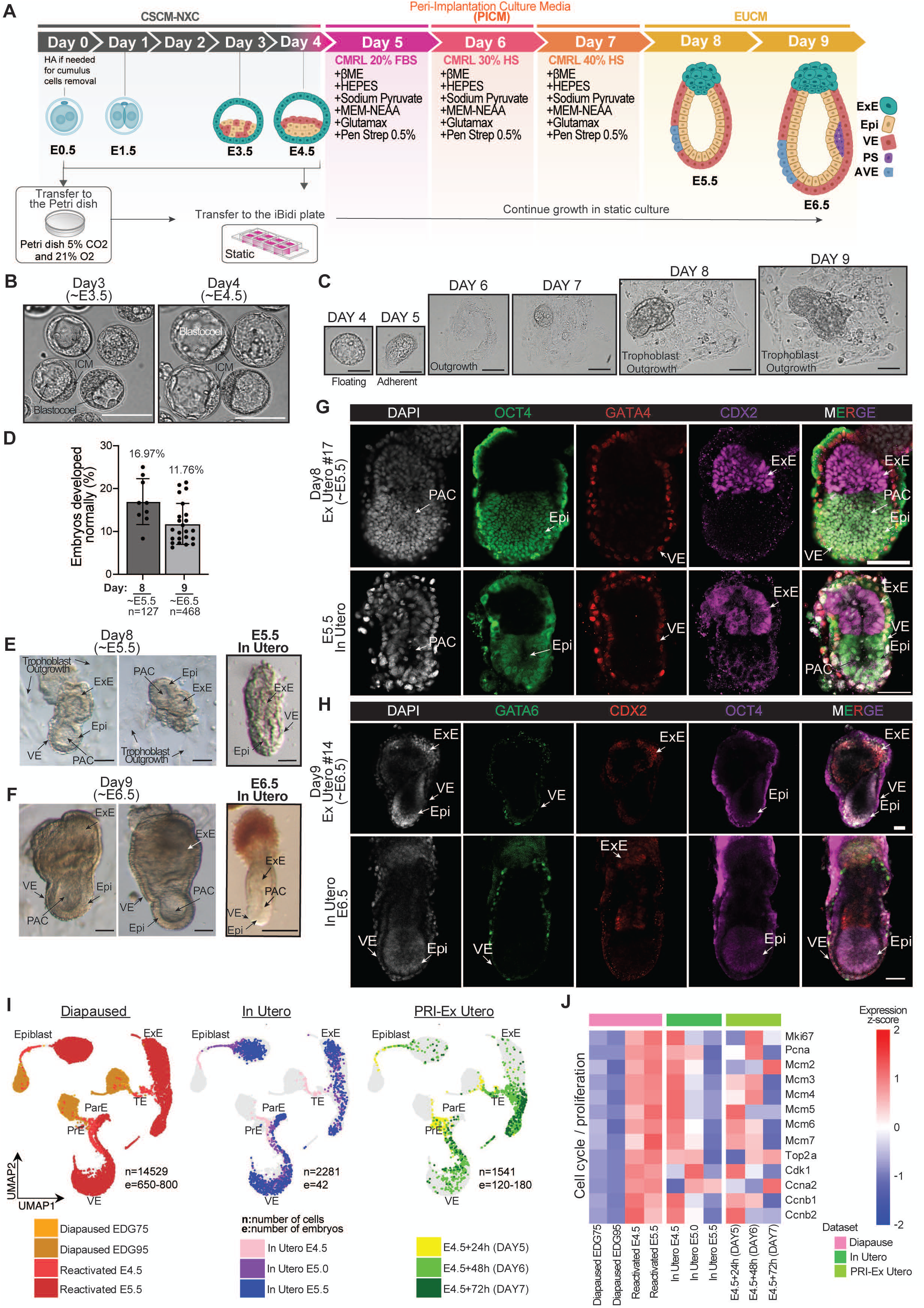
Continuous culture of mouse embryos from the totipotent zygote to egg-cylinder stages. **A,** Schematic illustration of the ex utero culture conditions from E0.5 zygotes to E6.5 egg-cylinder embryos, showing sequential CSCM-NXC, CMRL-based implantation media, which we termed PICM, and EUCM post-implantation conditions. **B,** Representative images of E3.5 and E4.5 stage blastocysts grown from cultured zygotes. Scale bars, 100μm. **C,** Morphological progression from blastocysts to egg-cylinder, culture days 4 to 9, following transfer of E4.5 embryos to in vitro implantation conditions. Whole progression takes place in the same iBidi plate well and follows the same embryo. Scale bars, 200 μm. **D,** Quantification of properly developed embryos at E5.5 (day 8) and E6.5 (day 9) stages. Each point represents an independent experiment; total numbers of embryos are indicated. The numerical values represented by bars are indicated on top of the error bars. **E, F,** Representative bright-field images of day 8 (∼E5.5) and day 9 (∼E6.5) PRI-ExUt embryos compared with stage-matched in utero embryos. Scale bars, 100 μm. **G,** Whole-mount immunostaining of day 8 PRI-ExUt cultured embryos and developmentally matched in utero E5.5 embryos stained for OCT4 (Epi), and GATA4 (VE) and CDX2 (ExE). **H**, Whole-mount immunofluorescence of day 9 PRI-ExUt and E6.5 in utero embryos for GATA6 (VE), CDX2 (ExE) and OCT4 (Epi). Scale bars, 100 μm. **I**, UMAP representation of integrated single-cell RNA-seq datasets for diapaused^69^, in utero peri-implantation^68^ and PRI-ExUt mouse embryos at the indicated time-points. Cluster annotation is based on in utero data annotation. Diapause samples (Diapaused EDG75 and Diapaused EDG95) are clustered separately from the non-diapause samples, suggesting a profound difference in gene expression between the cell states. Number of cells (indicated as ‘n’) and embryos (indicated as ‘e’) analyzed are indicated. Heatmap showing normalized expression (z-score) of cell cycle and proliferation-associated genes in epiblast cells from diapause, in utero and PRI-ExUt samples. In-utero and PRI-ExUt epiblast cells do not show the typical coordinated repression of cell cycle genes, that can be observed only in the diapause cells. Abbreviations: AVE, anterior visceral endoderm; EDG75 and EDG95, estimated day of gestation 7.5 and day 9.5; Epi, epiblast; ExE, extraembryonic ectoderm; FBS, fetal bovine serum; HA, hyaluronidase; HS, human serum; ICM, inner cell mass; PAC, pro-amniotic cavity; ParE, parietal endoderm; PrE, primitive endoderm; PS, primitive streak; TE, trophectoderm; VE, visceral endoderm.

When considering conditions for the next stage, IVI-like step, we focused on the most recently established IVI-like method to culture pre-implanted embryos (E3.5) into pre-gastrulating egg-cylinder-shaped embryos (up to E6–E6.5) over a protocol lasting 5-6 days ^10^. This was done by devising an in vitro culture (IVC) media either supplemented with FBS (IVC1) or knockout serum-replacement (IVC2) as originally described in the same study^15^. We therefore tested whether incorporating this previously stablished protocol would overcome this experimental obstacle. Embryos grown according to this protocol reached an E6.5-like stage^10^; however, as indicated in the original report, such embryos had altered morphology and smaller size than their in vivo counterparts, which likely explains why, in our hands, they were unable to develop beyond E6 stage, despite additional tests and modifications attempted herein. For example, further culture of egg-cylinder embryos beyond ∼E6.5-stage using IVC2 media resulted in restricted epiblast growth (**Extended Data Fig. 3A**), underscoring the need for further evaluation and revision of IVI-like conditions.

Unlike previous protocol in which E3.5 blastocysts were chosen as the starting material^15^, we found that it was preferable in subsequent culture optimizations to transfer zygote-derived E4.5 (protocol day 4) blastocysts that become zona-free through the partial or complete natural zona hatching that frequently occurs at E4.5, as this better allowed attachment to iBidi culture plates (**Fig. 1C**, **Extended Data Fig. 1B (**D4 timepoint)), which we found optimal for trying to recapitulate IVI-like conditions starting with such E4.5 blastocysts. Replacing IVC2-30% KSR media, previously described to culture mouse E3.5 blastocysts to ∼E6^15^, with EUCM medium containing either rat serum (RS) or human serum (HS), improved the morphological features of the egg-cylinder embryos obtained, generating egg-cylinders that more closely resembled their in utero counterparts in shape and size, yet at lower efficiency (**Extended Data Fig. 3B**). Thus, we sought to optimize a regimen that supported improved egg-cylinder formation efficiency and quality which was not achieved with EUCM that is DMEM based. We identified CMRL as most suitable basal media to achieve this goal, as it had previously been used to promote human blastoid growth towards early post-implantation stage^44^. We combined CMRL with serum instead of KSR to formulate what we term as Peri-Implantation Culture Media (**PICM**) (**Fig. 1A**). PICM was applied from day 4.5 through the end of day 7 while the embryos were cultured on uncoated iBidi plates, providing optimal IVI-like progression as part of the PRI-ExUt platform formulated herein. In this optimized PICM-based protocol, E4.5 blastocysts moved to iBidi plates were cultured for 36 hours in PICM containing 20% fetal bovine serum (FBS) (culture day 4), followed by 24 hours in PICM supplemented with 30% HS, and subsequent 24 hours in PICM containing 40% HS (**Fig. 1A**). This sequence of supplemented media promoted trophectoderm differentiation and attachment, trophoblast outgrowth, expansion of embryonic and extraembryonic compartments, and formation of a pro-amniotic cavity (**Fig. 1C**). Two days after transfer to PICM (day 6), attached embryos produced trophoblast outgrowths surrounding a growing inner mass (**Fig. 1C, Supplementary Video 1**). Remarkably, egg cylinders emerged after day 7 of the PICM regimen. Once egg-cylinder structures were observed, embryos were maintained in the same iBidi plates under EUCM conditions for the remaining culture days of the PRI-ExUt platform. EUCM had previously been developed to support ex utero development of natural post-implantation egg-cylinder stage embryos through gastrulation and organogenesis (**Fig. 1A)**. By day 8 of the protocol, corresponding approximately to E5.5 of in utero development, embryos had formed cup-shaped egg-cylinder structures containing organized OCT4^+^ epiblast (Epi) cells, GATA4^+^ visceral endoderm (VE) cells, and CDX2^+^ extra-embryonic ectoderm (ExE) cells, closely resembling stage-matched in utero embryos with an efficiency close to 17% (**Fig. 1D-E,G)**.

We note that the PICM based IVI-like regimen was established through multiple iterative tests and optimizations, most noteworthy of which is that Matrigel- and collagen-coating of iBidi plates during the IVI-like stage (days 5–7) was detrimental to egg-cylinder formation, whereas uncoated plates produced the best outcomes. (**Extended Data Fig. 4A**). Hypoxic (5% O_2_) conditions during pre-implantation culture were not superior for growth of in vitro implanted embryos compared to 20% oxygen, thus the latter was chosen for further experimentation (**Extended Data Fig. 4B**). Supplementation of PICM conditions with uteroglobin, IGF, FGF, triiodothyronine (T3), ROCK inhibitor (Y-27632), FGF2, Glucose, N2 supplement, B27 supplement, eIVC media including a version that is pre-conditioned for 24 hours on irradiated mouse fibroblasts (**See Methods**), or previously described IVC supplements^15^, including progesterone, ITS-X, β-estradiol and N-acetylcysteine, either did not further improve, or impaired, development to egg-cylinder stages (**Extended Data Fig. 5A-C**). Including HEPES, which had previously been found beneficial in EUCM medium used for promoting egg-cylinder-based gastrulation^3^, sodium pyruvate, NEAA, GlutaMAX, and no more than 0.5X Pen-Strep antibiotic, were all found to be required for an optimal outcome in PICM conditions (**Extended Data Fig. 4C**). Continuing the CSCM-NXC condition for 24 h after placing E4.5 blastocysts in iBidi plates and starting PICM on day 5 yielded overall similar results to the finalized protocol regimen in which PICM was started 24 hours earlier (**Extended Data Fig 5A (top rows), Extended Data Fig. 5B,D**).

On culture day 9, approximately equivalent to E6.5, ∼12% of cultured embryos showed proper growth and further elongation of the egg-cylinder structure, containing OCT4^+^ Epi, GATA6^+^ VE and CDX2^+^ ExE compartments, and proximal–distal embryo length was indistinguishable from that of in utero E6.5 controls (**Fig. 1F, H, Extended Data Fig. 3C, Extended Data Fig. 6A-D**). At this stage, BRACHYURY expression emerged within the posterior epiblast region, indicating symmetry breaking and primitive-streak initiation (**Extended Data Fig. 6D**). These results confirmed that embryos isolated at the zygote stage can be continuously cultured through blastocyst formation, implantation-like morphogenesis, egg-cylinder formation and the onset of gastrulation entirely ex utero.

Given that E4.5 blastocysts required three additional days in culture to progress to an E5.5 egg cylinder stage that could still develop normally to E6.5 and beyond, we wondered whether the ability to tolerate such a delay in blastocyst development during the IVI-like step resulted from PICM being able to induce the blastocysts to enter an in vitro induced diapause-like state^69^. Embryonic diapause is a reversible state of pre-implantation blastocyst dormancy characterized by a halt in developmental progression under stress conditions, while preserving the embryo’s capacity to reactivate and resume development when permissive conditions are restored^70,71^. Mouse diapause is characterized by unique molecular signatures involved in inducing and maintaining such a unique embryonic state, and can also be induced and modeled via in vitro grown ESCs and blastocysts^69^. To investigate this question, we performed single-cell RNA sequencing at +24, +48 and +72 hours after applying PICM based IVI-like culture on E4.5 blastocysts, and integrated these data with previously published diapause^69^ and in utero reference datasets^68^. We identified 9 different cell clusters, including Epiblast, Primitive endoderm (PrE), parietal endoderm (ParE), visceral endoderm (VE), as well as trophectoderm (TE) and extraembryonic ectoderm (ExE) (**Fig. 1I, Extended Data Fig. 7A**). Our analyses showed that in vitro cultured embryos at +24 h and +48 h remained transcriptionally less mature than in utero embryos, slowly progressing from E4.5 toward E5.5 through culture time (**Fig. 1I, Extended Data Fig. 7B**). By +72 hours after implantation in vitro, PRI-ExUt embryos progressed to an E5.5 state. Notably, when comparing transcriptional profiles to embryos which were induced into diapause at E4.5, diapaused cells from all three blastocyst lineages (Epi, ExE and PrE/VE) clustered separately from both in utero and PRI-ExUt cells (**Extended Data Fig. 7B**), indicating that PRI-ExUt embryos do not acquire a diapause-like transcriptional profile in any of these lineages at the stages examined (**Figure 1I, Extended Data Fig. 7B**). Furthermore, analysis of diapause-associated cell cycle and proliferation gene signatures demonstrated that such pattern was not observed in PRI-ExUt samples either (**Fig. 1J**). Further comparison of other diapause-associated transcriptomic modules, including cell-cycle arrest, mTOR/growth, MYC/biosynthetic pathways, lipid metabolism, autophagy, and YAP/TAZ-associated programs did not reveal a molecular state characteristic of diapause during PRI-ExUt culture, and showed a similar pattern to the one manifested by in-utero natural embryos, or that observed upon developmental reactivation from diapause (**Extended Data Fig. 7B, 8**). Together, these results exclude the hypothesis that PRI-ExUt embryos enter a canonical diapause state during the E4.5-E5.5 transition. Moreover, when comparing the dynamics of change in transcriptional similarity score to the final E5.5 in utero (**Extended Data Fig. 7C**), it became evident that PRI-ExUt +24 h, +48 h and +72 h samples did not show a halt in developmental progression, as known to occur during diapause. Instead, the PRI-ExUt embryos showed a gradual and continuous increase in their similarity score to E5.5 in utero stage, reflecting developmental progression at a slower pace, without any signs of halt or pausing, further arguing against the involvement of a diapause-like process in the PRI-ExUt platform. Together, the slowed developmental progression observed during the IVI-like step reveals an important developmental principle: chronological culture time becomes transiently uncoupled from developmental time during in vitro implantation-like stage. Time alignment is restored once embryos reach the egg-cylinder stage, after which PRI-ExUt embryos progress from E5.5 to E7.5 and beyond without further developmental delay.

### Cultured zygotes capture gastrulation in the PRI-ExUt platform

We next examined whether zygote-derived egg cylinders could progress further into gastrulation and adequately establish the major germ layers and extraembryonic lineages characteristic of E7.5 mouse embryos at mid-gastrulation. Day 9 PRI-ExUt E6.5 egg-cylinder embryos were maintained in EUCM for one more day (until end of Day 10) in iBidi plates (**Fig. 2A**), and displayed morphological features of successful mid-gastrulation, including an elongating primitive streak at the future posterior side, an organized epiblast compartment with an anterior head-fold region, as well as developing amnion and chorion tissues that delineated the amniotic and exocoelomic cavities (**Fig. 2B**), mirroring defining developmental milestones occurring at E7.5 days of in utero development. Whole-mount immunostaining revealed BRACHYURY^+^ primitive streak cells extending through the distal epiblast region, together with FOXA2^+^ definitive endoderm domains organized similarly to E7.5 in utero controls (**Fig. 2C-D**). Additional staining for OCT4, GATA4 and BRACHYURY confirmed the presence of epiblast, visceral/definitive endoderm and primitive streak populations (**Fig. 2D**). Extended analyses further identified FOXA2, SOX17, GATA4, SOX2 and OTX2 expression in appropriate endodermal and anterior domains, supporting correct gastrulation-stage patterning (**Extended Data Fig. 6E-G**).

**Figure 2.**
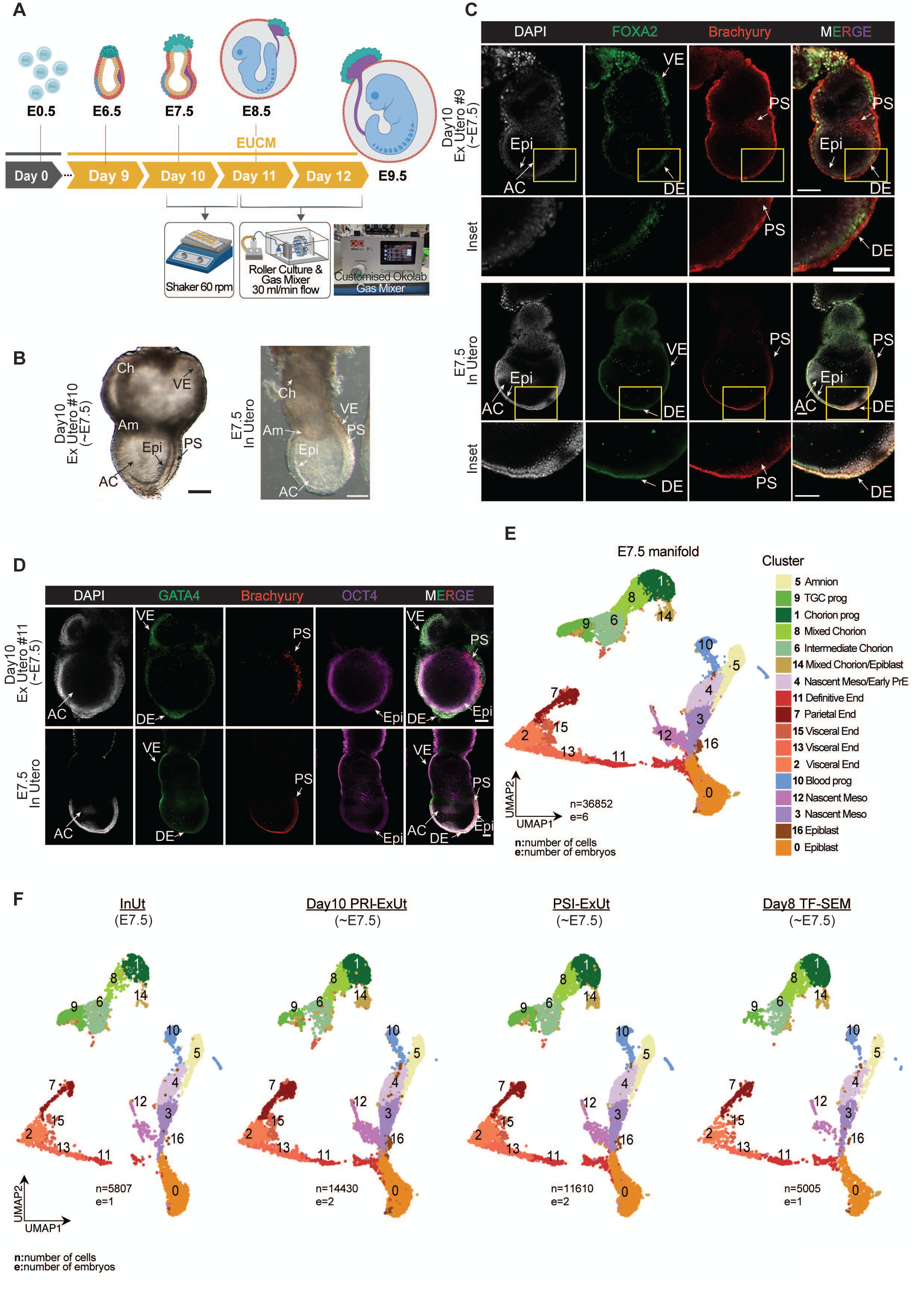
Ex utero culture media (EUCM) enables PRI-ExUt embryos to progress through gastrulation. **A, Schematic depiction of the PRI-ExUt culture method from day 9 through day 12 utilizing EUCM, including orbital-shaker** (with 60 rpm rotation speed) and roller-culture phases (30 ml/min gas flow) used for progression toward E9.5. **B,** Representative stereomicroscope images of day 10 PRI-ExUt embryos and stage-matched in utero E7.5 embryos. Scale bars, 100 μm. **C,** Representative whole-mount immunostainings of day 10 PRI-ExUt and E7.5 in utero embryos stained for FOXA2 (definitive endoderm), and BRACHYURY (Primitive Streak). Insets show a magnified view of the distal region. Scale bars, 100 μm. **D,** Whole-mount staining confocal imaging for GATA4, BRACHYURY and OCT4 in day 10 PRI-ExUt and E7.5 in utero embryos. Scale bars, 100 μm. **E,** Integrated E7.5 single cell RNA-seq UMAP manifold showing annotated cell lineages in E7.5 in utero and PRI-ExUt mouse embryos, as well as PSI-ExUt embryos and equivalent SEMs. **F,** Distribution of E7.5 in utero, day 10 PRI-ExUt, PSI-ExUt and day 8 TF-SEM cells across the integrated E7.5 manifold, showing high similarity between the cell composition of the different embryo models. Color code is as in (E). n = number of cells; e = number of embryos. Abbreviations: AC, amniotic cavity; Am, amnion; Ch, chorion; DE, definitive endoderm; Epi, epiblast; EUCM, ex utero culture medium; ExE, extraembryonic ectoderm; InUt, in utero; Meso, mesoderm; PrE, primitive endoderm; PRI-ExUt, pre-implantation-initiated ex utero culture; prog, progenitor; PS, primitive streak; PSI-ExUt, post-implantation-initiated ex utero culture; TE, trophectoderm; TF-SEM, transgene-free stem-cell-derived embryo model; TGC, trophoblast giant cell; VE, visceral endoderm.

To evaluate developmental fidelity at single-cell resolution, we performed single-cell RNA sequencing of day 10 PRI-ExUt embryos and integrated these data with stage-matched in utero embryos. We leveraged one of the main advantages of the mouse system, which is the availability of embryos and embryo models grown ex utero obtained from various sources for transcriptomic comparison. These include natural post-implantation E6.5 embryos dissected and grown ex utero through gastrulation and beyond^3^ (**P**o**S**t-implantation-**I**nitiated **Ex Ut**ero embryos - **PSI-ExUt**), and **T**ransgene-**F**ree **S**tructured **S**tem-Cell-Based **E**mbryo **M**odels (TF-SEMs), which are derived exclusively from in vitro grown naive PSCs and upon aggregation, bypass the blastocyst-like stage, directly generating post-implantation embryo models from E5.5 up to E8.5^72^. The integrated E7.5 manifold recovered 17 clusters, covering the expected embryonic and extraembryonic transcriptional states, including epiblast, nascent mesoderm, definitive endoderm, parietal endoderm, visceral endoderm, amnion, chorion progenitors and trophoblast giant-cell progenitor populations (**Fig. 2E, Extended Data Fig. 9A**). Importantly, we found no evidence for the absence of any known cell-type cluster, nor did we detect any unidentified cell populations in PRI-ExUt E7.5 embryos that were not present in the equivalent in utero developed counterparts (**Fig. 2F, Extended Data Fig. 9B**). Cluster-level comparison of average expression per gene showed strong transcriptional concordance between PRI-ExUt and in utero embryos (Pearson R > 0.94 across clusters; **Extended Data Fig. 10A**), and similar correlation to embryos isolated at E6.5 and grown ex utero to E7.5 (PSI-ExUt) (R > 0.97; **Extended Data Fig. 10A**). Thus, in terms of transcriptional similarity and cell-state identity, initiating culture before or after implantation does not compromise the state achieved at gastrulation in ex utero grown embryos or in those assembled ex utero solely from PSCs.

### Gastrulation completion and early organogenesis initiation in PRI-ExUt culture

We next tested whether PRI-ExUt cultured embryos could complete gastrulation and advance into early organogenesis. Day 10 egg-cylinder-stage E7.5 embryos were maintained in EUCM, but had to be moved from static conditions to a dynamic orbital shaker to support their further development (**Extended Data Fig. 1**), a condition previously shown to support growth of post-implantation embryos up to E8.5^25,72^, likely by facilitating better nutrient and oxygen diffusion. By day 11 of culture, embryos in EUCM progressed to a stage corresponding approximately to E8.25-E8.5, ranging from 2 to 8 somite pairs (**Fig. 3A, Extended Data Fig. 5B and 11A**). At this stage, embryos developed within a vascularized yolk sac with circulating red blood cells and displayed prominent neural folds, a forming neural tube, tailbud, foregut pocket and a morphologically evident beating heart (**Fig. 3A, Supplementary Video 2**).

**Figure 3.**
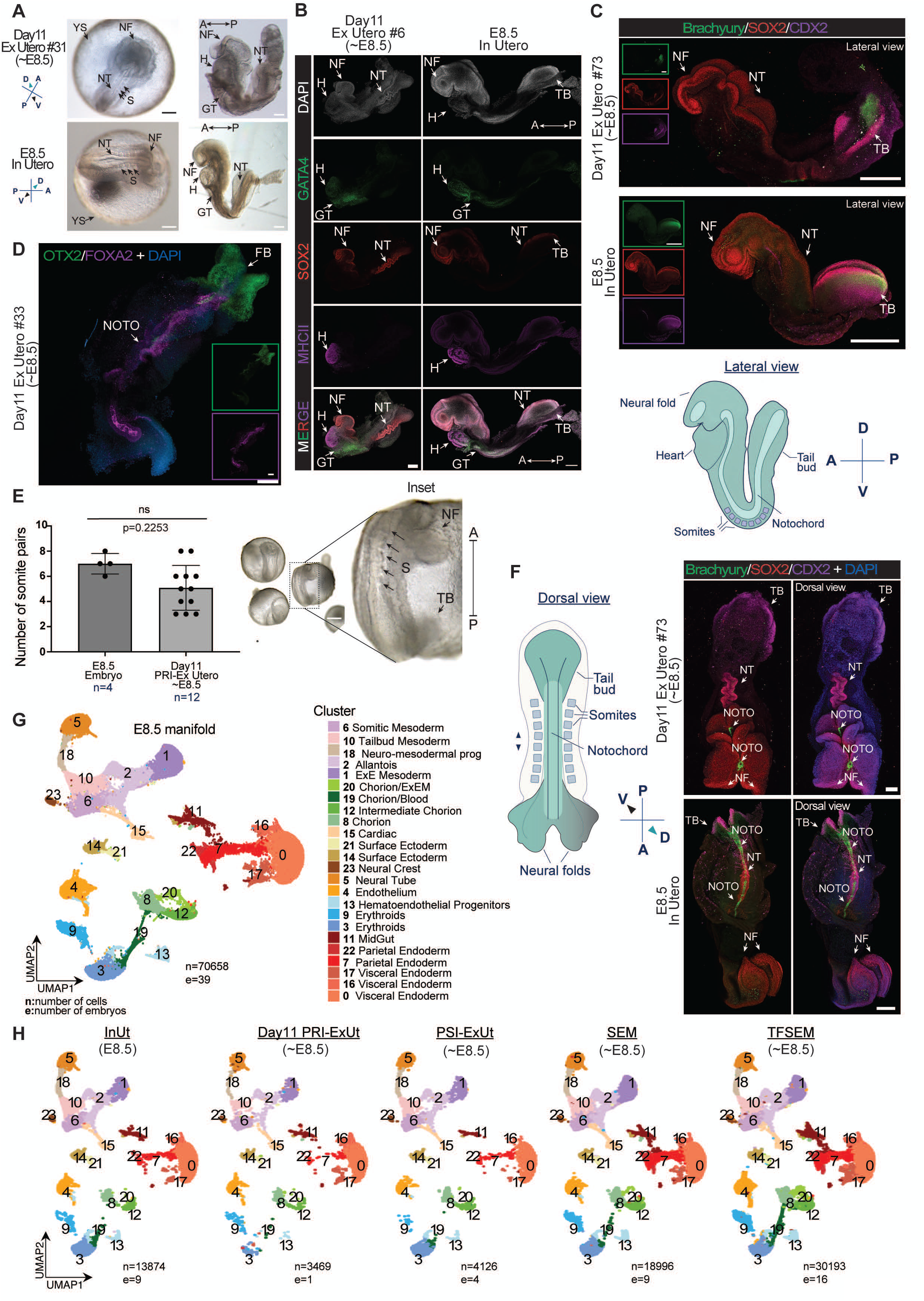
Characterization of PRI-ExUt embryos at the onset of organogenesis. **A,** Representative stereomicroscope images of day 11 PRI-ExUt and in utero E8.5 embryos shown within the yolk sac and after dissection. Scale bars, 200 μm. Axes of development, anterior (A), posterior (P), dorsal (D), and ventral (V) are indicated as shown on the compasses. **B,** Representative immunostainings of day 11 PRI-ExUt and developmentally matched E8.5 control embryos for GATA4, SOX2 and MHC-II. Scale bars represent 200 μm. **C,** Whole-mount immunostainings for BRACHYURY, SOX2 and CDX2 in day 11 PRI-ExUt and E8.5 in utero embryos. Insets show individual channels. Scale bars, 200 μm. **D,** Representative whole-mount immunofluorescence of day 11 PRI-ExUt embryo for OTX2 and FOXA2, showing anterior neural and axial domains. Scale bar, 100 μm. **E,** Somite numbers in day 11 PRI-ExUt and E8.5 in utero embryos. Right images, representative somite annotation on PRI-ExUt embryos. Unpaired two-tailed t-test was performed. Error bars represent standard deviation. Dots represent individual embryos; *n* = number of embryos quantified; ns = non-significant. Scale bar, 200 μm. p value is indicated on the graph. **F,** Representative dorsal-view immunofluorescence for BRACHYURY, SOX2 and CDX2 in day 11 PRI-ExUt and E8.5 in utero embryos. Schematic illustrates the corresponding dorsal anatomy. Scale bars, 200 μm. **G,** Integrated E8.5 single cell RNA-seq UMAP representation of single-cell RNA-seq for in utero E8.5 mouse embryos, ex utero day 11 PRI-ExUt and E8.5 equivalent SEMs showing annotated cell populations. **H,** Distribution of E8.5 in utero, day 11 PRI-ExUt, E8.5 PSI-ExUt, day 8 SEM and day 10 TF-SEM cells across the integrated E8.5 manifold, showing high similarity between the cell composition of the different embryo models. n = number of cells; e = number of embryos. Color code is as in (**G**). Abbreviations: A, anterior; D, dorsal; Epi, epiblast; ExE, extraembryonic ectoderm; FB, forebrain; GT, gut tube; H, heart; InUt, in utero; MHCII, myosin heavy chain II; NF, neural fold; NOTO, notochord; NT, neural tube; P, posterior; PRI-ExUt, pre-implantation-initiated ex utero culture; PSI-ExUt, post-implantation-initiated ex utero culture; S, somite; SEM, stem-cell-derived embryo model; TB, tail bud; TF-SEM, transgene-free stem-cell-derived embryo model; V, ventral; YS, yolk sac.

Whole-mount immunostaining confirmed the proper emergence of tissue architecture characteristic of E8.5 mouse development, adequately recapitulating to a very large extent the spatial program of in utero controls (**Fig. 3B-F, Extended Data Fig. 11**). E8.5 PRI-ExUt embryos formed prominent neural folds and a neural tube labelled by SOX2, OTX2 and FOXG1 marked the anterior neural region, while FOXA2 and BRACHYURY marked the developing notochord and posterior neuromesodermal progenitors (**Fig. 3B-D,F, Extended Data Fig. 11B-F, Supplementary Videos 3-5).** GATA4 labelled cardiac and gut endoderm-associated domains, while the embryonic heart, including emerging chamber-like morphology, was co-labelled with sarcomeric myosin II (MHC-II) (**Fig. 3B, Extended Data Fig. 11C, Supplementary Video 5**). As with in utero controls, CDX2 labelled the dorsal region at the tailbud of PRI-ExUt embryos, while the ventral area of the posterior embryonic part showed specified primordial germ cells, identified by SOX2 and OCT4 (**Fig. 3C, F, Extended Data Fig. 11B-C, Supplementary Video 4**), consistent with appropriate anterior-posterior patterning at early organogenesis stages.

Unbiased single-cell RNA sequencing of day 11 PRI-ExUt embryos further supported progression to organogenesis stages (**Fig. 3G, Extended Data Fig. 12**). Cultured embryos contained 24 clusters representing all major lineages present in E8.5 in utero embryos, spanning all three germ layers, extraembryonic tissues and their derivatives. These clusters included neural tube and neural crest, somitic and tailbud mesoderm, cardiac, midgut, as well as visceral and parietal endoderm, extraembryonic mesoderm, chorion and yolk sac lineages, hematoendothelial and erythroid compartments, and primordial germ cells, among others (**Fig. 3G, Extended Data Fig. 12**). Integration of scRNA-seq datasets from E8.5 PRI-ExUt, PSI-ExUt^3^, in utero controls and post-gastrulation TF-SEMs and SEMs^25,72^ demonstrated that, despite not developing within an in vivo uterine environment or undergoing in vivo implantation, the overall cell type distribution and proportion in PRI-ExUt cultured embryos closely resembled stage-matched natural embryos, both in utero and PSI-ExUt (**Fig. 3H, Extended Data Fig. 12B**). Notably, cluster-level transcriptomic correlation analysis showed close similarity between PRI-ExUt embryos and in utero controls (Pearson R > 0.90), as well as with embryos isolated at E6.5 and grown ex utero to E8.5 (R = 0.86–0.96), confirming high transcriptional fidelity at the onset of organogenesis (**Extended Data Fig. 10B**). These observations establish that continuous zygote-to-organogenesis culture supports proper tissue morphogenesis and molecular diversification of cell types into post-gastrulation early organogenesis-stages in the absence of in vivo uterine implantation.

### Embryo axial turning and development to E9.5 mid-gestation stage

A major developmental transition after E8.5 corresponds to axial turning, which is accompanied by continued organogenesis throughout the embryo, with particularly prominent development in the neural, cardiac, gastrointestinal, somitic and craniofacial systems. Progression beyond E8.5 in whole-embryo culture requires dynamic gas exchange and rotation to support rapidly expanding tissues^3,5^. Therefore, to sustain embryonic growth beyond E8.5-stage, we transferred day 11 PRI-ExUt embryos to a roller culture platform connected to an electronically controlled gas mixer (see **Methods**) and maintained them in EUCM for an additional 24-36 hours (**Fig. 2A, Extended Data Fig. 1**). A fraction of embryos cultured under rotating conditions progressed beyond E8.5, completed axial turning and progressed through neural tube closure, while those embryos maintained on an orbital shaker failed to turn and did not advance beyond E8.5 (**Fig. 4A, Extended Data Fig. 5E, F**). By day 12, PRI-ExUt embryos had undergone axial turning and displayed the external organization corresponding to that of their E9.5 in utero counterparts (**Fig. 4A**). Embryos were enclosed by the yolk sac and amnion, contained an extended allantois, and showed a curved body axis with the ventral region of the tail facing the ventral anterior region, as well as red blood cells circulating through the embryo and yolk sac vessels (**Fig. 4A, Supplementary Video 6**). Properly developed embryos presented all hallmarks of in utero development at E9.0-E9.5, with embryos ranging between 16-24 somites at 24-36 hours after transfer to rotating culture. The embryonic brain was regionalized, with clearly visible forebrain, midbrain and hindbrain and spinal cord. The optic vesicle, otic pit as well as up to 3 pharyngeal arches were evident, particularly the mandibular and maxillary arches (**Fig. 4A, B**). Whole-mount immunostaining for GATA4, SOX9 and TUJ1 identified properly organized gut and cardiac tissues, neural crest/chondrogenic progenitor domains, and differentiated neuronal populations, with tissue architecture closely matching in utero control embryos (**Fig. 4B, Supplementary Video 7**). In concordance with this developmental stage, the anterior neuropore was mostly closed by the end of culture, while the posterior neuropore remained open (**Fig. 4A,B**). We quantified culture efficiency by scoring stage-specific morphological landmarks (see **Methods**) from E7.5 to E9.5 (**Fig. 4C**). The morphology of the cultured embryos faithfully resembled their in utero counterparts, however, as typically observed for embryos and embryo models grown ex utero^3,5,25,61^, the proportion of correctly developed embryos in vitro, relative to the initial number of zygotes used on day 0 (we term this as absolute efficiency), declined progressively across culture time, with ∼6% reaching E7.5 morphology at day 10, ∼3% properly reaching E8.5-equivalent morphology at day 11, and ∼1% of zygotes developing properly to E9.5 after 12 days of culture (**Fig. 4C**). Importantly, given that large numbers of zygotes can be obtained per batch and correctly developed E6.5 egg-cylinder embryos can be readily identified by morphology in the same iBidi plates, we calculated efficiencies relative to this stage, which we term effective efficiency. Among egg-cylinder embryos that developed correctly to E6.5, 57.3% progressed to E7.5, 29.6% to E8.5, and 4.9% to E8.75–E9.5 (**Extended Data Fig. 4D**). Relative to correctly developed E8.5 embryos, which were selected for transfer to roller culture-based platform, 19.3% subsequently reached E9.5 (**Extended Data Fig. 4D**). Moreover, PRI-ExUt embryos that developed appropriately showed axial lengths comparable to stage-matched in utero embryos, with no significant differences at any stage between E6.5 and E9.5 (**Fig. 4D**).

**Figure 4.**
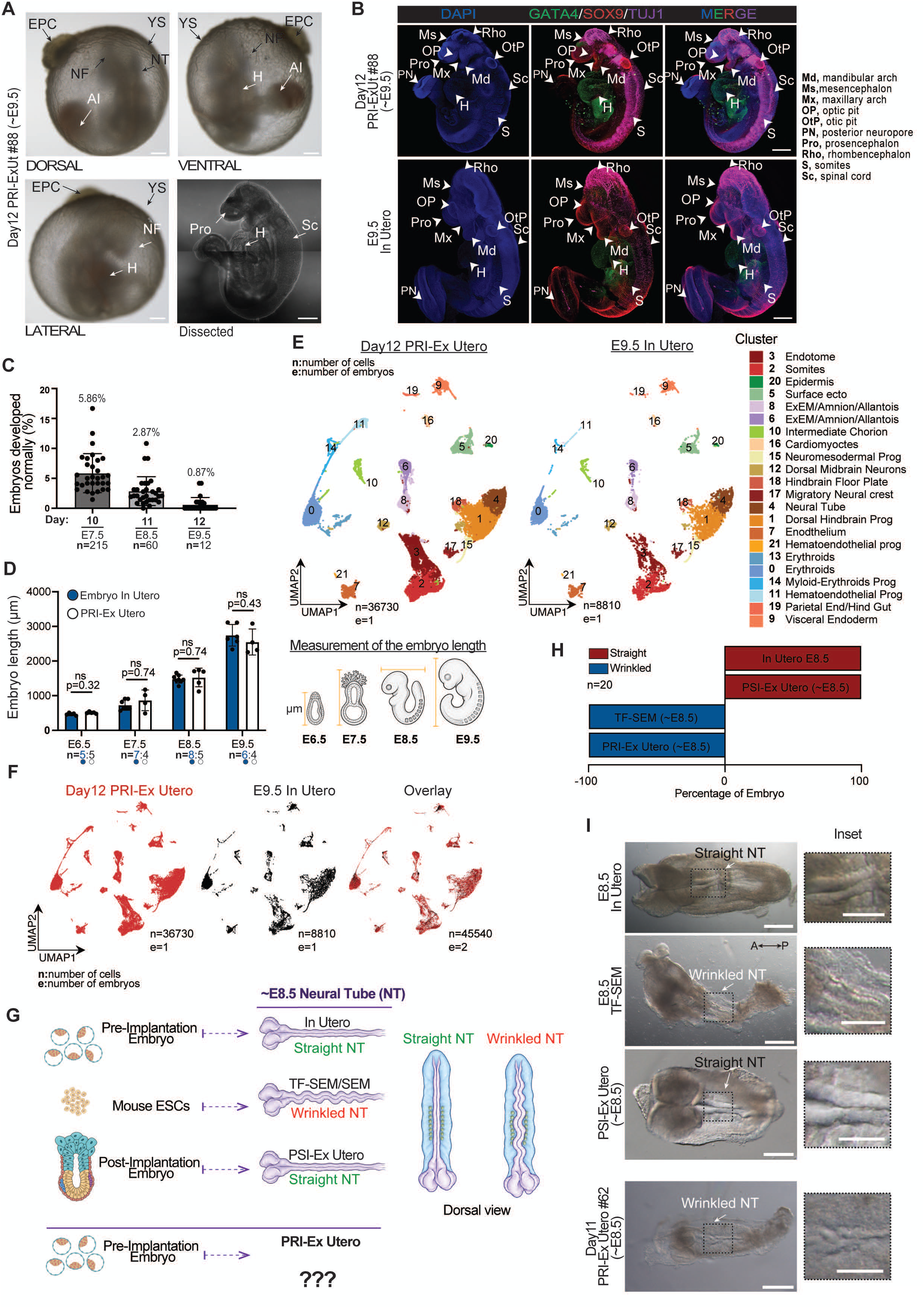
PRI-ExUt culture supports axial turning and development to E9.5. **A,** Representative E9.5 embryo morphology images of day 12 PRI-ExUt embryos shown before and after dissection from the embryonic yolk sac. Scale bars, 250 μm before dissection and 500 μm after dissection. **B,** Maximum intensity projection confocal images of day 12 PRI-ExUt and E9.5 in utero embryos stained for GATA4, SOX9 and TUJ1. Selected craniofacial, neural, cardiac and somitic structures are annotated. Scale bars, 500 μm. **C,** Quantification of properly developed embryos on day 10 (∼E7.5), day 11 (∼E8.5) and day 12 (E9.5). Dots represent percentage of embryos showing proper morphological landmarks specific to the individual developmental stage; total successful embryo numbers are indicated as n. The denominator for the efficiency calculation is derived from the starting number of E4.5 blastocysts. **D,** Comparison of embryo length in PRI-ExUt and stage-matched in utero embryos from E6.5 to E9.5. Dots represent individual embryos; bars show mean and S.D; two-sided unpaired *t*-tests; *P* values are indicated; ns = not significant. **E,** Integrated single-cell RNA-seq UMAP manifold of day 12 PRI-ExUt and E9.5 in utero embryos with annotated embryonic and extraembryonic populations. **F,** UMAP representation of day 12 PRI-ExUt embryos (red) compared to in utero E9.5 embryos (black) as individual groups or overlaid, showing high overlap between the manifolds of PRI-ExUt and in-utero embryos. n = number of cells; e = number of embryos. **G,** Schematic comparison of neural tube morphology in E8.5 in utero embryos, TF-SEMs, PSI-ExUt embryos and day 11 PRI-ExUt embryos. **H,** Proportion of embryos or embryo models displaying straight or wrinkled/undulating neural-tube morphology; *n* = 20 measured embryos per condition. **I,** Representative dorsal-view images showing straight or undulating neural-tube morphology in E8.5 in utero embryos and stage-matched TF-SEMs, PSI-ExUt embryos and PRI-ExUt embryos. Insets represent magnified dotted regions. Scale bars, 200 µm (main) and 100 µm (insets). Abbreviations: A, anterior; Al, allantois; D, dorsal; EPC, ectoplacental cone; ExEM, extraembryonic mesoderm; H, heart; InUt, in utero; Md, mandibular arch; Ms, mesencephalon; Mx, maxillary arch; NF, neural fold; NT, neural tube; OP, optic pit; OtP, otic pit; P, posterior; PN, posterior neuropore; PRI-ExUt, pre-implantation-initiated ex utero culture; Pro, prosencephalon; prog, progenitor; PSI-ExUt, post-implantation-initiated ex utero culture; Rho, rhombencephalon; S, somite; Sc, spinal cord; TF-SEM, transgene-free stem-cell-derived embryo model; V, ventral; YS, yolk sac.

Proper embryo morphology was accompanied by extensive and adequate cell-type diversification, as measured by integrated single-cell RNA-seq analysis of day 12 PRI-ExUt embryos and E9.5 in utero controls, with minimal variation in cell-type proportions between cultured embryos and in utero conditions (**Fig. 4E-F, Extended Data Fig. 13A**). All clusters identified in PRI-ExUt embryos overlapped extensively with transcriptional states found in control embryos and showed comparable representation of many embryonic and extraembryonic populations (**Fig. 4E-F, Extended Data Fig. 13**). Annotated cell lineages included embryonic and extraembryonic tissues, including somites, endotome (vascular endothelium derived from somites), surface ectoderm, hindgut, cardiomyocytes, neural tube, neural crest, regionalized brain progenitors, neuromesodermal progenitors, endothelium, hematoendothelial progenitors, myeloid-erythroid populations, chorion and amnion/allantois-associated lineages (**Fig. 4E, Extended Data Fig. 13A-C**). Further analysis of gene expression correlations between E9.5 in utero embryos and ex utero samples demonstrated that E9.5 embryos exhibited the highest correlations across developmental stages, ranging from 0.939 to 0.984 in all cell types (**Extended Data Fig. 10C**).

Finally, to determine whether PRI-ExUt embryos progress through developmental time in a coordinated manner, we integrated single-cell datasets across E7.5, E8.5 and E9.5 matched embryos and examined lineage trajectories. Epiblast and nascent mesoderm populations in E7.5-equivalent embryos gave rise to neural, mesodermal and neuromesodermal states at E8.5, followed by more differentiated somite, endotome, surface ectoderm, neural tube, dorsal midbrain, hindbrain floorplate, migratory neural crest and cardiac populations at E9.5 (**Extended Data Fig. 13D-E**). Importantly, PRI-ExUt cells followed the same temporal progression as in utero cells across these developmental intervals, indicating preservation of developmental timing and lineage maturation rather than asynchronous or aberrant differentiation. Analysis of trophoblast-derived populations revealed progressive remodeling of the extraembryonic compartment between E7.5 and E9.5, mimicking trajectories described for intrauterine development^73^ (**Extended Data Fig. 13F-G**). At E7.5, PRI-ExUt and in utero embryos contained a diverse repertoire of chorion progenitors, intermediate chorion and trophoblast giant cell (TGC)-derived lineages (**Extended Data Fig. 13B,G**). This diversity progressively decreased at E8.5 and E9.5 in both PRI-ExUt and their in utero counterparts, with intermediate chorion progressively becoming the predominant population, indicating that trophoblast-derived lineages can undergo substantial differentiation and maturation ex utero in the absence of direct maternal tissue interactions. TGC progenitors were retained at similar proportions at E9.5, whereas more differentiated TGC cell types detected in utero were reduced or absent in PRI-ExUt embryos (**Extended Data Fig. 13B,G**). A similar reduction in TGC diversity was observed in PSI-ExUt embryos, suggesting that this difference is associated with ex utero development rather than specifically with bypassing in vitro implantation (**Extended Data Fig. 13B,G**). Collectively, these analyses demonstrate that mouse embryos grown entirely ex utero from zygotes can establish embryonic and extraembryonic cell states that highly resemble their equivalent in utero counterparts across gastrulation, neurulation and organogenesis until mid-gestation.

Mouse SEMs and TF-SEMs are assembled exclusively from naive pluripotent stem cell-derived Epi-, PrE- and TE-like lineages^25,72^, bypassing the blastocyst-like stage altogether. Following co-aggregation, these lineages undergo appropriate sorting and post-implantation egg-cylinder morphogenesis entirely in suspension, generating E5.5-like egg-cylinder structures that can progress up to E8.5. However, SEMs uniformly display a characteristic buckled or undulating neural tube (**Fig. 4G**)^25,72^. In utero embryos, by contrast, show a relatively straight neural tube along the rostro-caudal axis. The recurrence of this malformation in SEMs, but not in stage-matched natural embryos, raises three possible explanations: (a) it results from the use of the roller culture system, whose rotational forces might deform the neural tube as it elongates at around E8.0-E8.5; (b) it is restricted to stem-cell-derived embryo models, in which issues related to suboptimal aggregation may leave the contributing lineages imbalanced; or (c) it reflects aberrant early post-implantation morphogenesis in embryos that never implant in the uterus, irrespective of whether they are stem-cell-derived. The first explanation can be excluded directly, as stage-matched E8.5 PSI-ExUt embryos, which are also grown in roller culture after dissection at E7.5, show no such defect. To distinguish between the remaining two, we reasoned that E8.5 PRI-ExUt embryos would be informative (**Fig. 4G**), since they implant in vitro yet are derived from natural zygotes rather than from in vitro grown stem cells. Thorough analysis revealed that PRI-ExUt embryos also frequently displayed the same buckled neural tube seen in SEMs (**Fig. 4I**), indicating that the phenotype is not a property of stem-cell-derived models as such, but rather of the one feature the two systems share: progression through post-implantation morphogenesis without implantation in the uterus. Notably, this malformation coincided with a reduced capacity of both PRI-ExUt embryos and SEMs to develop beyond E8.5-E8.75; although a small fraction of PRI-ExUt embryos subsequently reached E9.5, but none progressed further (**Fig. 4C**). Together, these comparisons across embryos and embryo models identify mechanical deformation of the neural tube as a recurrent feature of embryos that traverse the peri-implantation transition in vitro rather than in vivo, and suggest that events during this window can shape morphogenesis at later developmental stages.

We subsequently examined the trophoblast-derived Ectoplacental Cone (EPC), which leads maternal implantation and invasion in early post-implantation E5.5-E6.5 embryos across all four experimental systems (in utero controls, PSI-ExUt, PRI-ExUt and SEMs). Remarkably, in 100% of TF-SEM/SEMs and 57.1% of the analyzed PRI-ExUt embryos, where the EPC neither implants into nor invades the maternal uterus, this region was aberrantly covered by primitive endoderm-derived visceral endoderm, which went on to surround the entire embryo, EPC included, at stages right after implantation (**Extended Data Fig. 14**). In contrast, in utero controls and PSI-ExUt embryos at E6.5 displayed a more developed EPC, which was neither covered nor constrained by visceral endoderm (VE) (**Extended Data Fig. 14C**). These observations link the absence of in vivo implantation and uterine invasion to aberrant VE morphogenesis. Moreover, there is a negative correlation between the frequency of the latter inadequate VE morphogenesis, which is much higher in SEMs (**Extended Data Fig. 14B,D**), and the frequency of reaching E8.5 and E9.5 which is much higher in PRI-ExUT (**Fig. 3E**) compared to SEMs^25^ . How this subtle aberrant extraembryonic tissue organization observed at ∼E6.5 could be related to abnormal morphogenesis or the buckling of the elongating neural tube in E8.5 SEMs and PRI-ExUt embryos remains to be defined. Establishing such relationship will be important for future mechanistic studies, and may point to ways of extending the developmental potential of both systems beyond E8.5-E9.5.

### Enhanced blastoids complemented with PrESCs reach gastrulation in PRI-ExUt platform

Developing ex utero platform for growing natural mouse E6.5 egg-cylinder embryos to post-gastrulating E8.5-E12.5 was an instrumental step and reference platform whose usage enabled optimizing and obtaining up to E8.5 mouse SEMs solely from stem cells^3,25,57,58,61^. Similarly, we reasoned that the ability of PRI-ExUt to sustain the development of natural mouse blastocysts through implantation, gastrulation and organogenesis could provide a stringent functional reference for evaluating mouse blastoids in vitro developmental under the same culture conditions. Specifically, having established that PRI-ExUt supports continuous development of natural mouse embryos from the one-cell stage to approximately E9.5, we next asked whether the same culture environment could indicate or extend the developmental potential of established mouse stem-cell-derived blastoids systems^41^ . This distinction is important, since current state of the art in which failure of validated mouse blastoids to develop in regimens that do not adequately support the equivalent transitions in natural blastocysts, cannot readily be attributed to inherent limitations of the stem-cell-derived blastoids, rather than to the culture conditions themselves which haven’t been shown to support natural blastocyst growth towards post-implantation stages.

Therefore, we used the PRI-ExUt platform for assaying developmental potential of mouse blastoids that were generated and validated by some of most recently generated and commonly used protocols. First, we tested an established blastoid strategy based on aggregation of mouse embryonic stem cells (ESCs) and trophoblast stem cells (referred to herein as Established Blastoid Protocol #1^41^) (**Fig. 5A**). The resulting blastocyst-like structures displayed blastocyst-like morphology and contained spatially segregated OCT4^+^ epiblast and CDX2^+^ TE compartments, comparable to E3.5-E4.5 natural blastocysts (**Fig. 5B-D**), as originally reported^41^, and PrE contribution was rather more limited, with many structures containing few or no GATA4^+^ cells (**Fig. 5D**). After this validation and excluding the possibility that some of these blastocyst-like structure have similar morphology to blastocyst but have aberrant SOX17+ PrE distribution (**Extended Data Fig. 15 vs. Fig. 5D)**) which can be obtained when applying Post-implantation SEM conditions during blastoid formation stages, we applied PRI-ExUt on the latter validated blastoids. Although the latter validated blastoids attached under PRI-ExUt implantation conditions, they showed no progression toward organized egg-cylinder morphologies (**Fig. 5E**). We utilized and validated another recently established protocol to generate mouse blastoids, however such blastoids also failed to demonstrate progression to bona-fide egg-cylinder post-implantation stage upon applying PRI-ExUt strategy, regardless of donor ESC line tested (**Extended Data Fig. 16**).

**Figure 5.**
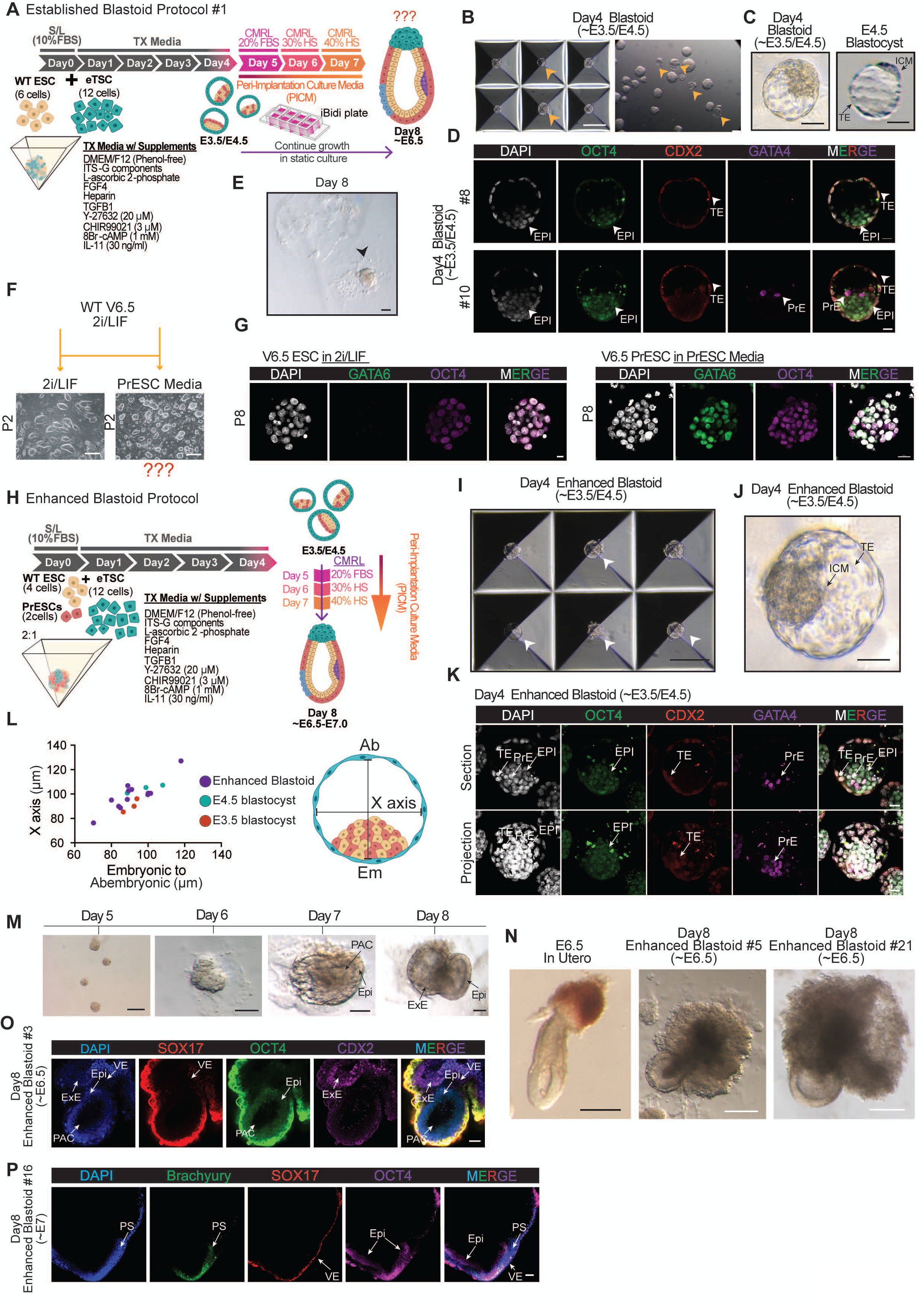
PRI-ExUt and enhanced primitive endoderm contribution support post-implantation development of blastoids towards mid-gastrulation. **A,** Schematic overview of the ESC-TSC blastoid protocol, which we termed Established Blastoid Protocol #1, followed by transfer of day 4 blastoids to PRI-ExUt implantation conditions. **B,** Representative day 4 blastoids generated using the ESC-TSC protocol. Yellow arrows point to properly cavitated structures. Scale bars, 200 μm and 50 μm. **C,** Representative brightfield images day 4 blastoids compared with E4.5 natural blastocyst. Scale bars, 50 μm. **D,** Whole-mount confocal images of day 4 blastoids immunostained for OCT4, CDX2 and GATA4, marking epiblast-like (EPI), trophectoderm-like (TE) and primitive-endoderm-like (PrE) compartments, respectively. Scale bars, 20 μm. **E,** Representative stereomicroscope images of failed day 8 structures obtained from ESC-TSC blastoids cultured under PRI-ExUt conditions. Black arrow points to a typical failed structure observed. Scale bar, 200 μm. **F,** Schematic depiction of strategy for establishing PrESCs from 2i/LIF mouse naive ESCs with the representative brightfield images of the cells in culture after 2 passages in the indicated media. Scale bars, 200 μm. **G,** Representative immunofluorescence for GATA6 and OCT4 in ESCs maintained in 2i/LIF or their derived PrESC cultures induced and maintained in PrESC medium. P indicates passage number in the respective culture media. Scale bar for 2i/LIF, 10 μm; scale bar for PrESCs, 20 μm. **H,** Schematic overview of the enhanced blastoid protocol incorporating naive ESCs and PrESC-like cells at a 2:1 ratio before addition of TSCs and subsequent growth in PRI-ExUt platform. **I, J,** Representative brightfield images of day 4 blastoids generated using the enhanced blastoid protocol. White arrows point to properly cavitated structures. Scale bars, 200 µm (**I**), 50 µm (**J**). **K,** Middle-section and maximum intensity projection views of day 4 PrE-enhanced blastoids stained for OCT4, CDX2 and GATA4, marking epiblast-like (EPI), trophectoderm-like (TE) and primitive-endoderm-like (PrE) compartments, respectively. Scale bars, 20 μm. **L**, Morphometric comparison of enhanced blastoids and natural E3.5 and E4.5 blastocyst dimensions along the x and embryonic-to-abembryonic axes. Dots represent individual embryos measured. **M,** Representative stereomicroscope images of PrESC-enhanced blastoid, (generated with enhanced blastoid protocol), progression from culture days 5 to 8 following transfer to PRI-ExUt conditions. Scale bars, 200 µm (day 5), 100 µm (day 6), and 50 µm (day 7 and day 8). **N,** Bright-field images of day 8 enhanced blastoid-derived structures compared with an E6.5 in utero embryo. Scale bars, 100 µm. **O,** Representative whole-mount immunofluorescence of a day 8 enhanced blastoid-derived egg-cylinder structure stained for CDX2 (ExE), SOX17 (VE) and OCT4 (Epi). Scale bars, 50 µm. **P,** Representative confocal image of a day 8 enhanced blastoid-derived structure showing BRACHYURY^+^ PS extending distally, together with SOX17 (VE) and OCT4 (Epi), developmentally corresponding to ∼E7. Scale bars, 50 µm. Abbreviations: Ab, abembryonic pole; Em, embryonic pole; Epi, epiblast; ESC, embryonic stem cell; eTSCs, embryo-derived trophoblast stem cell; ExE, extraembryonic ectoderm; FBS, fetal bovine serum; HS, human serum; ICM, inner cell mass; PAC, pro-amniotic cavity; PrE, primitive endoderm; PrESC, primitive endoderm stem cell; PRI-ExUt, pre-implantation-initiated ex utero culture; PS, primitive streak; TE, trophectoderm; VE, visceral endoderm.

We next moved to utilize the PRI-ExUt system to generate enhanced mouse blastoids with potential to develop into post-implantation embryo-like structures. Given that nearly all mouse and human blastoid generating protocols noted limited PrE abundance in generated structures^41,44,74^, which was also noted herein (Fig. 5D), we hypothesized that the limited growth of currently available mouse blastoids may be, at least partially, due to inefficient PrE specification that could impair post-implantation morphogenesis. Considering these results, we decided to pursue optimizations to boost the PrE abundance in the ICM compartment in the blastoids generated with the Established Blastoid Protocol #1. Notably, a recent study established novel growth conditions to derive **Pr**imitive **E**ndoderm **S**tem **C**ells (PrESC) from pre-implantation mouse embryos, that are double positive for OCT4+ (pluripotency) and GATA6+ (PrE) markers and can be indefinitely expanded in vitro in such conditions^75^ while maintaining ability to reconstitute the entire PrE/yolk-sac compartment in vivo. Recent work from our group indicated other unique conditions (termed alternative condition (AC) medium) that enhance the plasticity of in vitro expanded mouse ESCs towards extraembryonic lineages and can induce OCT4+ cells co-expressing GATA6, however these cells could not be stabilized as OCT4+/GATA6+ but were rather oscillating between the OCT4+GATA6-ESC and OCT4+/GATA6+ PrE-primed state^72^ . Given the latter two studies, we were motivated to generate a stable OCT4+GATA6+ PrESC population from naive in vitro grown ESCs, which has not been described before. To monitor extraembryonic lineage induction, we generated a triple-reporter ESC line carrying nuclear fluorescent reporters for OCT4, GATA6 and CDX2, labelling epiblast, PrE and TE cells, respectively (**Extended Data Fig. 17A**). Naive ESC growth conditions were switched from conventional 2i/LIF directly into the PrESC culture conditions established by Ohinata et al. for blastocyst-derived primitive endoderm stem cells (**Fig. 5F**)^75^. Flow cytometry analysis confirmed that ESCs maintained in 2i/LIF were predominantly OCT4+ with negligible activation of the GATA6 or CDX2 reporters, whereas transfer to PrESC conditions generated a GATA6^+^ population (∼18%) that retained OCT4 expression (**Extended Data Fig. 17B**). scRNA-seq analysis of mouse ESCs before and after transfer into PrESC conditions confirmed the induction of PrESC sub-population in vitro that co-expresses OCT4 and PrE markers (GATA4, GATA6, SOX17 and PDGFRa) (**Extended Data Figure 18**), alongside a retained refractory ESC population and the formation of an OCT4+ population that is distinct from ESCs and lacks PrE markers, which we term poised ESCs (**Extended Data Figure 18**),. Next, we assessed the fidelity of the in vitro generated PrESC cluster, which confirmed its high transcriptional similarity to previously described embryo derived PrESCs (**Extended Data Figure 19**). Upon subclone selection or sorting for the PDGFRa surface marker, followed by expansion in PrESC medium, we obtained cell cultures that uniformly expressed OCT4+GATA6+ in addition to other PrE markers, consistent with acquisition and maintenance of a PrESC-like identity (**Fig. 5G, Extended Data Fig. 20A**).

Next, the in vitro derived and expanded PrESCs were subsequently incorporated into blastoid assembly together with naive ESCs and TSCs as part of a modified protocol which we term the enhanced blastoid protocol (**Fig. 5H**). On the day of aggregation, ESCs and PrESCs were mixed at a 2:1 ratio in Serum/LIF media. The next day, the eTSCs cultured under TXV conditions were added on top of the already aggregated mix of ESCs and PrESCs in the TX media (supplemented with FGF4, heparin, TGFB1, 8Br-cAMP, IL-11, CHIR99021 and Y-27632). The resulting blastoids exhibited proper blastocyst morphology **(Fig. 5I-J)**, similar dimensions to mouse blastocysts **(Fig. 5L)**, and contained organized OCT4^+^/NANOG^+^ epiblast, surrounding CDX2^+^ TE layer, and a prominent GATA4^+^/GATA6^+^ PrE compartment (**Fig. 5K, Extended Data Fig. 20B-C**). Following transfer to PRI-ExUt conditions, PrESC-containing blastoids attached and showed improved post-implantation morphogenesis, generating egg-cylinder-like structures containing a cup-shaped epiblast expressing OCT4, a pro-amniotic cavity, a well-defined CDX2^+^ ExE and a SOX17^+^ visceral endoderm surrounding the egg-cylinder (**Fig. 5M-O, Extended Data Fig. 20D-F**). By culture day 8, structures reached morphologies resembling approximately E6.5-E7.0 embryos and initiated and progressed through gastrulation, as indicated by the emergence of BRACHYURY^+^ primitive-streak cells migrating distally within the embryonic compartment (**Fig. 5P**). Thus, improving PrE contribution substantially extended the post-implantation potential of mouse blastoids. Nevertheless, this progression was not sustained after culture day 9, when these structures showed progressive reduction or loss of the embryonic compartment while extraembryonic yolk sac-like tissues continued to expand (**Extended Data Fig. 20G-H**). Thus, the PRI-ExUt platform allowed evaluation and improvement of the in vitro developmental potential of mouse blastoids, by incorporating in vitro isolated PrESCs, which enabled such enhanced blastoid-derived structures to undergo implantation and initiate gastrulation without the use of any transgene or exogenous transcription factor over-expression. However, it was insufficient to support the sustained gastrulation-to-organogenesis progression observed in zygote-derived PRI-ExUt embryos that can reach E8.5-E9.5 stages.

## Discussion

Here, we establish and benchmark a continuous ex utero culture system (PRI-ExUt) that supports mouse development from the totipotent zygote (E0.5) to the E9.5 stage, representing nearly half of total gestation in mice. The system comprises a pre-implantation culture phase followed by sequential serum-based culture media optimized for sustaining implantation-like developmental progression and generating post-implantation-stage embryos, combining static culture, shaking-culture and optimized electronically controlled ex utero roller culture systems. Integrated morphological characterization, whole-mount immunostaining, and single-cell transcriptomics demonstrated that isolated zygotes can properly execute implantation, gastrulation, somitogenesis, axial turning and neural tube closure outside the uterus, while establishing both embryonic and extraembryonic lineages. Notably, progression across implantation required neither specialized three-dimensional implantation culture, exogenous extracellular matrices nor uterine-cell co-culture. Embryos instead attached to a two-dimensional surface and continued to develop in a matrix-free environment, one that permits direct observation and full experimental access. The persistence and progressive differentiation of trophoblast-derived lineages in PRI-ExUt embryos further indicate that substantial aspects of extraembryonic development can proceed independently of direct interaction with maternal tissues even when starting from pre-implantation stages and in utero implantation is bypassed. Although absolute efficiency levels were relatively low and declined with developmental stage progression, this platform establishes that a mammalian embryo, in this case the mouse, can autonomously execute the continuous developmental sequence from fertilization to early organogenesis, with morphological and molecular fidelity closely approaching that of embryos grown in utero, and to a substantially greater extent than described in pioneering studies using human blastocysts grown up to day 14^12,13,33^, or to those of non-human primates (NHPs) carried up to days 21–23^63,64^, when drawing a cross-species developmental stage analogy.

By granting direct experimental access to morphogenetic events occurring in natural embryos, whose study has been constrained by dependence on maternal tissues or by the discontinuity of existing in vitro systems, PRI-ExUt stands to expand the ways in which the first half of mouse gestation can be investigated. Perturbation, live imaging and dynamic morphogenetic analysis become possible across developmental transitions that previously had to be examined in separate systems. The platform should likewise expand the mechanistic study of how metabolic, mechanical, transcriptional and signalling programs are coordinated continuously, from the fertilized zygote to the organ-bearing embryo at mid-gestation.

Indeed, we demonstrate relevant examples of how the PRI-ExUt platform established herein can achieve some of these ambitions. PRI-ExUt allowed us to evaluate the developmental potential of previously established protocols for generating bona fide blastoids^41,55^, and to test how their development could be optimized in vitro. Importantly, when natural blastocysts have not been shown to progress adequately under the culture conditions used for blastoid experiments, it remains difficult to distinguish intrinsic deficiencies in the blastoid model from limitations imposed by the culture regimen itself. Establishing the conditions that support faithful development of natural blastocysts therefore provides an essential reference for interpreting and optimizing mouse blastoid models, and constitutes a relevant principle for future studies of human blastoids as well. Existing and optimized blastoids provide valuable models of blastocyst organization, implantation and early post-implantation morphogenesis up to mid-gastrulation, particularly when unique PrE cells induced from isogenic naive PSCs are incorporated into the ICM compartment. However, these enhanced egg-cylinder-stage blastoid-derived structures did not reproducibly progress through gastrulation and organogenesis, suggesting that current blastoid systems capture some, but not all, of the developmental properties required for sustained post-gastrulation mammalian embryogenesis. PRI-ExUt therefore provides both a culture system for natural embryos and a benchmark platform for stem-cell-derived pre-implantation embryo models, providing a calibrated system that can be used to identify improved cell lineage compositions or culture conditions that can extend blastoid developmental capacity.

A particular feature shared by PRI-ExUt embryos and SEMs was the emergence of a wrinkled/undulating neural tube morphology around E8.5^25,60,61,72,76^, a phenotype largely absent from natural and PSI-ExUt embryos^3,5^. The occurrence of this phenotype in natural embryos implanted in vitro suggests that this abnormality is not intrinsic to stem-cell-derived models but may instead originate from altered morphogenesis and axial elongation during egg-cylinder development in the absence of normal embryo-uterine interactions during implantation stages. Indeed, previous work has shown that mechanical forces exerted by maternal tissues contribute to egg-cylinder elongation and anterior-posterior axis formation^77^, indicating that robust early post-implantation morphogenesis may depend on uterine cues. Notably, neural tube undulation coincides with a developmental bottleneck around E8.5 in both PRI-ExUt embryos and mouse SEMs, raising the possibility that altered mechanical organization established during peri-implantation development becomes increasingly relevant as axial growth and tissue complexity increase. Uncovering the nature of these interactions, and determining whether artificially providing them could prevent neural tube undulation and extend mouse pre-implantation natural embryo or post-implantation SEM development beyond the E8.5-E9.5 “barrier” will be an important direction for future optimization.

We found that in vitro implantation-like step in our set-up is accompanied by a transient developmental slowdown. This in vitro-induced slowdown might suggest that the uterus normally provides physical, metabolic or other signaling inputs controlling developmental timing that are, so far, not fully understood and reproduced in vitro. Although embryos formed blastocysts on schedule and attached under implantation-like conditions, they required approximately three additional days to transition from E4.5 late blastocyst to the E5.5 egg-cylinder morphology. During this period, embryos remained transcriptionally delayed relative to in utero development and occupied an intermediate state between E4.5 and E5.5 that does not seem to represent the known canonical diapause state^69–71,78–81^, but rather exhibited a transient deceleration of peri-implantation development. A subset of embryos reached the E5.5 egg-cylinder stage (achieved on day 8 of the protocol) and subsequently resumed morphogenesis and progressed through gastrulation and organogenesis up to E9.5 (achieved on day 12 of the protocol) within 4 days (**Extended Data Fig. 1**), following kinetics largely similar to those of in utero development. Thus, in vitro implantation uncouples culture time from developmental time, while revealing that peri-implantation embryos can recover from a substantial temporal developmental slowdown/delay and adequately re-engage developmental progression. This developmental delay, and the ability to catch up within certain time windows, has implications for how extended human embryo culture should be discussed and regulated. The 14-day rule has provided an essential ethical and regulatory boundary for human embryo research, but it is based on the assumption that culture time and developmental time remain closely aligned. Our findings show that mammalian embryos may spend additional time in culture before reaching the equivalent chronological developmental stage, suggesting that future discussions of extended embryo culture should also consider developmental stage, morphology and molecular state, rather than culture time alone, when setting biologically meaningful limits or added regulations.

This study shows that the advanced mouse embryos cultured ex utero from the zygote closely resemble their in utero counterparts at both morphological and molecular levels, although the two are not identical. Our aim, however, is to establish an ex utero embryo model system based on pre-implantation embryos, as early as zygotes, as starting material, which reproducibly captures the major developmental milestones and forms the key embryonic structures in recognizable and developmentally appropriate configurations, even if it does not yet fully reproduce in utero development. One limitation of the current platform is its relatively low efficiency, particularly at the latest developmental stages, and this will need to improve before the system can support certain experimental applications. Several observations and recent developments in other related systems indicate that such improvement is likely feasible. Protocols for generating post-gastrulation mouse SEMs have progressively improved in both fidelity and efficiency^25,72^ and natural mouse embryos explanted at E6.5 can now be cultured to E12.5, rather than to E11 only as initially demonstrated, by using an improved roller culture setup and culture conditions at later stages^5^. These advances suggest that further optimization, together with a mechanistic understanding of how embryo self-organization proceeds ex utero, could substantially improve both culture efficiency and reproducibility from pre-implantation stages. Developing fully defined, serum-free conditions is another important technical priority that may improve yield and reduce variability while potentially alleviating the approximately three-day developmental delay observed during the E4.5-to-E5.5 transition. Simpler culture platforms and growth conditions are also likely to emerge over time, and may ultimately match or exceed the results reported here. Finally, determining whether embryos cultured continuously from the zygote can progress past E9.5 toward late organogenesis and fetal stages (E11.5-E12.5), as has been achieved when ex utero embryogenesis is initiated from mouse post-implantation embryos^5^, would further expand the experimental opportunities this platform offers.

Our success in establishing and benchmarking PRI-ExUt from mouse zygotes to E9.5, broadly corresponding to human Carnegie stages 12-13 (∼26-32 days post fertilization), informs ongoing international ethical discussions and legislative efforts to revisit the limits placed on human embryo culture beyond day 14. Most consequential would be to establish whether the approach devised here can be adapted to grow human pre-implantation embryos through gastrulation and organogenesis, which may offer insight into windows of early human development that have so far been inaccessible. This is particularly relevant since, to date, human and non-human primate pre-implantation embryos cultured in vitro recapitulate only rudimentary features of post-implantation differentiation and remain structurally far from their in utero counterparts. Before the present study, this limitation raised the question of whether it was justified to apply continuous pre-implantation-to-organogenesis culture to scarce and ethically sensitive human embryonic material before faithful development through organogenesis had been achieved in any mammalian species. Such research should be conducted only after serious and open ethical discussion, and following the relevant legal and ethical approvals^40,82,83^.

## Extended Data Figure Legends

**Extended Data Figure 1.**
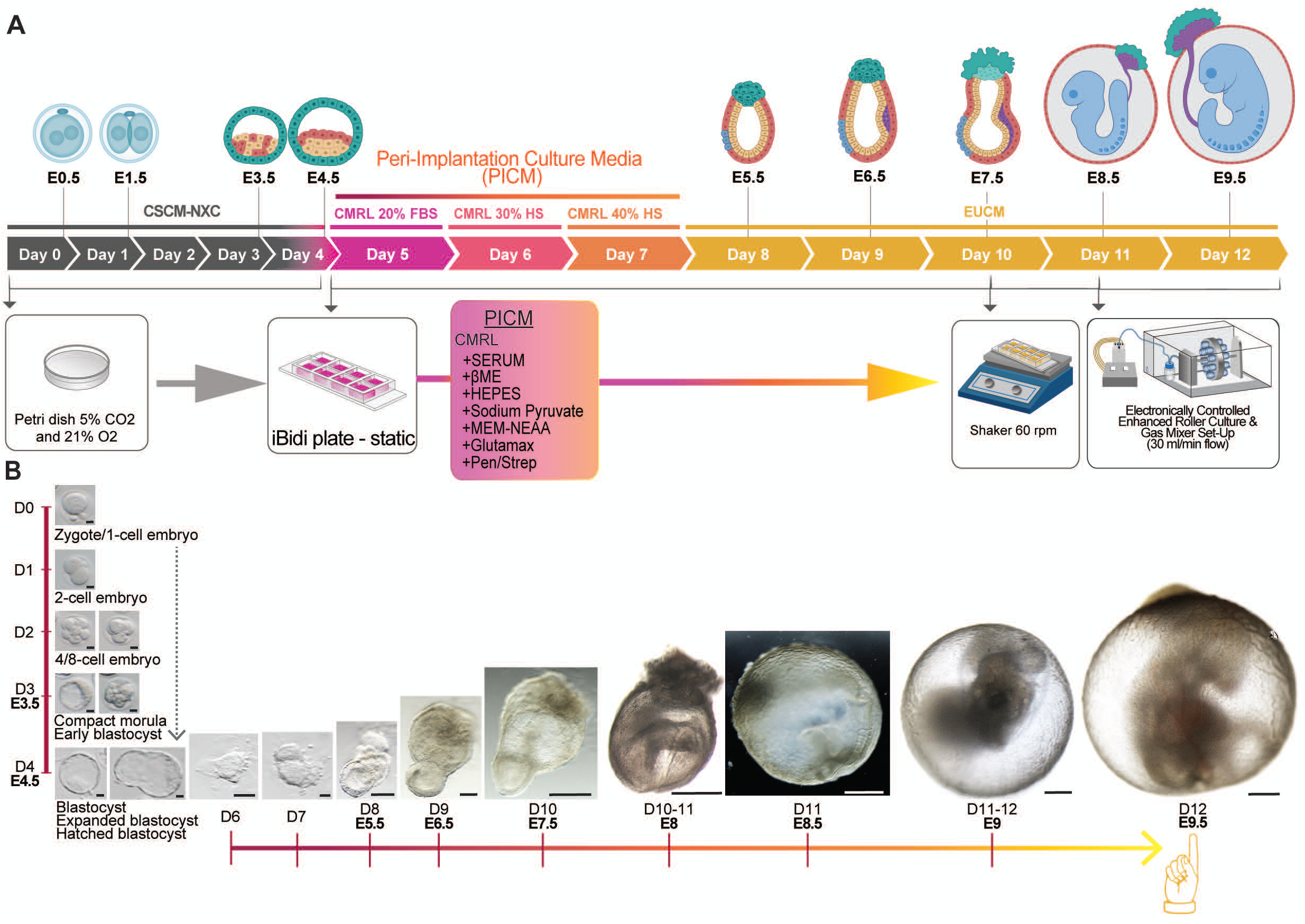
PRe-implantation-Initiated Ex Utero culture (PRI-ExUt) is a continuous ex utero culture platform from zygote to E9.5 mid-gestation in mice. **A,** Schematic depiction of the finalized complete **PRe-implantation-Initiated Ex Utero culture (PRI-ExUt)** platform and protocol to grow mouse E0.5 zygotes to E9.5-stage embryos (protocol day 12), indicating sequential use of media and static, shaker and roller-culture phases and typical morphology and developmental stage observed at each protocol day. **B,** Representative morphological progression from the one-cell mouse embryo through blastocyst, egg-cylinder, gastrulation and organogenesis stages during PRI-ExUt culture. Scale bars, 25 µm E0.5-E4.5; 50 µm day 5-8; 100 µm day 9; 200 µm day 10; 500 µm day 10-11 and 11; 1 mm day 11-12 and 12. Abbreviations: CMRL, CMRL 1066 medium; CSCM-NXC, Continuous Single Culture-NX Complete pre-implantation media; EUCM, ex utero culture medium; FBS, fetal bovine serum; HS, human serum.

**Extended Data Figure 2.**
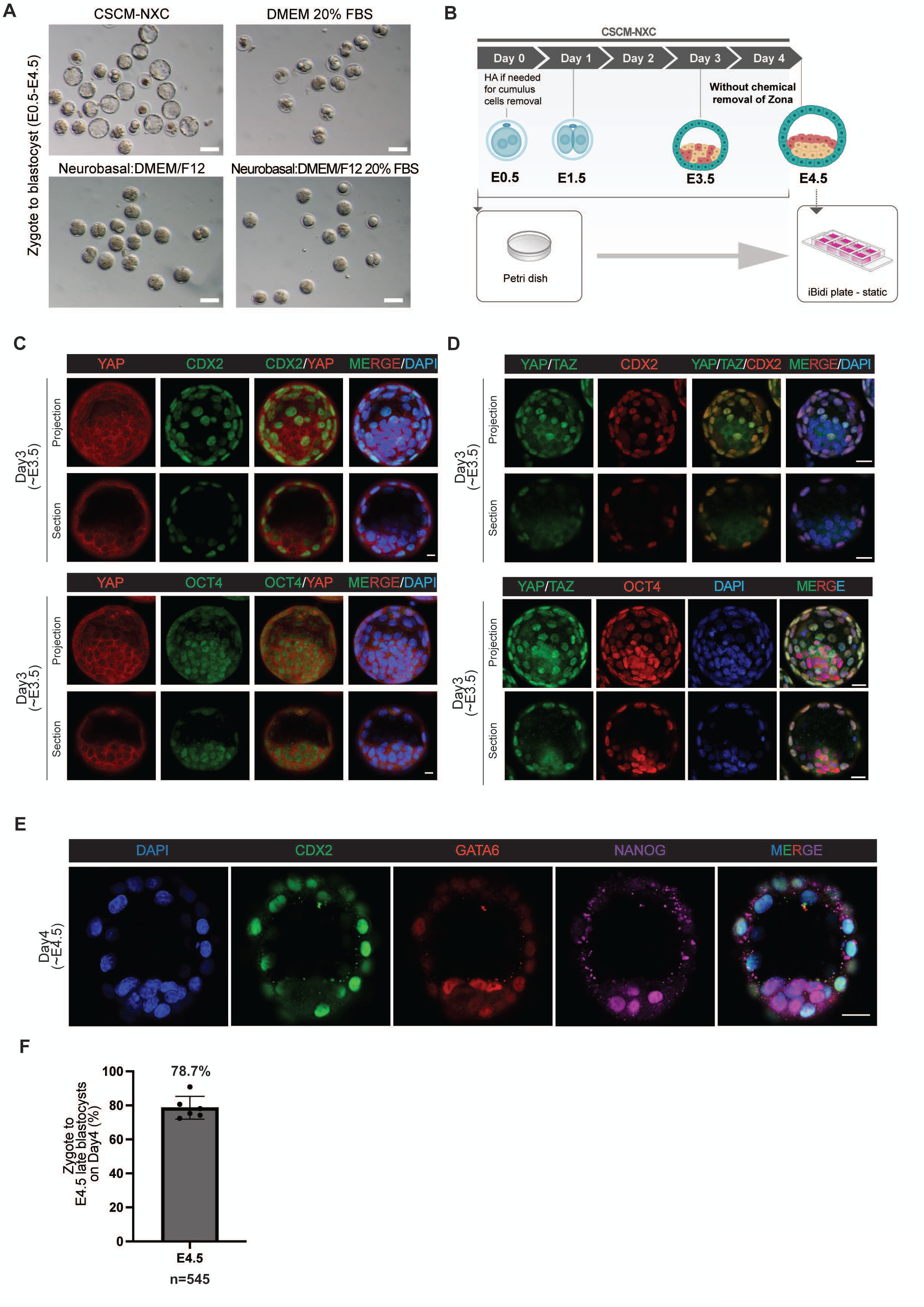
Lineage characterization of mouse blastocysts generated after up to 4 days in CSCM-NXC media from in vitro cultured zygotes. **A,** Representative bright field images comparing culture success of different basal media during in vitro pre-implantation development to E4.5. Only CSCM-NXC was found conducive for this. Scale bars, 100 μm. **B,** Schematic for the pre-implantation embryo culture protocol adopted herein. Naturally hatched blastocysts (without chemical removal of the zona pellucida) were chosen for continued growth. Abbreviations: HA, hyaluronidase. **C,** Representative immunostainings of E3.5 blastocysts, obtained from zygotes grown 3 days in CSCM-NXC, stained for YAP and OCT4 or YAP and CDX2, showing the nuclear YAP localization in TE lineage exclusively. Scale bars, 10 μm. **D**, Representative immunostainings of E3.5 blastocysts stained for YAP/TAZ and OCT4 or YAP/TAZ and CDX2 showing proper TE allocation in cultured blastocyst. Scale bars, 20 μm. **E,** Representative immunostainings of E4.5 blastocysts obtained from cultured zygotes for 4 days in CSCM-NXC, stained for GATA6 (PrE), CDX2 (TE) and NANOG (Epi). Scale bars, 20 μm. **F,** Quantification of properly developed embryos on day 4 (∼E4.5). Dots represent percentage of embryos showing proper cavitation specific to the E4.5 stage; total successful embryo numbers are indicated as n.

**Extended Data Figure 3.**
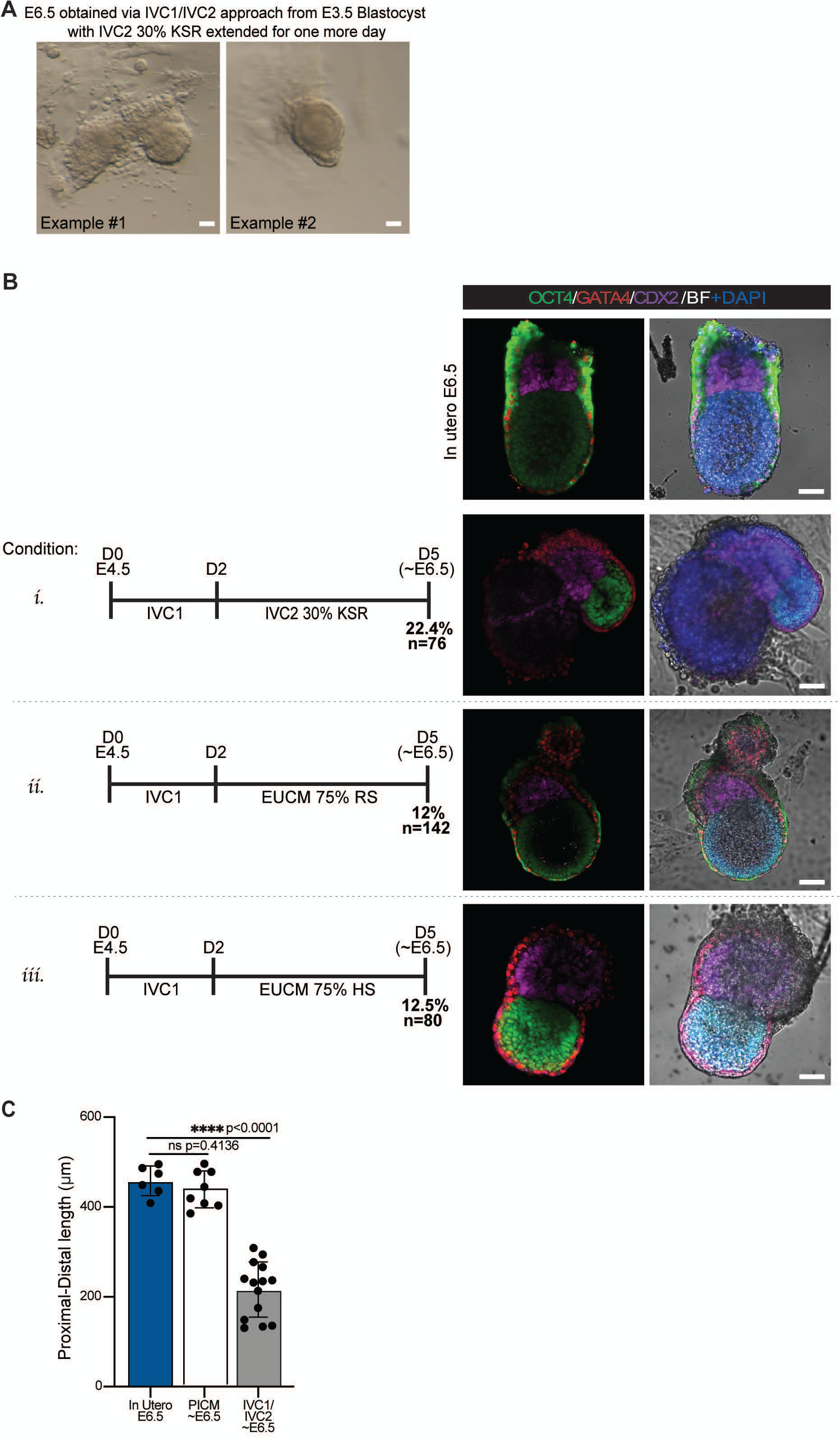
Previously established IVC conditions do not sustain development beyond early egg-cylinder stages. **A,** Representative morphology of egg-cylinders obtained from previously published IVC1/IVC2 strategy (condition i in **B**) following one more day of culture with IVC-20% KSR from day 5 to day 6. Further growth or maturation of egg-cylinders was not observed on day 6 following this step. **B,** Comparison of IVC culture regimens and efficiency percentages of properly developed embryos when starting from E4.5 blastocysts (defined here as D0). n = number of embryos evaluated, and whole-mount immunofluorescence for OCT4 (Epi), GATA4 (VE) and CDX2 (ExE) in embryos generated under IVC followed by IVC2-30% KSR or EUCM with different serum supplementations. Scale bars, 100 μm. **C,** Proximal-Distal length of E4.5 derived PICM E6.5, IVC1/IVC2 E6.5 embryos (Bedzhov et al. Cell 2014), compared to in utero E6.5 embryos. Dots represent individual embryos; bars show mean and S.D; two-sided unpaired *t*-tests; *P* values are indicated; ns = not significant. Scale bars, 50 μm. Abbreviations: EUCM, ex utero culture medium; FBS, fetal bovine serum; HS, human serum; IVC, in vitro culture; KSR, Knockout Serum Replacement; PRI-ExUt, pre-implantation-initiated ex utero culture; RS, rat serum.

**Extended Data Figure 4.**
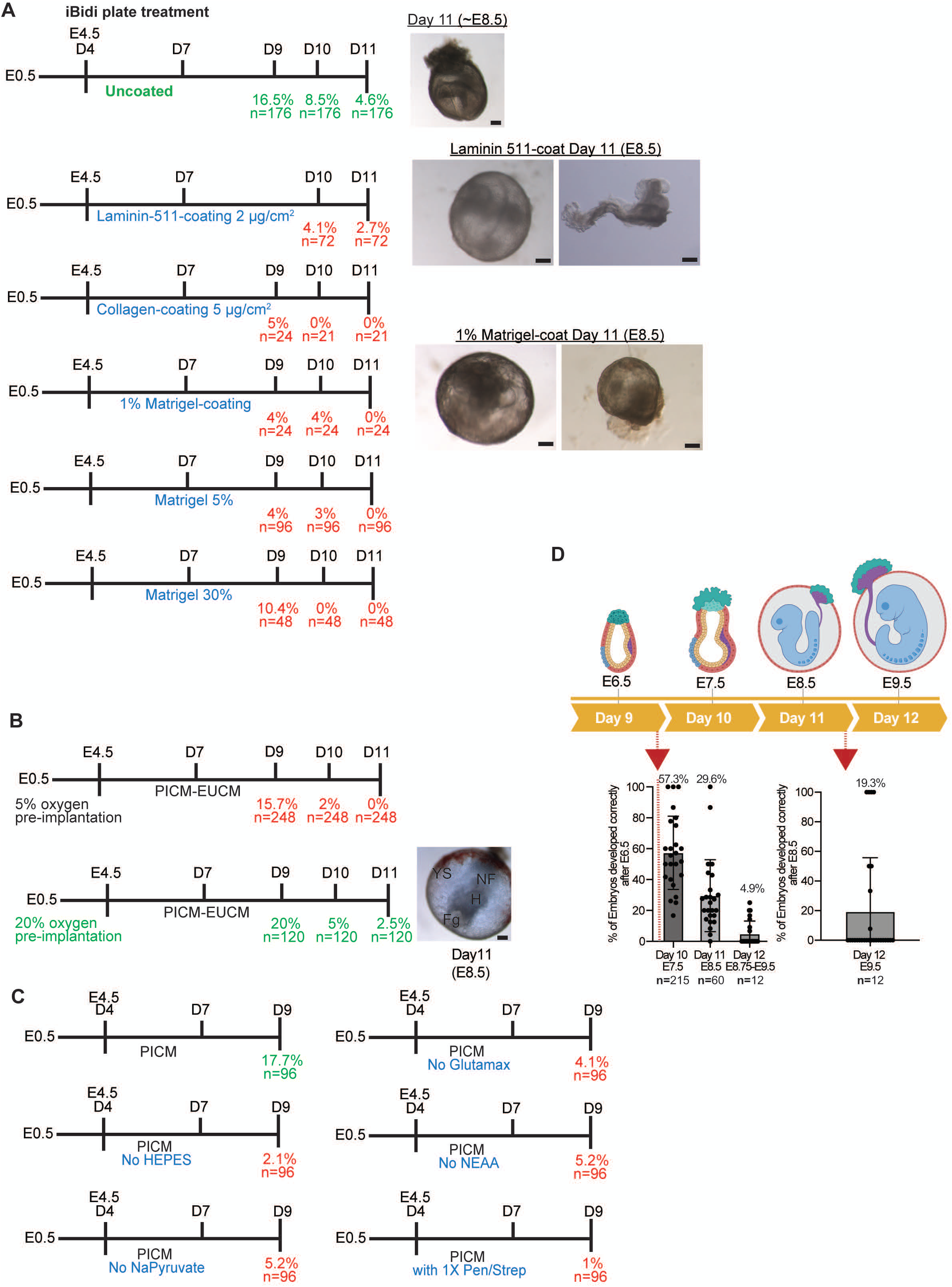
Optimization of pre- and peri-implantation PRI-ExUt culture conditions. **A**, Comparison of extracellular-matrix coatings and 3D-Matrigel embedding concentrations during in vitro implantation culture until day 11. **B,** Comparison of 5% and 20% O₂ during pre-implantation culture and subsequent PRI-ExUt development. **C,** Efficiencies for the indicated culture condition relative to optimized control conditions. Schematic representation comparing culture optimizations for PRI-ExUt relating to evaluating effect of supplementing PICM during IVI-like stage (day4-7) with the indicated supplements. Developmental outcome and efficiencies at day 9 (∼E6.5) are indicated. In all conditions, EUCM was used after Day 7 (**Extended Data** Fig. 1). n = number of embryos evaluated **D,** The percentages of properly developed embryos based on the picked properly developed E6.5 and E8.5 starting embryo numbers. Scale bars from **A-B** represent 200 μm. n = number of embryos evaluated. Abbreviations: PICM, peri-implantation culture medium; EUCM, ex utero culture medium; FBS, fetal bovine serum; RS, rat serum.

**Extended Data Figure 5.**
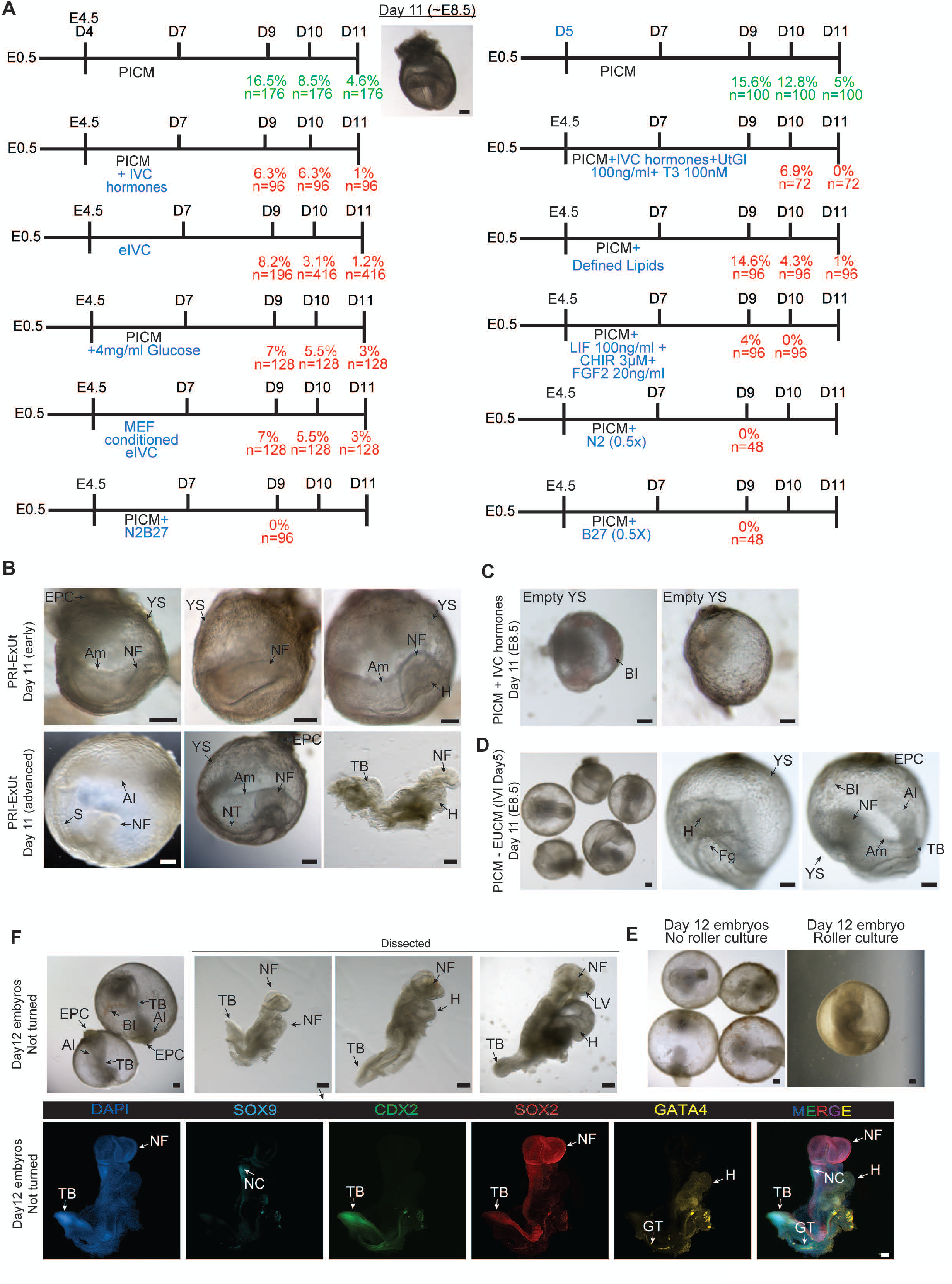
Optimization of PRI-ExUt culture conditions for mouse zygote development beyond gastrulation. **A,** Schematic representation comparing culture optimizations for PRI-ExUt relating to evaluating effect of supplementing PICM during IVI-like stage (day4-7) with the indicated supplements. Developmental outcome and efficiencies at days 9-11 (E6.5-E8.5) are indicated. In all conditions, EUCM was used after Day 7 (**Extended Data** Fig. 1). n = number of embryos evaluated. **B,** Representative bright-field images of early and advanced ∼E8.5 embryos generated from mouse zygotes using the optimized PRI-ExUt regimen. **C,** Representative success outcomes after applying PRI-ExUt in which PICM was supplemented with added hormonal components that are typical additions to previously described IVC media. IVC-associated hormonal components lead to a sharp decline in efficiency on D11, leading to largely empty yolk sac. **D,** Representative day 11 embryos following transfer to PRI-ExUt conditions at day 5 instead of E4.5. **E,** Representative day 12 embryos showing lack of further growth when maintained without roller culture unlike the positive control cultured with roller culture that has properly developed and reached Day 12 (corresponding to ∼E9.5). **F,** Representative day 12 embryos showing failure of axial turning when maintained without roller culture. A corresponding whole-mount immunofluorescence for SOX9, CDX2, SOX2 and GATA4 of an unturned day 12 embryo is shown. All scale bars represent 200 μm. Abbreviations: Al, allantois; Am, amnion; BI, blood island; CHIR, CHIR99021; eIVC, enhanced in vitro culture; EPC, ectoplacental cone; EUCM, ex utero culture medium; Fg, foregut; GT, gut tube; H, heart; IVC, in vitro culture; IVI, in vitro implantation; LIF, leukemia inhibitory factor; LV, left ventricle; MEF, mouse embryonic fibroblast; NC, neural crest; NF, neural fold; NT, neural tube; S, somite; T3, triiodothyronine; TB, tail bud; UtGl, uteroglobin; YS, yolk sac.

**Extended Data Figure 6.**
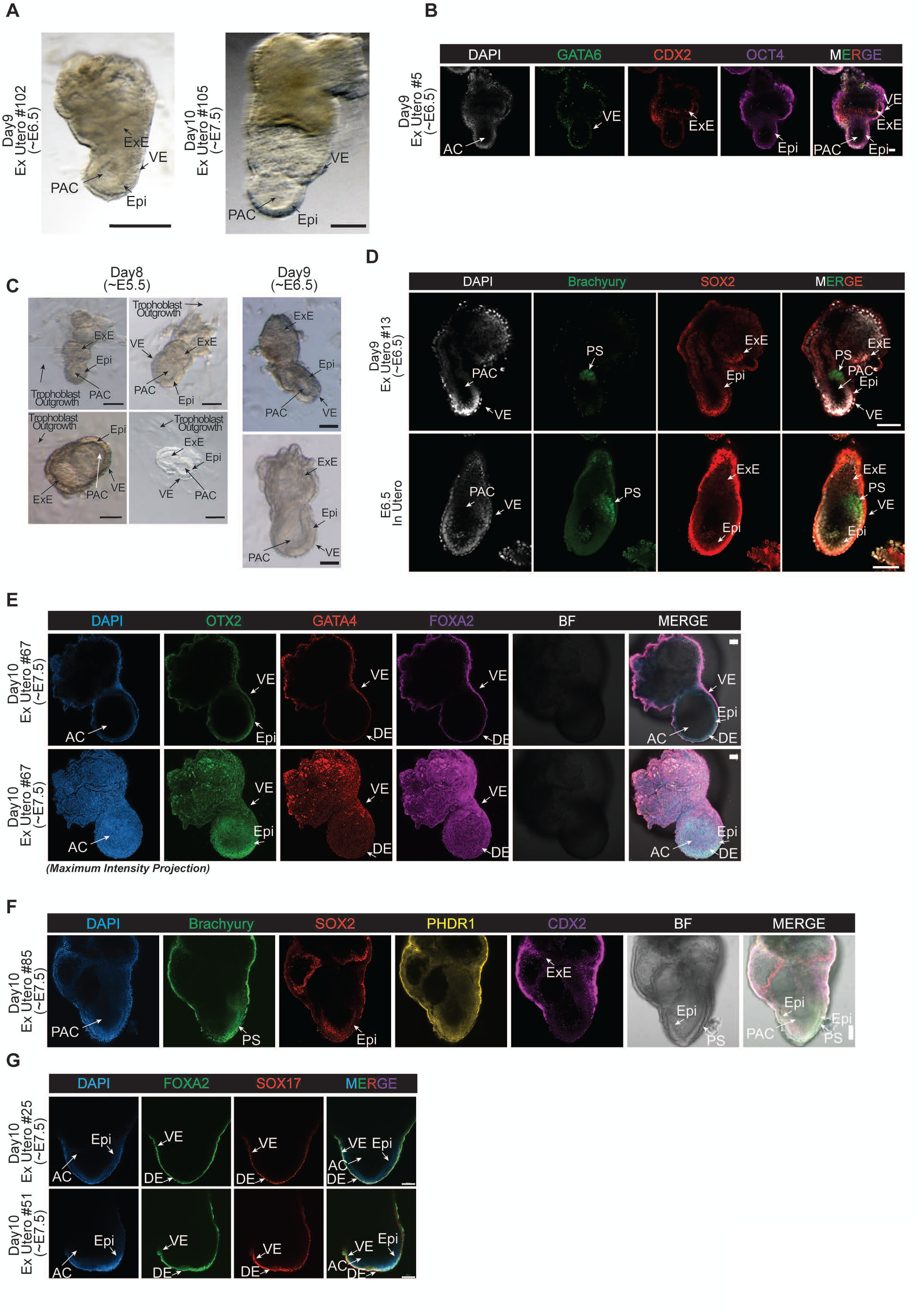
Characterization of egg-cylinder stage embryos generated with PRI-ExUt culture. **A,** Representative images of day 9 (∼E6.5) and day 10 (∼E7.5) embryos cultured under PRI-ExUt conditions. Scale bars, 100 μm. **B**, Representative immunostainings of day 9 PRI-ExUt cultured embryos stained for the Epi, VE and ExE markers OCT4, GATA6, and CDX2, respectively. Scale bars, 50 μm. **C,** Morphological progression of PRI-ExUt embryos from E5.5 to E6.5-equivalent stage. Scale bars, 50 μm. **D,** Whole-mount immunostaining of day 9 PRI-ExUt embryos stained for BRACHYURY (Primitive Streak), and SOX2 (ExE and Epi) compared with stage-matched in utero embryo. Scale bars, 100 μm. **E,** Middle section and maximum intensity projection confocal images for OTX2, GATA4 and FOXA2 expression in day 10 PRI-ExUt embryos. Scale bars, 100 μm. **F,** Representative immunostainings of day 10 PRI-ExUt cultured embryos stained for BRACHYURY (PS), and SOX2 (ExE and Epi), CDX2 (ExE) and PHDR1 (cell membranes). Scale bars, 100 μm. **G,** Representative immunostainings of day 10 PRI-ExUt cultured embryos stained for FOXA2 and SOX17 (labeling VE and DE). Scale bars, 100 μm in **D-G**. Abbreviations: AC, amniotic cavity; DE, definitive endoderm; Epi, epiblast; ExE, extraembryonic ectoderm; PAC, pro-amniotic cavity; PS, primitive streak; VE, visceral endoderm.

**Extended Data Figure 7.**
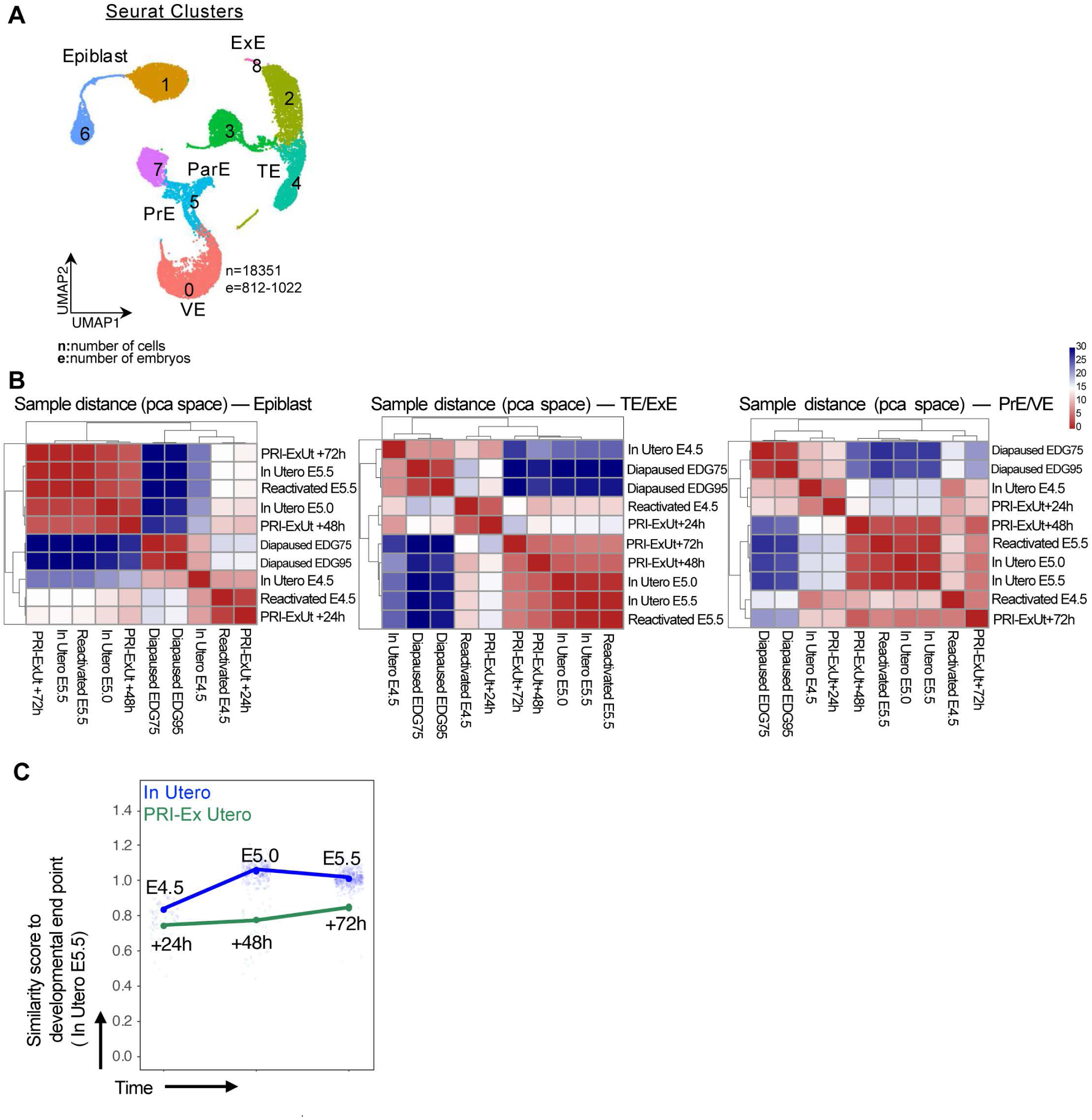
PRI-ExUt embryos exhibit delayed peri-implantation development without entering diapause. **A,** Seurat clusters of the integrated peri-implantation UMAP (shown in Fig. 1i). The lineages the clusters correspond to are as follows: 1: epiblast; 6: diapaused epiblast; 2+4+8: ExE ectoderm; 3: TE + diapaused TE; 5: ParE and PrE; 7: diapaused PrE. n = number of cells; e = number of embryos. **B,** Transcriptional similarity represented by sample distance in PCA space of epiblast, extraembryonic ectoderm (TE and ExE) and extraembryonic endoderm (PrE and VE) populations across PRI-ExUt, diapause, and in utero peri-implantation protocols. Notably, diapause samples (Diapaused EDG75 and Diapaused EDG95) cluster separately from PRI-ExUt samples. **C,** Average similarity score to mature Seurat cluster in epiblast lineage showing the dynamics of differentiation in embryos grown in PRI-ExUt protocol (green) vs. in utero (blue). The similarity score was calculated with Seurat AddModuleScore(), based on top 1000 genes in the following Seurat clusters: epiblast-1, Endoderm-0, Ectoderm-2&4. PRI-ExUt+24h, PRI-ExUt+48h and PRI-ExUt+72h correspond to +24 hrs., +48 hrs. and +72 hrs. after in vitro implantation respectively, while in utero samples correspond to In Utero E4.5, In Utero E5.0 and In Utero E5.5. PRI-ExUt embryos demonstrate a mostly slow and steady transcriptional progression, different from diapause. Abbreviations: EDG75 and EDG95, estimated day of gestation 7.5 and 9.5. ExE, extraembryonic ectoderm; ParE, parietal endoderm; PrE, primitive endoderm; PRI-ExUt, pre-implantation-initiated ex utero culture; TE, trophectoderm; VE, visceral endoderm.

**Extended Data Figure 8.**
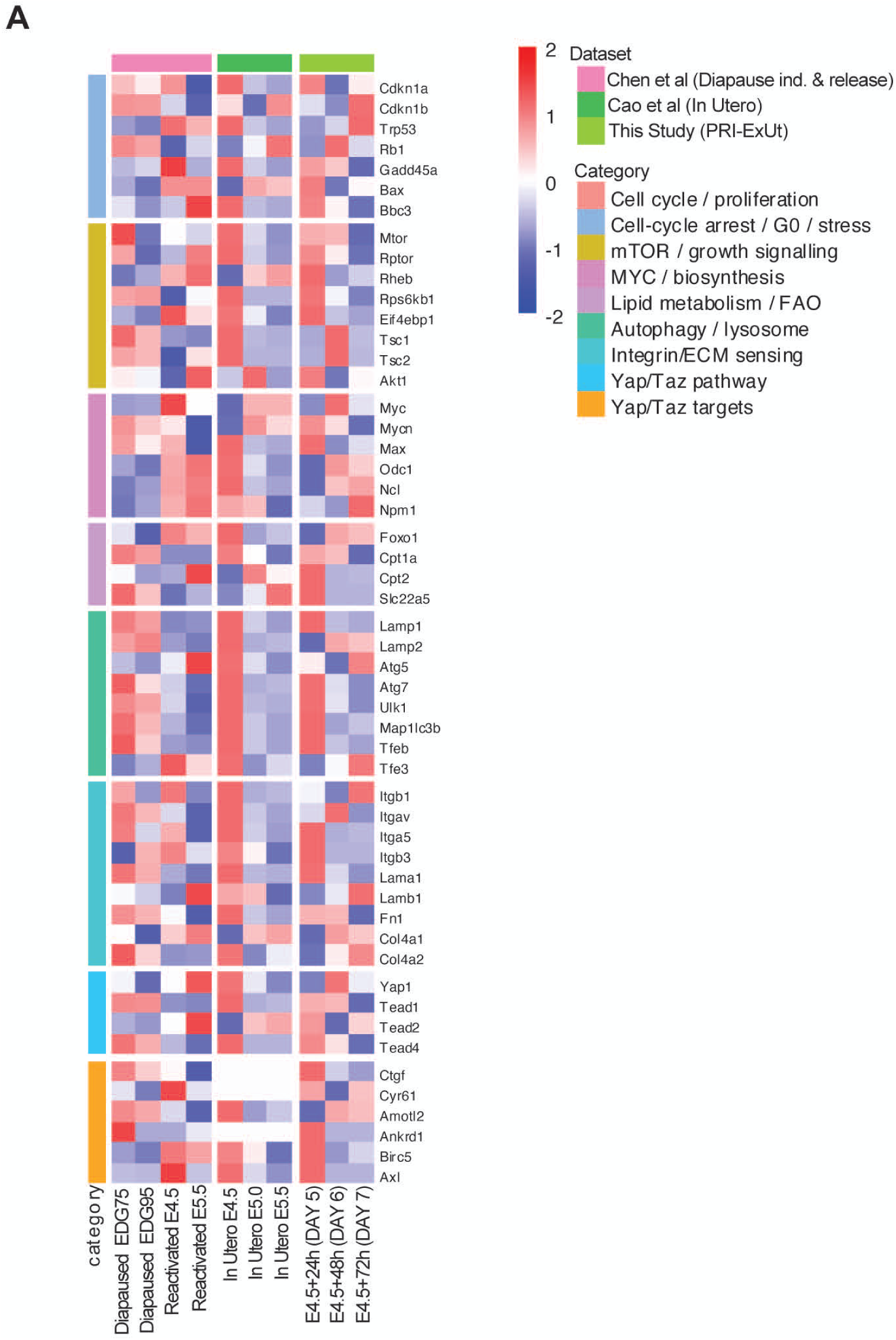
Diapause-associated gene programs are not selectively activated in PRI-ExUt embryos. **A,** Heatmaps showing normalized expression (z-score) of gene programs associated with diapause, proliferation/cell-cycle arrest, mTOR/growth signaling, MYC/biosynthetic activity, lipid metabolism, autophagy, extracellular-matrix sensing and YAP/TAZ signaling in epiblast populations from published diapause^69^, in utero^68^ and PRI-ExUt datasets. Abbreviations: ECM, extracellular matrix; EDG75 and EDG95, estimated day of gestation 7.5 and 9.5; FAO, fatty-acid oxidation; G0, quiescent cell-cycle state; PRI-ExUt, pre-implantation-initiated ex utero culture.

**Extended Data Figure 9.**
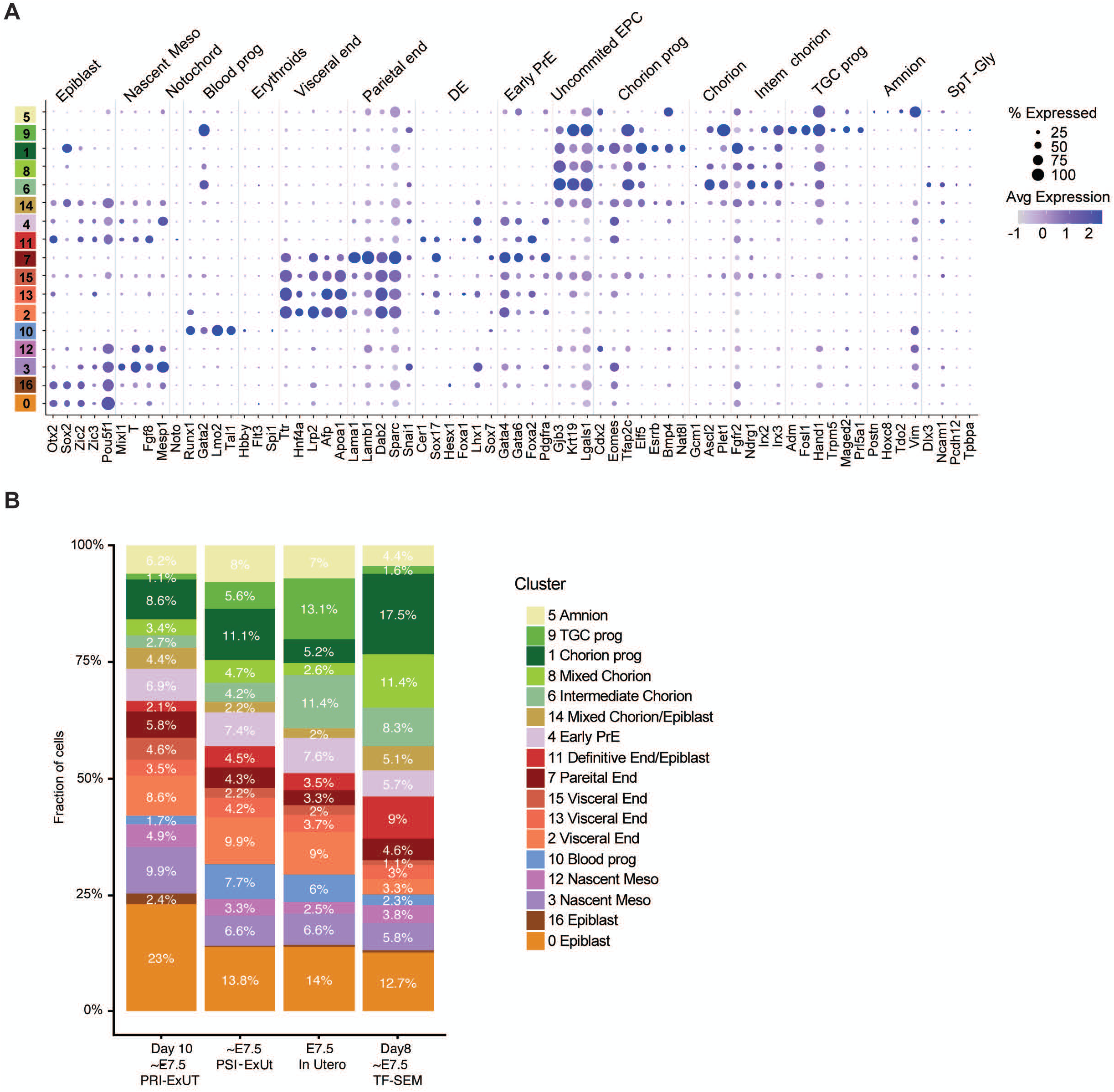
Single cell transcriptomic characterization of day 10 (E7.5) PRI-ExUt embryos. **A,** Dot plot illustrating the expression of selected lineage markers across annotated E7.5 cell populations. Dot size denotes percentage of cells expressing the marker, and shades indicate normalized average expression. **B,** Relative abundance of annotated cell populations in day 10 PRI-ExUt, E7.5 PSI-ExUt, E7.5 in utero and day 8 (∼E7.5) TF-SEM samples. Abbreviations: DE, definitive endoderm; EPC, ectoplacental cone; InUt, in utero; Meso, mesoderm; PrE, primitive endoderm; PRI-ExUt, pre-implantation-initiated ex utero culture; prog, progenitor; PSI-ExUt, post-implantation-initiated ex utero culture; SpT-Gly, spongiotrophoblast/glycogen trophoblast; TF-SEM, transgene-free stem-cell-derived embryo model; TGC, trophoblast giant cell.

**Extended Data Figure 10.**
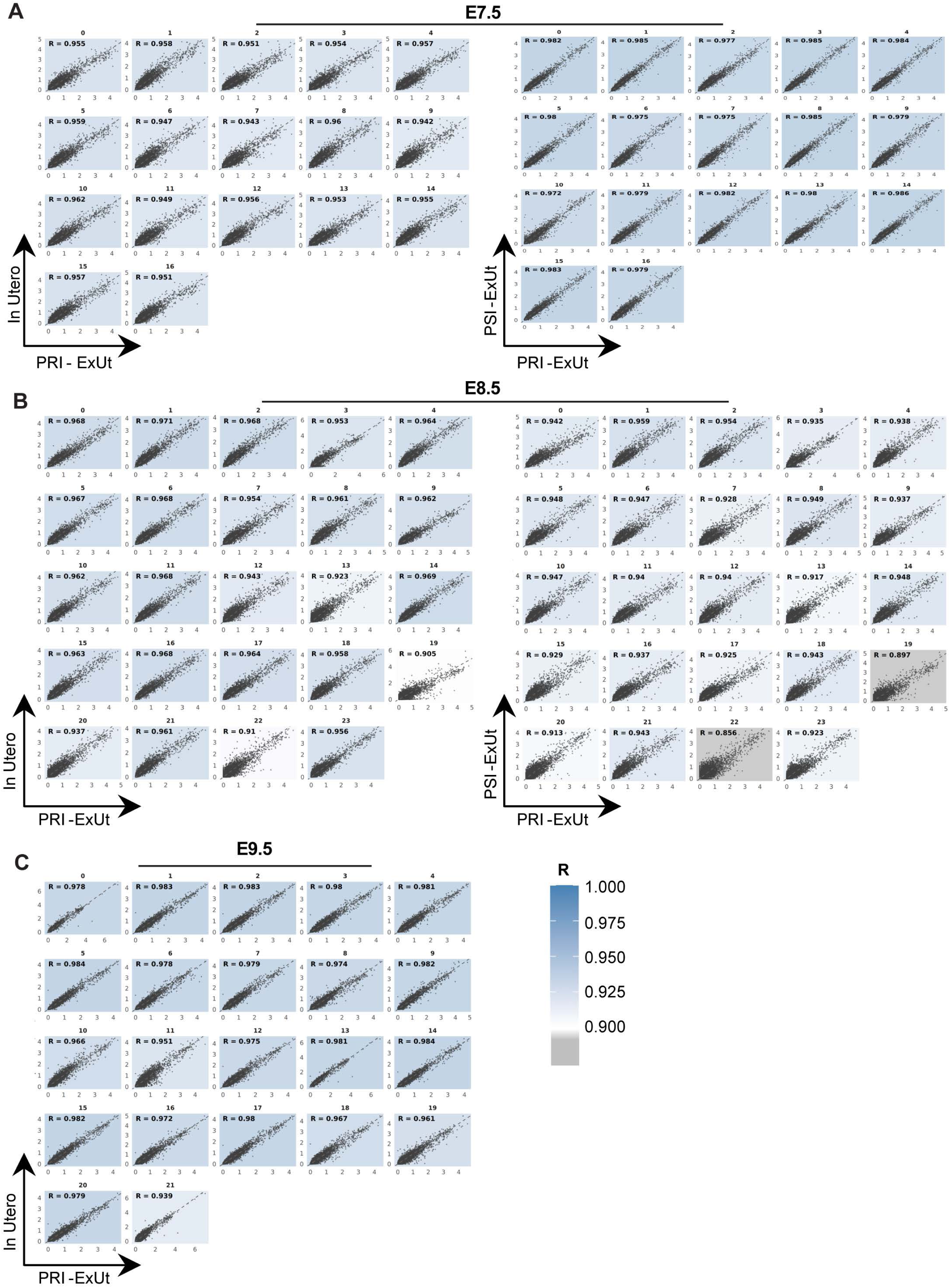
Transcriptomic correlation of PRI-ExUt embryos with stage-matched natural embryos. **A-C,** Scatter plots comparing average normalized gene expression across all genes between PRI-ExUt and in utero or PSI-ExUt cells within matched cell populations at E7.5 (**A**), E8.5 (**B**), and E9.5 (**C**). X and Y axes = average normalized expression. Each plot indicates the Pearson correlation coefficient (*R*); shading denotes correlation strength. These plots show the high correlation between the transcriptional patterns of the different embryo growth protocols.

**Extended Data Figure 11.**
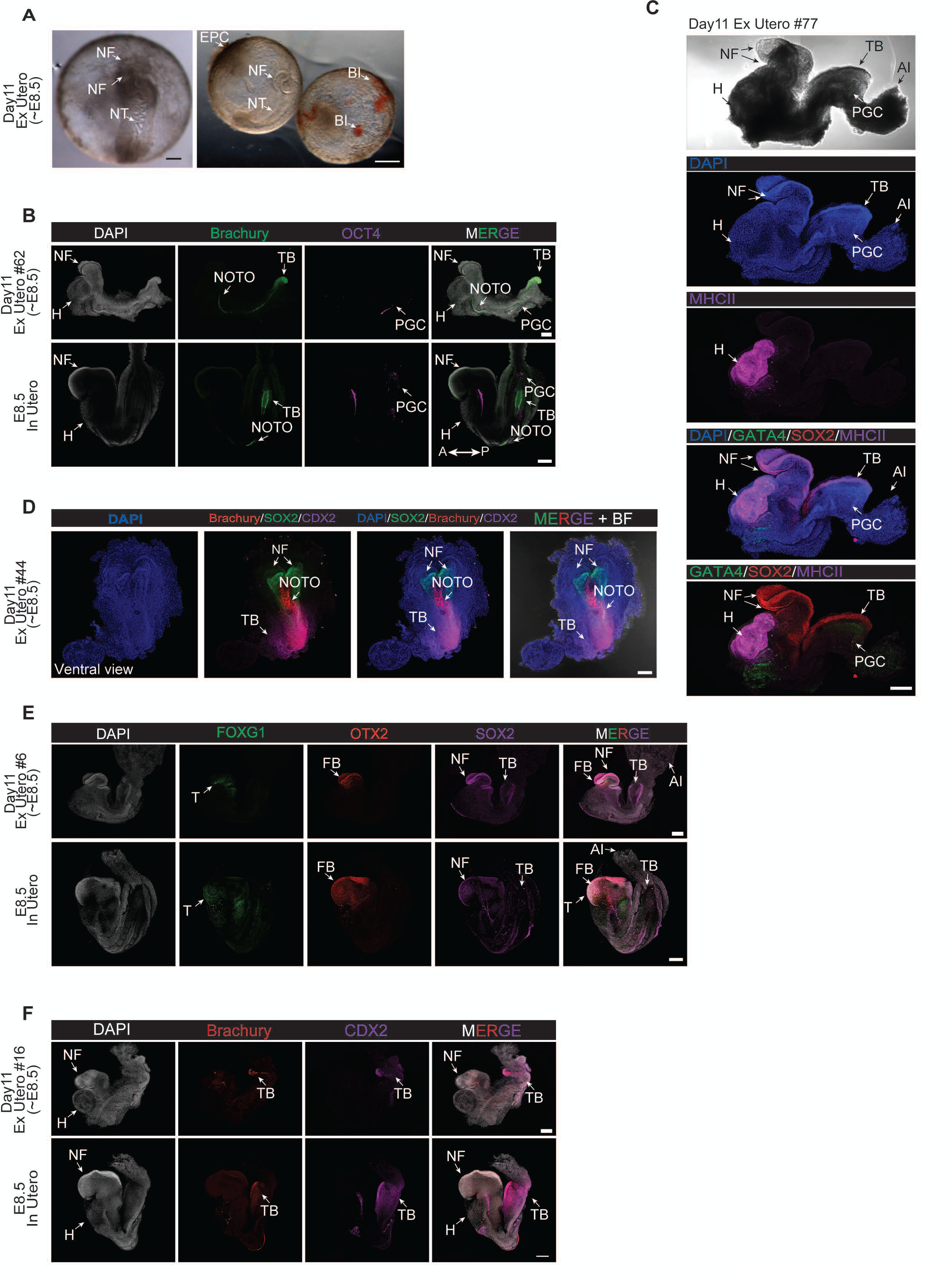
Characterization of PRI-ExUt embryos during early organogenesis. **A,** Representative images of day 11 (∼E8.5) embryos cultured under PRI-ExUt conditions with correct morphology. **B,** Whole-mount immunofluorescence of day 11 PRI-ExUt embryos and developmentally matched in utero E8.5 embryo for BRACHYURY and OCT4, showing notochord (NOTO)/ tailbud (TB) domains and primordial germ cells (PGCs). **C,** Representative immunostainings for GATA4 (Heart and Gut Endoderm), SOX2 (Neural domains, PGCs) and MHC-II (Heart) in a day 11 PRI-ExUt embryo. **D,** Representative ventral view immunostaining for BRACHYURY, SOX2 and CDX2 in day 11 PRI-ExUt embryos. Scale bars represent 100 μm. **E-F,** Representative immunostainings of day 11 PRI-ExUt embryos and stage-matched in utero E8.5 embryos showing correct localization of anterior markers FOXG1 and OTX2 (**E**) and posterior markers BRACHYURY and CDX2 (**F**). Scale bars in **A-C**, **E-F**, 200 μm. Abbreviations: A, anterior; Al, allantois; BI, blood island; EPC, ectoplacental cone; FB, forebrain; H, heart; HB, hindbrain; MHC-II, myosin heavy chain II; NF, neural fold; NOTO, notochord; P, posterior; PGC, primordial germ cell; T, telencephalic primordium; TB, tail bud.

**Extended Data Figure 12.**
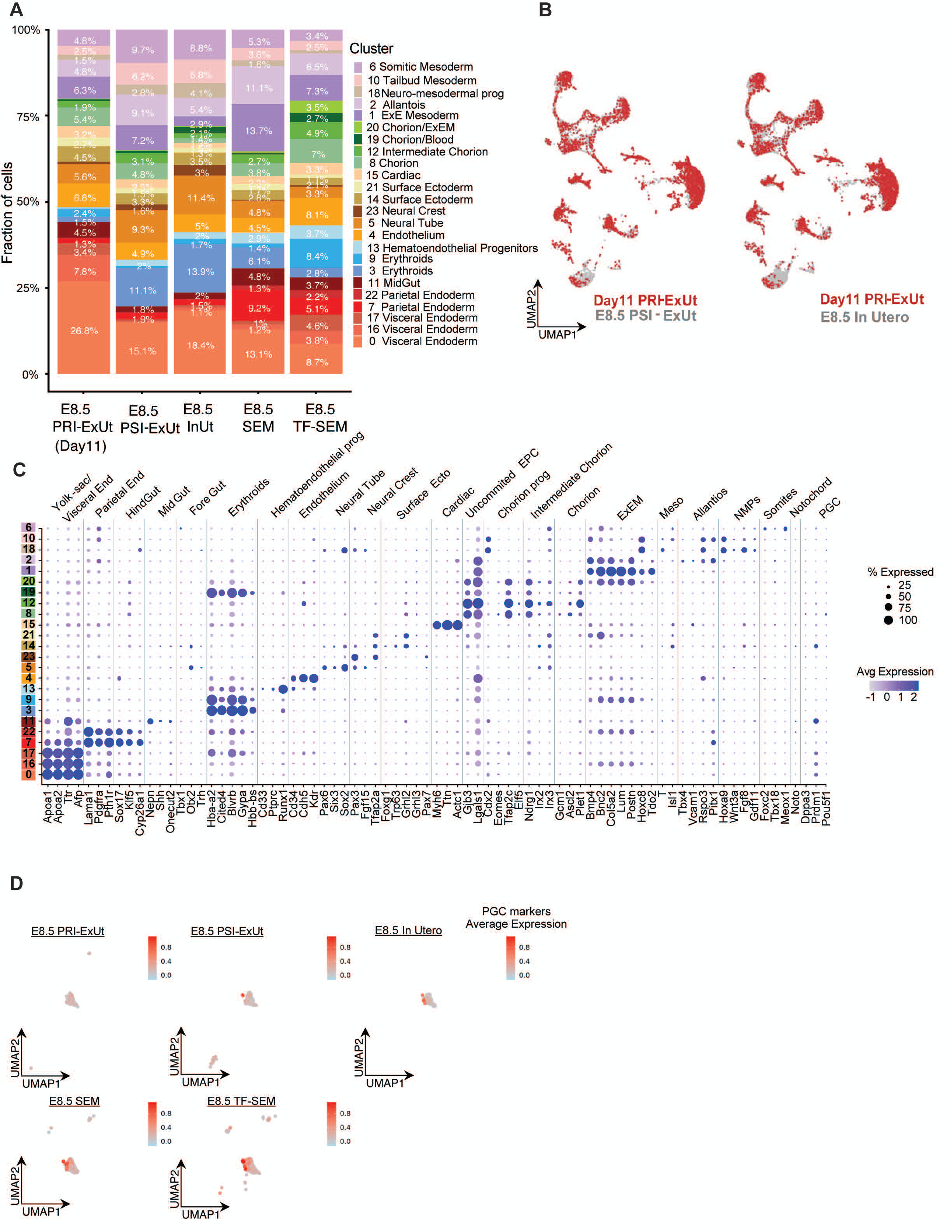
Single-cell transcriptomic comparison of day 11 PRI-ExUt embryos with stage-matched PSI-ExUt, in utero and SEMs. **A,** Relative abundance of annotated cell populations in day 11 PRI-ExUt, PSI-ExUt, E8.5 in utero, SEM and TF-SEM samples. **B,** UMAP transcriptomic comparison of day 11 PRI-ExUt with PSI-ExUt or E8.5 in utero embryos. **C,** Dot plots illustrating the expression of selected lineage markers across annotated cell populations. Dot size denotes the percentage of cells expressing the marker, and shades indicate normalized average expression in these cells. **D,** Average expression of PGC markers (Pou5f1, Nanog, Klf4, Sox2, Dppa3, Prdm1, Bmp4, Prdm14, Alpl & Nanos3) across cluster 8 of the integrated E8.5 manifold (PRI-ExUt, PSI-ExUt, E8.5 in utero, SEM and TF-SEM samples), showing a similar expression of PGC markers in a handful of cells across different embryo models. Abbreviations: EPC, ectoplacental cone; ExEM, extraembryonic mesoderm; InUt, in utero; NMP, neuromesodermal progenitor; PGC, primordial germ cell; PRI-ExUt, pre-implantation-initiated ex utero culture; prog, progenitor; PSI-ExUt, post-implantation-initiated ex utero culture; SEM, stem-cell-derived embryo model; TF-SEM, transgene-free stem-cell-derived embryo model.

**Extended Data Figure 13.**
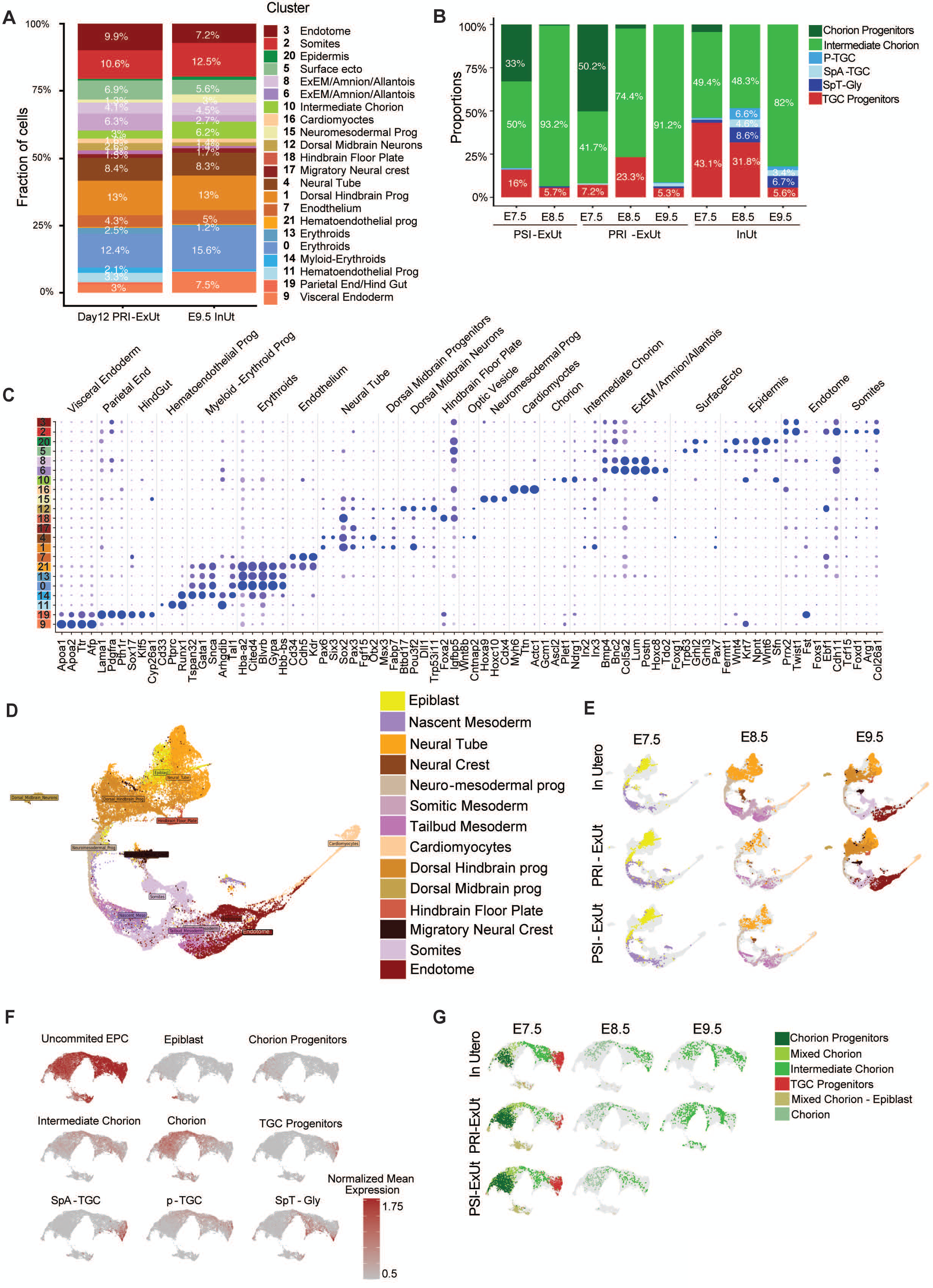
Single cell transcriptomic and lineage progression analysis of day 12 PRI-ExUt embryos across stages up to E9.5. **A,** Cell type composition of day 12 PRI-ExUt and E9.5 in utero embryos. **B,** Relative abundance of trophoblast/chorion-associated populations across E7.5, E8.5 and E9.5 in PRI-ExUt, PSI-ExUt and in utero samples. Cell annotation as in (F). Notably, embryos grown ex utero show reduced numbers of, or completely lack, advanced TGC types: p-TGC, SpA-TGC, SpT-Gly. **C,** Dot plot illustrating the expression of selected markers in E9.5 cell clusters. Dot size denotes the percentage of cells expressing the marker, and shades indicate normalized average expression in these cells. Color code is as in A. **D,** Integrated UMAP of neuromesodermal-lineage populations across developmental stages. **E,** Temporal distribution of epiblast and mesoderm-derived cell states throughout E7.5, E8.5 and E9.5-equivalent stages in PRI-ExUt, PSI-ExUt and in utero embryos, showing similar dynamics of neuro-mesodermal lineage development across embryo growth protocols. Colors as in (D). **F,** Mean normalized expression of ExE-derived lineages as follows: Uncommitted_EPC (Gjb3, Krt19, Lgals1, Cald1, Ctsl), Epiblast (Otx2, Sox2, Pou5f1), Chorion_progenitors (Sox3, Bmp4, Sox2), Chorion (Irx4, Esx1, Id1, Id3, Phlda2, Klhl13), Intermediate_Chorion (Ascl2, Cited1, Gjb3, Ndrg1, Irx2, Irx3), TGC_progenitors (Adm, Fosl1, Hand1, Trpm5, Maged2, Prl5a1), p_TGC (Serpinb9d, Hsd3b6, Rhox6, Cts7, Prl2c3), SpA_TGC (Ctla2a, Pecam1), SpT_Gly (Dlx3, Car2, Ncam1, Tpbpa). **G,** UMAP distribution of ExE-derived lineage populations spanning E7.5, E8.5 and E9.5 PRI-ExUt, PSI-ExUt and in utero samples from E7.5 to E9.5. Abbreviations: EPC, ectoplacental cone; ExEM, extraembryonic mesoderm; InUt, in utero; p-TGC, parietal trophoblast giant cell; PRI-ExUt, pre-implantation-initiated ex utero culture; prog, progenitor; PSI-ExUt, post-implantation-initiated ex utero culture; SpA-TGC, spiral artery-associated trophoblast giant cell; SpT-Gly, spongiotrophoblast/glycogen trophoblast; TGC, trophoblast giant cell.

**Extended Data Figure 14.**
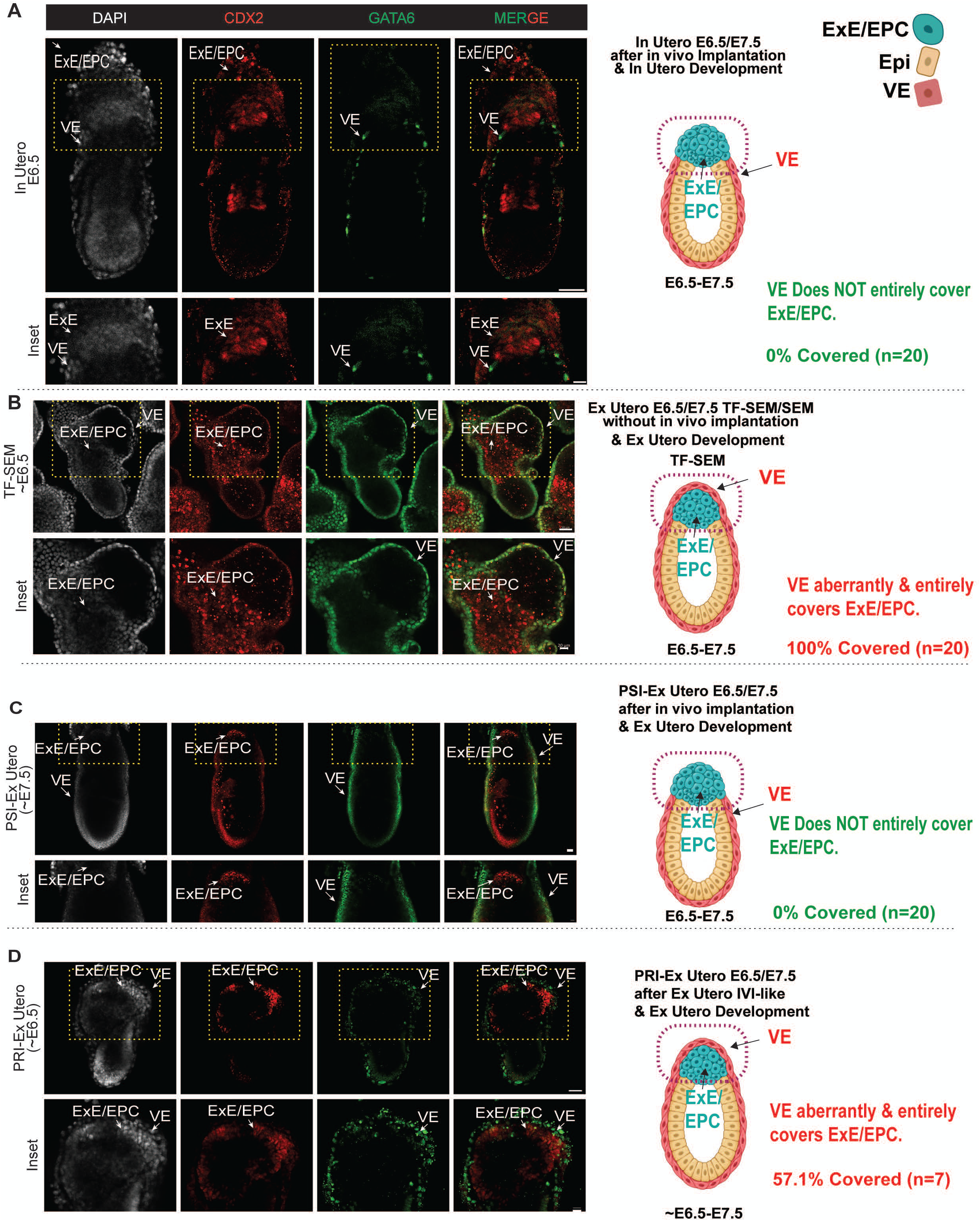
Aberrant extra-embryonic organization in mouse SEMs and PRI-ExUt compared to other in utero or ex utero grown embryos. **A,** Representative immunostaining of ExE and VE lineages marked by CDX2 and GATA6, respectively, of in utero embryos that had undergone in utero implantation. VE lineage does not cover the ExE compartment in the total number of embryos immunostained (n=20). The image is intentionally reused from Figure 1, with altered markings to emphasize the relevant region being discussed and compared with other samples below. **B,** Representative immunostaining of ExE and VE lineages marked by CDX2 and GATA6, respectively, of TF-SEM/SEMs that developed ex utero. VE lineage aberrantly covers the ExE and EPC compartment in 100% of the total number of embryos immunostained (n=20). **C,** Representative immunostaining of ExE and VE lineages marked by CDX2 and GATA6, respectively, of post-implantation natural embryos that developed ex utero from E6.5. VE lineage does not cover the ExE compartment in the total number of embryos immunostained (n=20). **D,** Representative immunostaining of ExE and VE lineages marked by CDX2 and GATA6, respectively, of PRI-ExUt embryos that developed ex utero. VE lineage aberrantly covers the ExE compartment in 57.1% of the total number of embryos immunostained (n=7). The image is intentionally re-used from Figure 1H. Schemes on the right side of each immunostaining set in A-D illustrate simplistically VE vs. ExE/EPC organization.

**Extended Data Figure 15.**
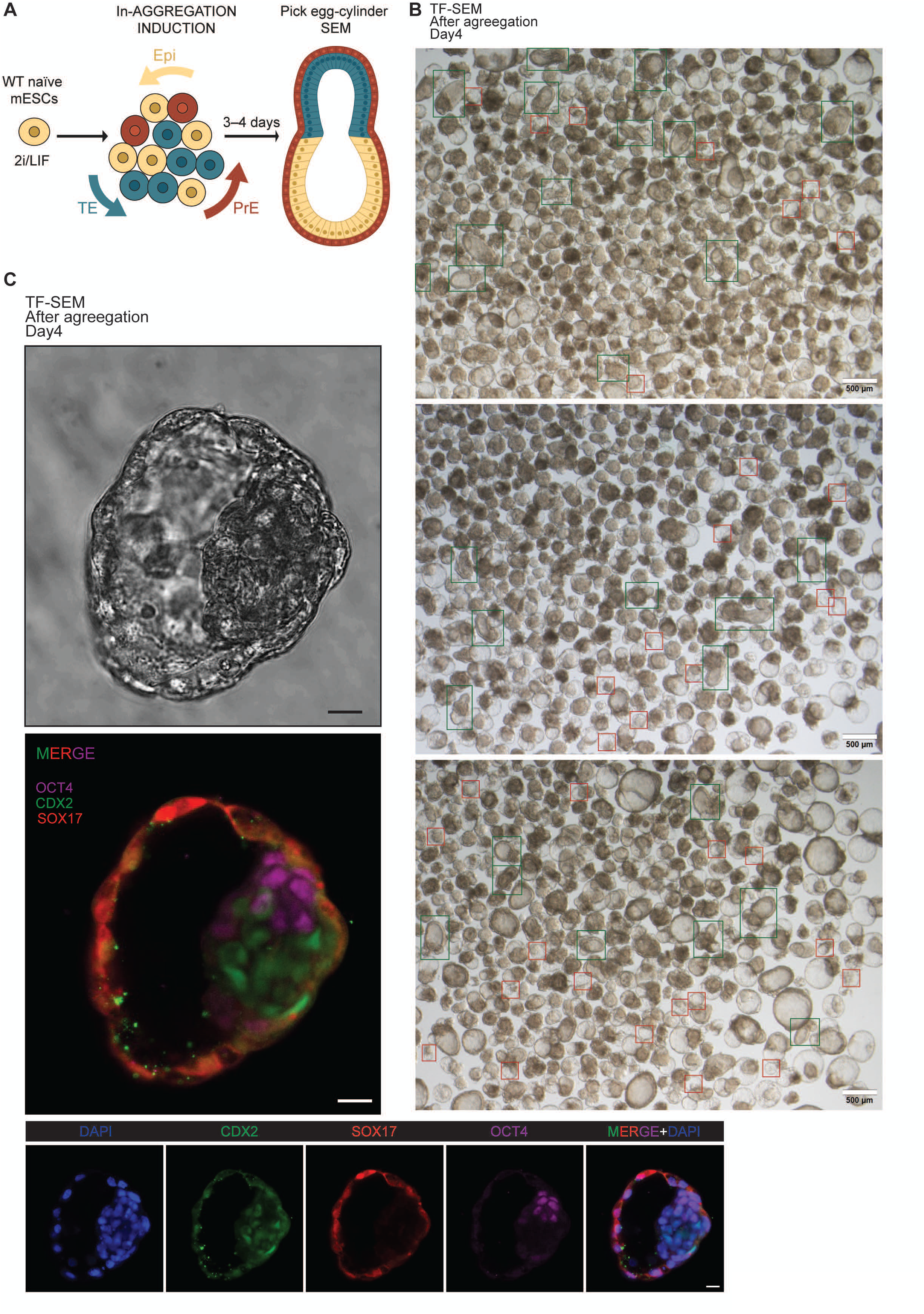
Characterization of cavitated structures generated during mouse SEM generation excludes possible formation of bona fide blastoids. **A,** Schematic representation of the correct egg-cylinder structure generation during mouse SEM protocol involving chemical induction of extra-embryonic alongside embryonic cell mixtures, as previously reported^72^ . **B,** Stereomicroscope images of the structures generated during the TF-SEM protocols. The green boxes indicate correct egg-cylinder morphology that is chosen for further growth, while red boxes indicate cavitated structures that resemble blastoids. Scale bars are indicated on the image. **C,** Representative brightfield image of such blastoid-like structures generated during TF-SEM protocol, however immunostaining of such structures for CDX2, SOX17 and OCT4, confirms they are entirely surrounded by SOX17+ PrE cells and not CDX2+ TSC cells which are rather found inside the false ICM-like region, thus excluding formation of bona fide blastocyst-like embryo models. Scale bars indicated 20 μm.

**Extended Data Figure 16.**
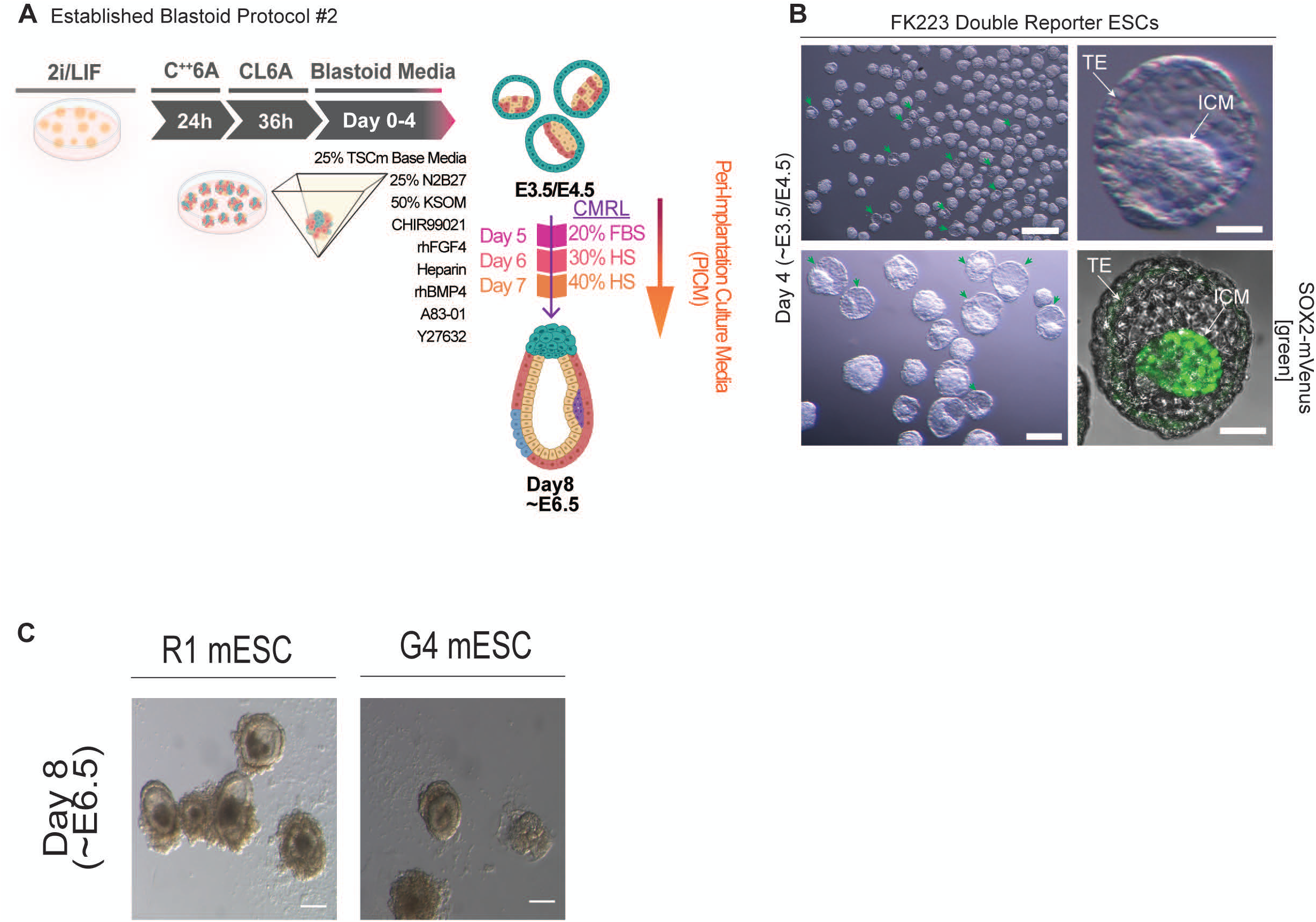
Evaluating in vitro developmental potential of mouse blastoids generated following applying chemically induced embryo-founder conditions. **A,** Schematic representation of the blastoid generation protocol following sequential chemical induction of embryo founder cells as previously published^60^ and subsequent transfer to PRI-ExUt culture. **B,** Representative day 4 blastocyst-like structures generated using FK223 double reporter ESCs (SOX2-mVenus; T:mCherry) following chemical founder cell induction, confirming SOX2+ Epiblast within ICM as expected. Scale bars, 100 μm (left panel), 25 μm (right panel). **C,** Representative stereomicroscope images of day 8 (∼E6.5) structures following transfer to PRI-ExUt conditions of chemically induced blastocyst-like structures from R1 or G4 ESC lines, indicating lack of an ExE compartment in attached structures, making them inherently abnormal. Similar results were obtained from all 4 mouse lines tested: V6.5, V6.5 reporter line, R1 and G4 mouse ESCs. Scale bars, 50 μm. Abbreviations: FBS, fetal bovine serum; HS, human serum; ICM, inner cell mass; TE, trophectoderm; TSCm, trophoblast stem-cell medium.

**Extended Data Figure 17.**
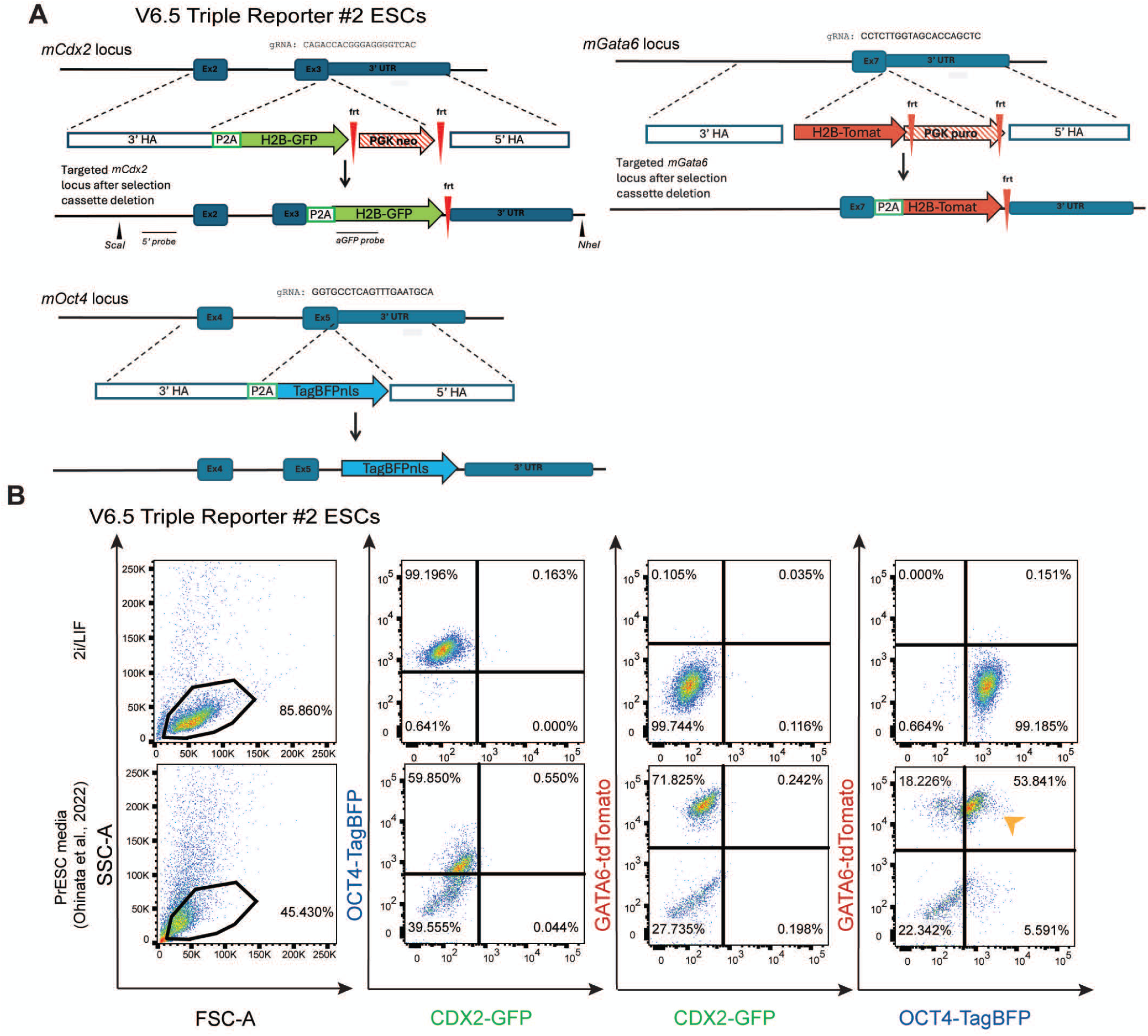
Targeting scheme and characterization of ESC derived PrESCs with OCT4/CDX2/GATA6 triple-reporter line. **A,** Schematic overview of the targeting strategy used to generate the V6.5 triple-reporter ESC line carrying OCT4-BFP, CDX2-GFP and GATA6-tdTomato reporters. Clone#2 was validated as correctly targeted and used for further analysis. **B,** Flow cytometry analysis of reporter expression in cells maintained in 2i/LIF or transferred to PrESC medium^75^, which results in a sub-population consisting of OCT4+GATA6+ (note orange arrowheads) and CDX2-negative, that can be further subcloned or purified by sorting. Abbreviations: HA, homology arm; NLS, nuclear localization signal; P2A, porcine teschovirus-1 2A peptide; PGK, phosphoglycerate kinase promoter; PrESC, primitive endoderm stem cell.

**Extended Data Figure 18.**
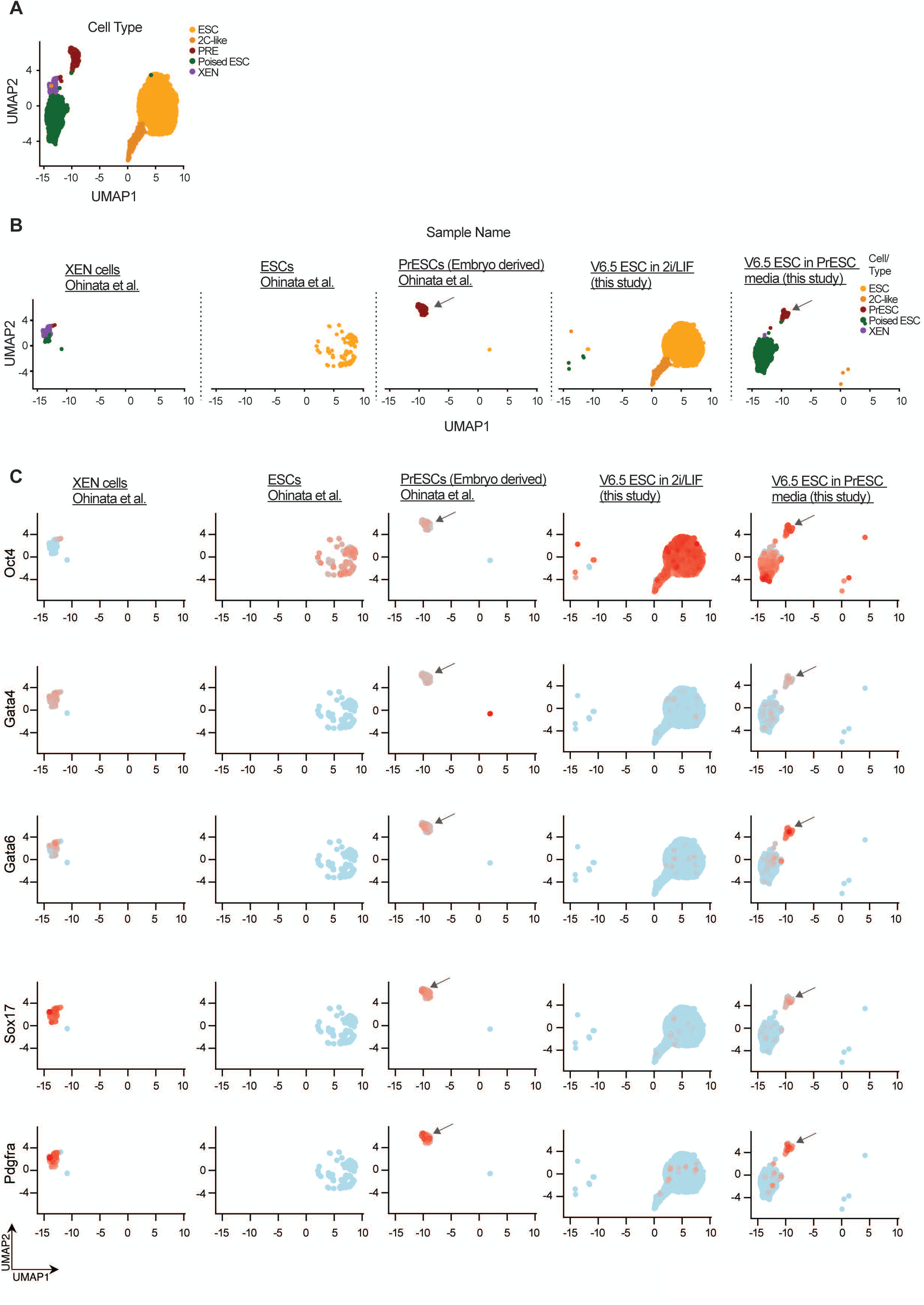
PrESC can be in vitro induced from mouse ESCs. **A,** Integrated single cell RNA-seq UMAP representing single-cell RNA-seq measured in V6.5 ESCs expanded for 2 passages in 2i/LIF conditions or PrESC medium (Ohinata et al. Science2022)^75^, as well as 3 previously published samples (Ohinata et al. Science2022)^75^ measured in ESC, XEN and embryo derived PrESC. Colors indicate the following cell types: ESC, 2C-Like, PrESC, Poised ESC and XEN. **B,** UMAP representation of (a), divided by sample. Colors indicate cell types. **C**, Expression value of the indicated genes, presented on the UMAP and divided by samples. ESCs and PrESCs, but not XEN cells, express the Oct4 pluripotency marker, whereas the PrE markers Gata4, Gata6, Sox17 and Pdgfra are expressed only in PrESCs and XEN cells, and not in ESCs. This scRNA-seq based analysis indicated that bona fide PrESC subpopulation can be induced from in vitro grown mouse ESCs after short expansion in PrESC media, and not only from blastocysts as previously described^75^.

**Extended Data Figure 19.**
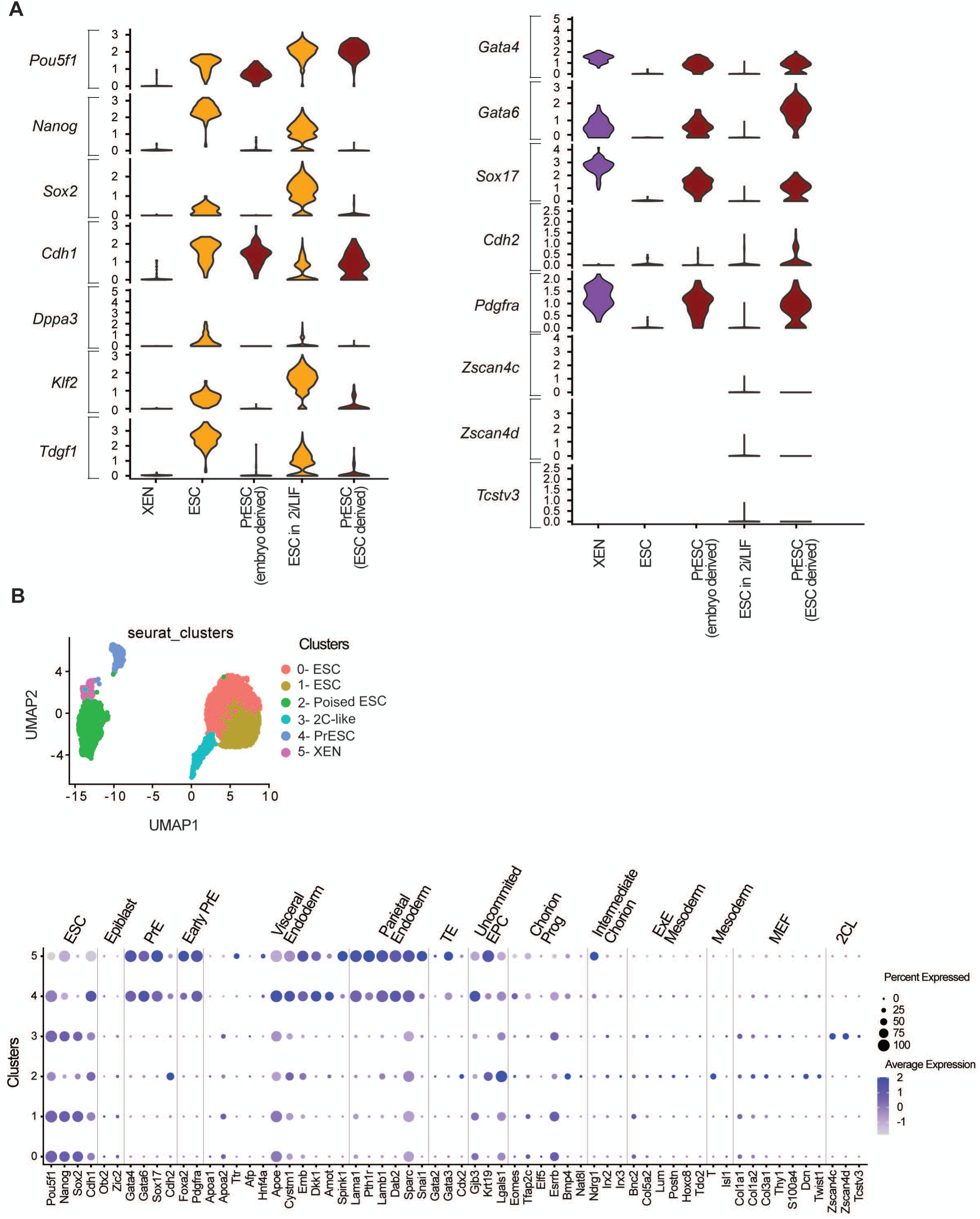
Transcriptional identity of in vitro derived PrESCs from mouse ESCs. **A,** Expression distribution of selected genes in selected clusters as following: XEN, ESC and embryo derived PrESCs^75^, ESC in 2i/LIF and ESC derived PrESC cluster (this study). In vitro induced PrESCs are transcriptionally similar to previously described embryo derived PrESCs^75^, and not to XEN or ESCs. **B,** Integrated single cell RNA-seq UMAP representing single-cell RNA-seq measured in the samples shown in **A** and the identity of 6 cell clusters obtained. At the lower part of the panel, dot plot illustrating the expression of selected lineage markers across annotated Seurat clusters described above in same panel. Dot size denotes percentage of cells expressing the marker, and shades indicate normalized average expression.

**Extended Data Figure 20.**
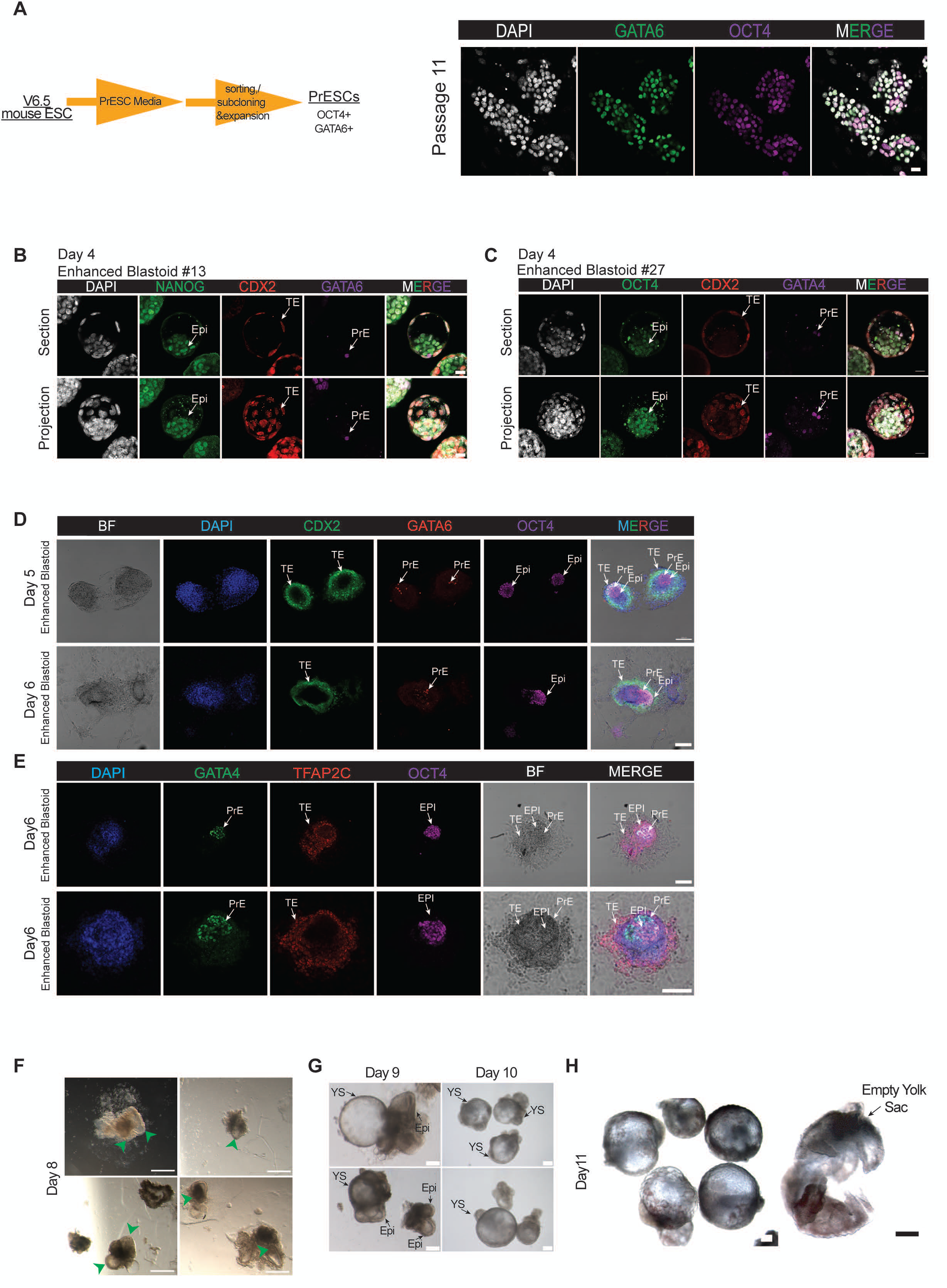
Characterization and developmental potential of PrESC-enhanced blastoids. **A**, Uniform OCT4 and GATA6 co-expression in V6.5 ESC derived and sorted PrESC that were induced and maintained in PrESC conditions at the indicated passage. Scale bars, 20 μm. **B, C,** Representative confocal images of PrESC-enhanced blastoids showing NANOG^+^/OCT4^+^ epiblast, GATA4^+^/GATA6^+^ PrE and CDX2-positive TE compartment. Scale bars, 20 μm. **D-E,** Lineage organization at day 5 and 6 of PRI-ExUt culture applied on the enhanced blastoids, as assessed by immunostaining for CDX2 or TFAP2C together with GATA6/GATA4 and OCT4. Scale bars, 100 μm. **F-G,** Representative stereomicroscope images of day 8-10 PRI-ExUt cultured enhanced blastoids generated with the enhanced blastoid protocol. The images at day 9 show residual epiblast compartments while day 10 showed empty yolk sacs that lack an embryo proper. Green arrowheads point to properly developed structures. Scale bars, 200 μm. **H,** Representative day 11 structures from enhanced blastoids showing continued yolk-sac expansion while lacking an identifiable embryo proper. Scale bars, 200 μm. Abbreviations: AC, alternative condition; Epi, epiblast; PrE, primitive endoderm; PrESC, primitive endoderm stem cell; TE, trophectoderm; YS, yolk sac.

## Supplementary Videos Legends

**Supplementary Video 1. Time-lapse imaging of mouse blastocyst in vitro adherence and egg-cylinder morphogenesis during PRI-ExUt culture protocol.**

Representative time-lapse bright-field imaging of E4.5 blastocysts following transfer to PRI-ExUt in vitro implantation conditions optimized herein. Initially floating blastocysts attach to the uncoated culture surface, undergo trophoblast outgrowth and progressive post-implantation morphogenesis, ultimately generating elongated egg-cylinder-like structures. Time after initiation of imaging is indicated; total imaging time, 120 hours. Scale bar, 100 μm.

**Supplementary Video 2. Beating heart in E8.5 (protocol day 11) PRI-ExUt mouse embryo cultured ex utero from the zygote-stage.**

Representative live bright-field stereoscope imaging of a day 11 PRI-ExUt mouse embryo, equivalent to E8.5-stage, showing rhythmic cardiac contractions in an embryo displaying prominent neural folds, neural tube, tail bud and allantois. The embryo was dissected out from the yolk sac. Anatomical structures are indicated. Scale bar, 200 μm.

**Supplementary Video 3. Characterization of neural, notochord and posterior axial tissues in E8.5 PRI-ExUt mouse embryos.**

Three-dimensional rendering of whole-mount immunofluorescence comparing a day 11 PRI-ExUt embryo with a stage-matched E8.5 in utero embryo. Embryos were stained for BRACHYURY (green), SOX2 (red), CDX2 (magenta) and DAPI (blue), highlighting neural tissues and posterior axial domains, including the developing neural tube, notochord and tail-bud-associated regions. Scale bars, 200 μm.

**Supplementary Video 4. Characterization of neural and cardiac tissues in mouse E8.5 PRI-ExUt embryos.**

Three-dimensional rendering of whole-mount immunofluorescence comparing a day 11 PRI-ExUt embryo with a stage-matched E8.5 in utero embryo. Embryos were stained for MHC-II (magenta), SOX2 (green) and DAPI (blue), highlighting the developing heart and neural tissues. Scale bars, 200 μm.

**Supplementary Video 5. Characterization of forebrain, notochord and cardiac tissues in E8.5 PRI-ExUt mouse embryos.**

Three-dimensional rendering of whole-mount immunofluorescence comparing a day 11 PRI-ExUt mouse embryo with a stage-matched E8.5 in utero embryo (used as a reference control). Embryos were stained for FOXA2 (magenta), OTX2 (green), MHC-II (red) and DAPI (blue), highlighting notochordal and floor plate-associated FOXA2 domains, anterior neural OTX2 expression, and cardiac MHC-II expression. Scale bars, 100 μm.

**Supplementary Video 6. Morphology and cardiac activity of PRI-ExUt mouse embryos cultured from the totipotent zygote until mid-gestation (E9.5).**

Representative live bright-field stereoscope imaging of PRI-ExUt mouse embryos cultured for 12 days from the E0.5 zygote until ∼E9.5 stage. Embryo is shown within the intact yolk sac. Key anatomical structures are indicated. Scale bar, 250 μm.

**Supplementary Video 7. Comparative 3D reconstruction of organogenesis in E9.5 PRI-ExUt mouse embryos to in utero grown control embryos.**

Three-dimensional rendering of whole-mount immunofluorescence comparing a protocol day 12 PRI-ExUt embryo with a stage-matched E9.5 in utero embryo (used as a reference control). Embryos were cleared and stained for TUJ1 (magenta), SOX9 (red), GATA4 (green) and DAPI (blue), revealing extensive neuronal differentiation, SOX9-positive neural crest, bone/chondrogenic progenitors, and GATA4-positive cardiac and gut-associated tissues. Scale bars, 300 μm.

## Materials and Methods

### Animals

Female mice 5-8-week old CD1(ICR) were mated with CD1(ICR) or BDF1 male studs (obtained from Harlan or Charles Rivers). Female mice 5-8-week old CFW, C57BL/6 or BDF1 were mated with male studs from the matching genetic background (obtained from Harlan or Charles Rivers). Insemination was verified the next morning by the presence of a copulatory plug, and this day was defined as E0.5 days post coitum (d.p.c). All animal experiments were performed according to the Animal Protection Guidelines of the Howard Hughes Medical Institute Janelia Research Campus and the Weizmann Institute of Science and were approved by relevant HHMI Janelia IACUC (#24-0256.02), or Weizmann Institute IACUC (04640724-3, 07901123-1, 03370423-1, 09521222-1). All females used for ex utero culture of pre-implantation embryos were hormone primed, while females for obtaining post-implantation control embryos were not hormone primed. All animals had continuous access to water and food. All mice were housed under standard 12-hour light/12-hour dark cycle conditions in a specialized and certified animal facility.

### Mouse zygotes isolation and culture

Female mice (5-8-week old CD1(ICR) or BDF1) were super-ovulated by injecting 5 I.U. of pregnant mare serum gonadotropin (PMS), followed by 5 I.U. of human chorionic gonadotrophin (HCG) 46 hours later. These females were then mated with CD1(ICR) or BDF1 studs. Insemination was verified the next morning by the presence of a copulatory plug; this day was considered E0.5. Mouse zygotes were recovered by flushing the oviduct with M2 medium at E0.5. Adherent granulosa cells were removed from zygotes by incubating them in hyaluronidase (300 µg/mL in M2) (Sigma, H3506). The zona pellucida was removed at E3.5 using acidic Tyrode’s (Sigma, T1788), but was mostly not needed as the blastocysts on E4.5 hatched naturally by themselves and this stage was used predominantly. Embryos were cultured at 37°C in a CO_2_ incubator with an atmosphere of 20% O_2_ and 5% CO_2_. Embryos were grown from E0.5 to E4.5 in Continuous Single Culture Complete Media (CSCM-NXC) with HSA (Fujifilm, 90165) and later transferred to the enhanced in vitro implantation protocol from day 4 to 8. Bright field pictures of the embryos were taken every 24 hours using a Discovery V.20 stereoscope (Zeiss) or SMZ25/SMZ18 stereoscopes (Nikon).

Additional media components tested included the following: 0.22% sodium lactate (SIGMA, L7900), 100nM 3,3′,5-Triiodo-L-thyronine (T3) (SIGMA, T6397)), N2 (GIBCO, 17502048), B27 (GIBCO, 17504044), MEM Amino acids (SIGMA, M5550), CD Lipid concentrate (GIBCO, 11905031), Uteroglobin (Peprotech, 150-18), Human Insulin (SIGMA, I-1182), 100 ng/mL Human IGF-1 (Peprotech, 100-11), bFGF (R&D, 4114-TC), CHIR99021 (Axon Medchem, 1386), Y27632 (Axon Medchem, 1683), human LIF (Peprotech, 300-05), Geltrex (Invitrogen, A1413202), Rat Collagen Type I (Sigma, C3867) and Matrigel (BD, FAL356231). For culture optimization, different combinations and proportions of Fetal Bovine Serum (FBS), Rat Serum (RS), Human Cord Blood Serum (HCS), Human Adult Blood Serum (HAS), hormone supplements and basal media were tested.

A previously established in vitro culture (IVC) protocol^15^ was used as a control condition as follows: E3.5/E4.5 flushed blastocysts were cultured for 2 days in IVC1 media (Advanced DMEM/F12 (GIBCO, 12634010) containing 20% Fetal Bovine Serum (FBS) (GIBCO, 10270106 or 16141079) and supplemented with 1× GlutaMAX (GIBCO, 35050061), 1 mM sodium pyruvate (GIBCO, 11360070 or SIGMA, S8636), penicillin (25 units/ml)/streptomycin (25 mg/ml) (GIBCO, 15140163 or Biological industries– Sartorius, 03-031-1B), 1× ITS-X (ThermoFisher, 51500056), 8 nM ß-estradiol (Sigma, E8875), 200 ng/ml progesterone (Sigma, P0130), and 25 mM N-acetyl-L-cysteine (Sigma, A7250). After 48 hours (culture day 6), media was replaced with IVC2, which is identical to IVC1, but contains 30% KSR (Invitrogen, 10828-028) instead of 20% FBS. Embryos were cultured in IVC2 for up to 3 days (until day 9), replacing half of the media on a daily basis. Enhanced IVC, an alternative IVC media, which in combination with EUCM can support growth to E9.5 days of development (although with lower efficiency than CMRL-based media) was also standardized and used in this study. Enhanced IVC consisted of the same basal components and supplements of IVC2 media with several modifications: 30% KSR was replaced by 30% of HCS, plus 11 mM HEPES (GIBCO), 0.22% sodium lactate (Sigma), N2 (1×), B27 (0.5×) (GIBCO) and 75 nM 3,3′,5-Triiodo-L-thyronine (Sigma, T6397). At day 8, eIVC2 was replaced by EUCM. Half of the media volume was refreshed every 24 hours until the last day of static culture (day 10). For further development, the shaker and roller culture were used as described for PRI-ExUt (**Extended Data Fig. 1**) with EUCM.

### Post-implantation in utero control embryo dissection

Mouse embryos were obtained from non-hormone primed pregnant mice sacrificed by cervical dislocation from E5.5 to E9.5. Subsequently, embryos were dissected out from the uterus in PBS. The embryos were carefully dissected from the decidua and parietal yolk sac, leaving the intact ectoplacental cone attached to the egg cylinder, as previously described^3,4^. Briefly, the decidua was isolated from the uterine tissue and opened by introducing the forceps adjacent to the embryo in parallel to its proximal-distal axis and subsequently opening the forceps. Afterwards the embryo was grasped from the decidua using the forceps, and the parietal yolk sac was peeled off the embryo using two 55 forceps (Dumont, 11255-20).

### Stem cell lines

The following mouse ESC lines were used: BVSC N2 (Mixed BDF1 X B6 background; Male ESC line), WT V6.5 ESCs (C57B6/129sJae; Male ESC line), KH2-WT ESC (Male ESC line) line carrying both M2RtTa allele in the Rosa26 locus and a modified Col1 locus with FRT site to efficiently insert Tet-ON regulated alleles as previously described. G4 ESC and R1 ESC (with Oct4-GFP reporter) lines (male ESC lines; 129XC57BL/6 F1 hybrid genetic background strain) (kindly provided by A. Nagy), and V6.5 Triple Reporter #2 (OCT4-TagBFPnls, CDX2-H2B GFP and H2B-tdTomato).

### Naive ESC and other stem cell line in vitro culture conditions

Golden stocks of mouse ESCs and iPSCs were cultured on feeder layer of irradiated mouse embryonic fibroblast (MEFs) in conventional Fetal Bovine Serum (FBS) based mouse ES medium (Serum/LIF) that consists of 1x DMEM (Gibco, 41965), supplemented with 20% FBS (Heat inactivated and filtered, Gibco, 10270-106), 1 mM GlutaMAX (Gibco, 35050061), 1% penicillin/streptomycin (Biological Industries — Sartorius 03-031-1B), 1% sodium pyruvate (Biological Industries – Sartorius 02-340-1B), 0.1 mM beta-mercaptoethanol (Thermo 31350010), 1% non-essential amino acids (Biological Industries – Sartorius 01-340-1B), 20 ng/ml recombinant human LIF (in-house prepared). Serum/LIF conditions were used to maintain the golden stocks, as these conditions do not cause global erosion and loss of imprinting as seen upon long term expansion in serum free 2i/LIF conditions. For converting the cells into naive ESCs in 2i/LIF conditions, ESCs were grown with the serum-free chemically defined N2B27-based media: 240 ml Neurobasal (Thermo 21103049), 240 mL DMEM-F12 (Thermo, 21331), 5 ml N2 supplement (in-house prepared), 5 ml B27 supplement (in-house prepared), 1 mM GlutaMAX (Gibco, 35050061), 1% non-essential amino acids (Biological Industries — Sartorius 01-340-1B), 0.1 mM β-mercaptoethanol (Thermo 31350010), 1% penicillin-streptomycin (Biological Industries — Sartorius 03-031-1B), 1% sodium pyruvate (Biological Industries – Sartorius 02-340-1B). Naive 2i/LIF conditions for murine PSCs consist of 20 ng/ml recombinant human LIF (in-house made), small molecule inhibitors CHIR99021 (3 μM, Axon MedChem 1386) and PD0325901 (1 μM, Axon MedChem 1408). For TF-SEM generation, murine naive PSCs (either in 2i/LIF or Serum/LIF as indicated) were expanded on 0.2% gelatin-coated plates. ESCs and iPSCs were cultured under 2i/LIF conditions for at least three passages before being used for experiments. The cells grown with 2i/LIF media were used up to 10 passages to prevent the accumulation of chromosomal abnormalities and global loss of imprinting naturally occurring in 2i/LIF conditions. The ESCs/iPSCs were passaged by dissociation with TrypLE (Gibco, 12604054) or Trypsin-EDTA 0.25% every 3-4 days.

### Transgene-Free Naive PSC-derived Embryo Model Generation (TF-SEM)

To generate TF-SEMs via the rapid induction regimen, 2i/LIF conditions were used as starting conditions (unless stated otherwise regarding Serum/LIF), and initially the extraembryonic lineages were induced with Multi-Lineage Promoting Media 1 (MLPM-1) and afterward by Multi-Lineage Promoting Media 2 (MLPM-2). MLPM-1 consists of N2B27 media (240 ml Neurobasal (Thermo, 21103049), 240 mL DMEM-F12 (Thermo, 21331), 5 ml N2 supplement (in-house prepared), 5 ml B27 supplement (in-house prepared), 1 mM GlutaMAX (Gibco, 35050061), 1% non-essential amino acids (Biological Industries — Sartorius 01-340-1B), 0.1mM β-mercaptoethanol (Thermo 31350010), 1% penicillin-streptomycin (Biological Industries — Sartorius 03-031-1B), 1% sodium pyruvate (Biological Industries – Sartorius 02-340-1B)) with CHIR99021 (6μM, Axon MedChem, 1386), REPSOX (5μM, Sigma R0158), AM580 (0.025 μM,

Sigma A8843), and sodium butyrate (NaB) (0.5mM, Sigma B5887). Please note that LIF (STAT3 activator) had to be excluded from MLPM-1 (the first 24 hours of the rapid induction regimen). For SEM formation, naive mouse ESCs were dissociated with TrypLE (and not Trypsin) for 5 minutes at 37°C. TrypLE was diluted with MEF media and cells were centrifuged and supernatant was discarded. The cells were then split and seeded onto 0.2% gelatin coated plates at ratios of 1:5-1:10 depending on the initial confluency, in MLPM-1 supplemented with Y-27632 (5μM, AXON MEDCHEM) for 24 hours. After 24h treatment with MLPM-1, media was changed to MLPM-2 for 24h. MLPM-2 consists of N2B27 base media (240 ml Neurobasal (Thermo, 21103049), 240 mL DMEM-F12 (Thermo, 21331), 5 ml N2 supplement (in-house prepared), 5 ml B27 supplement (in-house prepared), 1 mM GlutaMAX (Gibco, 35050061), 1% non-essential amino acids (Biological Industries — Sartorius 01-340-1B), 0.1 mM β-mercaptoethanol (Thermo 31350010), 1% penicillin-streptomycin (Biological Industries — Sartorius 03-031-1B), 1% sodium pyruvate (Biological Industries – Sartorius 02-340-1B)) with CHIR99021 (3μM, Axon MedChem 1386), REPSOX (5 μM, Sigma R0158), AM580 (0.025 μM, Sigma A8843), recombinant human LIF (20 ng/ml). Please note that NaB must be excluded from MLPM-2. At the end of MLPM-2 treatment, the cells were dissociated with TrypLE for 5 mins at 37°C into single cells. After centrifugation (1300 rpm for 3 minutes), the cells were resuspended with multi-lineage maturation media (MLMM) supplemented with Y-27632 (Axon Medchem 1683, 5μM). MLMM consists of 1x DMEM (Gibco, 41965) base media with Activin A (20ng/ml, Peprotech 120-14E), human recombinant FGF4 (25ng/ml, Peprotech 100–131) (which can replace FGF2 to support primed pluripotency, yet is also efficient at enhancing TSC identity), XAV939 (2μM, Sigma X3004), TRULI (2μM, Sigma, SML3634), Heparin (1μg/ml, Sigma H3149), ROCK inhibitor Y-27632 (5μM, Axon Medchem 1683), L-ascorbic acid 2-phosphate (Vitamin C) (57.91μg/ml, Sigma, A8960), 1 mM GlutaMAX (GIBCO, 35050061), 20% heat inactivated FBS, 1% ITS-G supplement (Thermo, 41400045), 1% non-essential amino acids (Biological Industries — Sartorius 01-340-1B), 0.1 mM β-mercaptoethanol (Thermo, 31350010), 1% penicillin-streptomycin (Biological Industries — Sartorius 03-031-1B), 1% sodium pyruvate (Biological Industries – Sartorius 02-340-1B). Then, the cells were passed through a 70 μm cell strainer to remove any potential clumps. The cells were counted to determine the volume necessary for seeding 30 cells/microwell into the wells of 24 well plate of AggreWell 400 microwell plates (STEMCELL Technologies 34415), which were pretreated and prepared for use according to manufacturer’s instructions. The cells (36000 cells per well) were seeded in 1 ml MLMM in individual wells of 24-well AggreWell 400 plates. After 24h, half of the MLMM was gently replaced with fresh MLMM for an additional 24h. Then MLMM media was completely removed, and to remove any traces of MLMM, the aggregates were washed once with Aggregation media (AM) (DMEM supplemented with 20% heat inactivated FBS, 1 mM GlutaMAX (GIBCO, 35050061), 1% non-essential amino acids (Biological Industries — Sartorius 01-340-1B), 0.1 mM β-mercaptoethanol (Thermo 31350010), 1% penicillin-streptomycin (Biological Industries — Sartorius 03-031-1B), 1% sodium pyruvate (Biological Industries – Sartorius 02-340-1B)), and grown for 24h in AM. The next day, 500 μl of AM media is withdrawn and 1.5 ml EUCM2 is added. EUCM2 consists of^2^: 30% heat inactivated FBS (Sigma, F7524), 1% ITS-X supplement (Thermo, 51500-056), extra-added 1 mM sodium pyruvate (Biological Industries – Sartorius 02-340-1B), 1 mM GlutaMAX (GIBCO, 35050061), extra added 1 mg/ml D(+)-Glucose Monohydrate (J.T. Baker, 0113) (e.g. add 500mg per 500mL media), 1% penicillin-streptomycin (Biological Industries — Sartorius 03-031-1B), 8 nM β-estradiol (Sigma, E8875), 200 ng/ml Progesterone (Sigma, P0130), 25μM N-acetyl-L-cysteine (Sigma, A7250) and optionally: T3 100nm (Sigma, T6397) and 0.5% CMRL media (GIBCO 11530037). On day 6, 1.5 ml of EUCM2 media is gently withdrawn from the Aggrewells. The structures were transferred as previously described using disposable plastic Pasteur pipettes to 6 well suspension culture plates (Greiner, 657185) containing 3 ml fresh EUCM2 media per well. The plates were placed on a shaker rotating at 60 rpm (Thermo 88881102 and 88881123) in tissue culture incubator at 37°C, 20%O2 and 5%CO2. After 24h, high-quality egg cylinder-shaped TF-SEMs were picked and transferred to glass culture bottles (50 structures per bottle) containing 4 ml freshly prepared ex utero culture media (EUCM).^9^ EUCM was prepared as previously described (consists of 25% DMEM (GIBCO 11880 – DMEM, low glucose, with pyruvate, no glutamine, no phenol red (and no HEPES)), 50% rat serum (Harlan), and 25% human adult serum (HAS) (in-house prepared) supplemented with a final concentration of 1x GlutaMAX (GIBCO, 35050061), penicillin (25 units/ml)/streptomycin (25 mg/ml) (GIBCO, 15140163 or Biological industries– Sartorius, 03-031-1B), extra added 1 mM sodium pyruvate (Biological industries, 030421B), extra added 4 mg/mL of D-glucose (J.T. Baker), and added HEPES (11mM final concentration) (Gibco, 15630056)). The bottles were placed on the electronically controlled ex utero rolling culture system, rotating at 30 revolutions per minute at 37°C in 21% O2, 5% CO2 and all other parameters as previously described. Every 24 hours, 3.5 ml EUCM media was removed, and replaced with 4 ml fresh EUCM media. On day 8, only the properly developed E7.5 SEMs were manually picked for further growth under stereomicroscope. On day 10, properly developed E8.5-like TF-SEMs were picked for processing and analysis. All cell line and TF-SEM maintenance, induction and expansion were done in 37°C, 5%CO2 and 20%O2. All media for TF-SEM generation is pre-heated and pre-gassed (pre-equilibrated) for at least 1hour (e.g. EUCM is pre-heated for at least an hour by placing it inside a glass bottle on the rotating culture; AM and EUCM2 are preheated in regular TC incubators). We would like to state that despite the many optimization tests executed herein, we cannot exclude that alternative induction, aggregation or growth conditions might yield similar or enhanced results relative to the ones reported herein. For some PSC lines, TF-SEM formation efficiency was higher if cells were expanded in naive FBS/LIF compared to naive 2i/LIF starting conditions, while an opposite pattern could be observed for other lines. While nearly all PSC lines tested were able to generate up to E7.5 TF-SEMs, some lines still failed to generate E8.5 TF-SEMs.

### Maintenance of Embryo Derived Trophoblast Stem Cells (eTSCs)

eTSCs were cultured with the TXV media conditions on MEF layers. TXV media consists of DMEM/F12 (without HEPES, Thermo, 21331), L-ascorbic acid-2-phosphate (64μg/ml, Sigma, A8960), sodium bicarbonate (543 mg/liter, S6761, Sigma Aldrich), insulin (19.4 mg/liter, Sigma I-1882), holo-transferrin (10.7 mg/liter, T0665, Sigma), sodium selenite (14 μg/liter, Sigma, S5261), FGF4 (25 ng/ml, Peprotech, 100-31), heparin (Sigma, H3149, 1 μg/ml), TGFB1 (Peprotech, 100-21, 2 ng/ml), human BMP7 (30 ng/ml, 120-03P), IL-11 (50 ng/ml, Peprotech, 200-11), Activin A (50 ng/ml, Peprotech, AF-120-14E), LPA (5 nM, Tocris, #3854), 8Br-cAMP (200 μM, Sigma, B7880). Y-27632 (2 μM, Axon Medchem 1683) was added for 24h after passaging. Due to the instability of the 8Br-cAMP, 8Br-cAMP was added fresh during daily media changes.

### Generation and Maintenance of Primitive Endoderm Stem Cells (PrESCs) from Mouse ESCs

To generate PrESCs, 2i/LIF mouse naive ESCs were first washed with PBS and then cultured with PrESC media with MEF cells.

### Generation of V6.5 Triple Reporter #2 mESC Line

For making CRISPR constructs gRNA sequences targeting the stop codons of the relevant genes were designed using the CRISPOR tool and cloned into the px330 vector. V6.5 mESCs were electroporated with mGata6-H2B-tdTomato_puro, mCdx2_H2B_GFP_neo and the CRISPR constructs. After double antibiotic selection cells were co-transfected with mOct4_tagBFPnls donor and CRISPR constructs, and with Flippase coding plasmid to remove selection cassettes from the Gata6 and Cdx2 genes. A few days later NLStagBFP positive cells – that were mostly correctly targeted with Oct4_tagBFPnls – were sorted out and seeded at clonal density. The resulting clones were isolated, expanded and verified by PCR with external primers for 3’ and 5’ ends, and by FACS.

### Flow Cytometry

Flow cytometry analysis and sorting were performed on a BD FACS-Aria III. Cells were harvested with TrypLE. Cells were initially gated using FSC-A versus SSC-A to select the main cell population and exclude debris. CDX2–GFP, GATA6–tdTomato and OCT4–TagBFP reporter fluorescence was evaluated using pairwise fluorescence plots. Positive and negative populations were defined using thresholds established with WT V6.5 and V6.5 triple reporter #2 cultured under 2i/LIF conditions. Representative scatter gates and fluorescence quadrant gates for cells cultured in 2i/LIF, PrESC conditions and following 48 h of induction are shown in **Extended Data Fig. 17B**. Flow cytometry data was analyzed and visualized with FlowJo v10.7.

### Mouse blastoids generation following the previously established protocol #1

Mouse blastoids were generated following previously established protocol #1^41^. Briefly, ESCs were aggregated (6 cells/microwell). The cells were seeded in S/L that consists of 1x DMEM (Gibco, 41965), supplemented with 10% FBS (Heat inactivated and filtered, Gibco, 10270-106), 1 mM GlutaMAX (Gibco, 35050061), 1% penicillin/streptomycin (Biological Industries — Sartorius 03-031-1B), 1% sodium pyruvate (Biological Industries – Sartorius 02-340-1B), 0.1 mM beta-mercaptoethanol (Thermo 31350010), 1% non-essential amino acids (Biological Industries – Sartorius 01-340-1B), 10 ng/ml recombinant human LIF (in-house prepared) further supplemented with 5 μM Y27632 (Axon Medchem, 1683) in Aggrewell 400 plates (STEMCELL Technologies, 34415). The next day compact aggregates were observed. On top of the formed aggregates, eTSCs are seeded at a ratio of 12 cells/microwell in TX media with supplements. However, before the seeding, MEFs were excluded by culturing the cell mix with MEF media on 0.2% gelatin-coated plates for 20 minutes. After MEF depletion, TSCs were seeded. TX media with supplements consists of DMEM/F12 (without HEPES, Thermo, 21331), L-ascorbic acid-2-phosphate (64μg/ml, Sigma, A8960), sodium bicarbonate (543 mg/l, S6761, Sigma Aldrich), insulin (19.4 mg/l, I-1882, Sigma), holo-transferrin (T0665, Sigma), sodium selenite (14 μg/l, S5261, Sigma), FGF4 (25 ng/ml, Peprotech, 100-31), heparin (Sigma, H3149, 1 μg/ml), TGFB1 (Peprotech, 100-21, 15 ng/ml), 8Br-cAMP (1 mM, Sigma, B7880), IL-11 (30 ng/ml), CHIR99021 (3μM, Axon MedChem, 1386) and Y-27632 (20 μM, Axon Medchem 1683). On Day 2, wells are supplemented with fresh 1 mM 8BR-cAMP. On Day 3, the cavitated structures can be observed. On Day 4, the structures with correct blastocyst morphology were picked for processing further. For culturing the blastoids, PRI-ExUt culture method was employed without any changes to the protocol applied to natural blastocysts derived from zygotes described before.

### Mouse blastoid generation following the previously established protocol #2

For the generation of blastoids following the previously established protocol #2^60^, the 2i/LIF mouse naive ESCs that were cultured on gelatin were seeded into C++6A media that consists of N2B27 media supplemented with CHIR99021 (9 μM, Axon MedChem 1386), REPSOX (10μM, Sigma R0158), AM580 (0.05 μM, Sigma A8843) and cultured for 24h. Next, the cells were cultured with CL6A media medium consisting of N2B27 media supplemented with CHIR99021 (3 μM, Axon MedChem 1386), REPSOX (10μM, Sigma R0158), AM580 (0.05 μM, Sigma A8843), and 20ng/ml LIF (in-house made) for 36h. Following the sequential treatment with C++6A and CL6A, the cells were aggregated at a ratio of 25 cells/microwell to generate blastoids in blastoid medium consisting of 25% TSC basal medium (RPMI-1640 supplemented with 20% FBS, 1 mM GlutaMAX (Gibco, 35050061), 1% non-essential amino acids (Biological Industries — Sartorius 01-340-1B), 0.1 mM β-mercaptoethanol (Thermo 31350010), 1% penicillin-streptomycin (Biological Industries — Sartorius 03-031-1B), 1% sodium pyruvate (Biological Industries – Sartorius 02-340-1B), 25% N2B27 media, and 50% KSOM supplemented with Y-27632 (2 μM, Axon Medchem 1683), FGF4 (12.5 ng/ml, Peprotech, 100-31), BMP4 (5 ng/ml, Peprotech, 120-05ET), CHIR99021 (3μM, Axon MedChem, 1386), heparin (Sigma, H3149, 1 μg/ml), A83-01 (Axon Medchem, 1421). The media was changed every day until Day4. On Day4, the cavitated structures were collected for downstream analysis or culturing under PRI-ExUt culture. PRI-ExUt culture method was employed without any changes to the protocol applied to natural blastocysts derived from zygotes described before.

### Isolation of human umbilical cord blood serum and adult blood serum

Collection of human cord blood serum for ex utero culture of embryos was done as described previously^3^. For the purpose of this study, healthy pregnant women who were referred to the prenatal clinic at the Rambam Medical Center or at the University of Texas Southwestern Medical Center were asked to give their informed consent to have blood collected from their umbilical cord, as approved by a Rambam Medical Center Helsinki committee (#RMB-0452-15) and the UTSW IRB committee with Parkland Site Approval (#STU-112010-047). The source of each collection underwent full anonymization and was not identified by name or other designation, and the extracted serum was only used as described herein. Healthy women over the age of 18 and under 40, who gave their consent and were scheduled for caesarian section delivery by their obstetrician following a prenatal clinic visit, were eligible for cord blood collection. We excluded women who gave vaginal birth as well as women with any chronic illness or active medical conditions, including gestational diabetes or hypertension. On the day of scheduled caesarian delivery and to ensure fresh isolation and processing of serum, a team stood by for cord blood collection and serum extraction. Immediately upon delivery of the infant, the umbilical cord was double clamped 5-7 cm from the umbilicus and transected between the clamps. Blood was collected for the purpose of this study, only after the infant was removed from the field of surgery and umbilical blood was drawn for clinical tests as needed. To avoid any traces of hemolysis, blood was manually drawn by the obstetrician surgeon, using a large bore 14-gauge needle and a 50 mL syringe, directly from the umbilical vein while the placenta remained in situ. This was done to avoid any coagulation of blood before collection which could lead to traumatic hemolysis, and to take advantage of the enhanced blood flow generated by uterine contraction. Next, the collected blood was freshly and quickly distributed to 5 mL pro-coagulant sterile test tubes (Greiner Bio-One, Z Serum Sep Clot Activator, #456005) and cooled to 4°C for 15 minutes, to allow full coagulation. Then, coagulated test tubes were centrifuged at 2500G for 10 minutes in a cooled 4°C centrifuge. Any tube that showed signs of hemolysis (such as light pink-red colored serum) was discarded. The separated serum (yellow-colored) was collected using a pipette and filtered through a 0.22 µm filter (Nalgene, #565-0020) and then inactivated in 55°C bath for 30-45 minutes. The inactivated serum was next distributed to aliquots and placed in a -80°C freezer for storage for up to six months. Shipping temperature was kept at -70°C using dry ice and any thawed serum was refrozen once. At the Weizmann Institute, human adult blood serum was collected in house from healthy adult donors (avoid donors with elevated blood lipid level or BMI>30) and freshly prepared with the same protocol (IRB# 2710-1) described for umbilical cord blood serum, while at Janelia Research Campus commercial human adult blood serum specifically for embryo culture was acquired from Human Cell Biosciences (cat# FP-006-C100). In all cases, human serum was thawed, kept at 4°C for maximum 24 hours, and refrozen once.

### Morphological evaluation of mouse embryo development

Embryos were imaged daily using a Discovery V.20 stereoscope (Zeiss) or SMZ25/SMZ18 stereoscopes (Nikon). Developmental stage of the embryos was determined according to Downs & Davies^84^. Assessment of appropriate embryo development was performed based on formerly established morphological features^85–89^ from E5.5 to E9.5, as previously described^3,15^. All flushed viable zygotes were cultured without selection. At E3.5/E4.5, only properly developed blastocyst with a clear inner cell mass, trophoblast and blastocoel were selected for in vitro implantation. Subsequently, egg cylinders resembling to E5.5 post-implantation embryos arose at culture day 8, and only those with cylinder shape, clear extraembryonic ectoderm (ExE), cup-shaped epiblast, and visceral endoderm (VE) surrounding both compartments we considered as properly developed. The same features were considered for E6.5 stage embryos (commonly culture day 9), in addition to the presence of cavities in the embryonic and extraembryonic compartments, which can be unified to form the pro-amniotic cavity, as well as noticeable radial symmetry breaking in the epiblast indicated by the primitive streak. At day 10 (∼E7.5), properly developed embryos generally ranged from the late-streak stage to neural plate stage. The amnion and the chorion were formed, generating three cavities in the embryo (amniotic, exocoelomic and ectoplacental cavities), the allantoic bud may be present at the base of the primitive streak, and the anterior ectoderm begins to form the future neural groove. At culture day 11, correctly developed embryos vary between developmental stages equivalent to E8.0/E8.5 days of in utero development (headfold stage to somitogenesis; Theiler stage 12). Headfold stage embryos present well-defined headfolds, elongated allantois, and lack foregut invagination. At the somite-stages, embryos are curved dorsally and present ∼2-8 pairs of somites, closing but unfused neural folds, the yolk sac developed to enclose the embryo and presents visible blood islands; blood circulated though the yolk sac primitive capillary plexus and through the embryo proper; the allantois started to fuse with the chorion, and the invaginating foregut pocket and beating heart are formed in the frontal region of the embryo. 24-36 hours later (day 12), PRI-ExUt embryos match E9.5 in utero controls. At this stage, embryos present 16-24 somites (Theiler stage 13 and 14), axial turning leads to the dorsal part of the embryo to face outside, the forelimb buds are clearly present, highly organized yolk sac blood vessels with circulating blood are observed, three-chambered heart, the cranial part of the neural tube is closed while the posterior neuropore is closing with small opening remaining, the brain has been regionalized into forebrain, midbrain and hindbrain, and the otic pit is present and separated from the epidermis; the maxillary process, mandibular and hyoid branchial arches are visible, and the optic vesicle is established. For calculating efficiency of ex utero culture throughout the manuscript, the total number of embryos assessed per condition in every sampled timepoint is indicated. The efficiency percentages in main figures are calculated based on the starting number of embryos. For the effective efficiency percentages after picking the correct structures, the number of correct embryos at the specified stage is determined as the denominator for the efficiency calculations. Embryos dissected, fixed or moved to other conditions at any point during the time-course, are subtracted from the total where relevant.

The main developmental abnormalities observed in the fraction of cultured defective embryos are the following: At E5.5, lack of extraembryonic ectoderm (rounded epiblast surrounded by VE only). At E6.5, underdeveloped epiblast, folded or double epiblast, round epiblast with no apparent symmetry breaking. E7.5, same phenotypes as for E6.5, as well as expansion of extraembryonic tissues without epiblast growth. At E8.5 commonly observed defects are either smaller neural folds, lack of somites, myocardial edema, shorter antero-posterior axis, underdeveloped posterior region, or development of the yolk sac without the embryo proper. At the time equivalent to E9.5, the most observed defects are failure of axial turning and lack of further body growth, pericardial effusion, intraembryonic hemorrhage, limited growth of the forebrain/facial regions, and lack of connection of the umbilical cord to the ectoplacental cone.

### Whole-mount immunostaining of E5.5 - E8.5 mouse embryos

Ex utero and in utero dissected embryos (without Reichert’s membrane for E5.5-E7.5 embryos, or without yolk sac and amnion for E8.5/E9.5 embryos) were washed once with 1× PBS and fixed with 4% PFA EM grade (Electron microscopy sciences, 15710) in PBS at 4°C over-night. Embryos were then washed in PBS for 5 minutes 3 times, permeabilized in PBS with 0.5% Triton X-100/0.1M glycine for 30 minutes, blocked with 10% normal donkey serum/0.1% Triton X-100 in PBS for 1 hour at room temperature (RT), and incubated over-night at 4°C or room temperature (RT) with primary antibodies, diluted in blocking solution. After, embryos were rinsed 3 times for 5 minutes each in PBS/0.2% TritonX-100, incubated for 2 hours at RT with secondary antibodies diluted 1:200 in blocking solution (all secondary antibodies were from Jackson ImmunoResearch or Abcam), counterstained with DAPI (1 µg/ml in PBS) for 10 minutes, and washed with PBS for 5 minutes 3 times. For details on primary antibodies and imaging see antibody dilutions section.

### Whole mount immunostaining and clearing of E9.5 mouse embryos

Clearing of embryos E9.5 was performed using iDISCO^90^ or SHIELD/SWITCH^91^ (LifeCanvas) with some modifications. Embryos were dissected in PBS and fixed in 4% PFA EM grade (Electron microscopy sciences, 15710) at 4°C overnight. For iDISCO, after blocking, embryos were incubated with primary antibodies diluted in PBS/0.2%Tween-20 with 10 µg/ml heparin (PTwH)/5% DMSO/3% Donkey Serum at 37°C for 24 hours. Afterwards, samples were washed in PTwH for 24 hours (15 minutes, 30 minutes, 1 hour, 2 hours, and overnight washes) and incubated with adequate secondary antibodies (1:200) plus DAPI (1 µg/ml) diluted in PTwH/3% Donkey Serum at 37°C for 24 hours. Embryos were then washed in PTwH for one day (5 min, 15 min, 30 min, 1 hour, 2 hours, and overnight washes), dehydrated in methanol/H_2_O series (1 hour each) and then incubated overnight in 100% methanol. Embryos were incubated in 66.6% DCM/33.3% methanol on shaker for 3 hours, followed by 100% DCM (Sigma; 270997) for 5 min, and finally cleared and stored in Benzyl Ether (Sigma; 108014) or Ethyl Cinnamate (Sigma; 112372). For SHIELD/SWITCH, embryos were incubated overnight at 4°C with gentle shaking in freshly prepared SHIELD-OFF (LifeCanvas) solution. The next day, samples were transferred to SHIELD-ON (LifeCanvas) solution and incubated overnight at 37 °C with gentle agitation, followed by two brief washes in PBS (5 min each). Samples were then delipidated overnight in Delipidation Buffer (LifeCanvas) at 45°C with shaking. After delipidation, embryos were blocked for 24 hours at 37°C in PBSTG (PBS containing 0.5% Triton X-100 and 0.2% gelatin). Primary antibody incubation was carried out for 1 day at 37°C in PBSTG with continuous shaking, followed by 1 day wash (5 min, 15 min, 30 min, 1 hour, 2 hours, and overnight washes) in 0.1% Triton X-100 in PBS. Samples were incubated with adequate secondary antibodies (1:500 in PBSTG, filtered through a 0.22µm syringe filter) for 1 day at 37°C. Afterwards, embryos were washed three times in PBS (5 min each), and index-matched in EasyIndex (LifeCanvas) overnight at room temperature. All incubations were performed in the dark unless specified otherwise.

### Antibody dilutions for immunofluorescence

Mouse monoclonal anti-Myosin Heavy Chain II (MF-20) (R&D Systems, MAB4470; 1:500); Goat anti-SOX2 (R&D Systems, AF2018; 1:200); Rabbit anti-SOX9 (Millipore, AB5535; 1:200); Mouse anti-βIII-tubulin (Tuj1) (Covance, MMS-435P; 1:1000); Rabbit anti-GATA4 (Abcam, ab84593; 1:200); Rabbit anti-FOXA2 (Abcam, ab40874; 1:100); Goat anti-SOX17 (R&D Systems, AF1924; 1:100); Rabbit anti-BRACHYURY (Cell Signaling Technology, 81694; 1:100); Goat anti-BRACHYURY (R&D Systems, AF2085; 1:100; Rabbit anti-CDX2 (Abcam, ab76541; 1:100); Mouse anti-OCT3/4 (Santa Cruz Biotechnology, sc-5279; 1:100); Goat anti-OTX2 (R&D Systems, AF1979; 1:100); Goat anti-TFAP2C/AP2γ (R&D Systems, AF5059; 1:100); Rabbit anti-FOXG1 (Abcam, ab18259; 1:200); Goat anti-GATA6 (R&D Systems, AF1700; 1:100); Rabbit anti-EOMES/TBR2 (Abcam, ab23345; 1:50); Rabbit anti-NANOG (Bethyl, AF1997; 1:100); Goat anti-CDX2 (AF3665). Rhodamine-Phalloidin (PHDR1) (ThermoFisher, R415; 1:400) was used to label cell membranes.

### Confocal microscopy

Whole-mount immunofluorescence images were acquired either with a Zeiss LSM700 inverted confocal microscope (Carl Zeiss) equipped with 405 nm, 488 nm, 555 nm and 635 nm solid state lasers, using an EC Plan Neofluar 10× air objective (numerical aperture 0.3), or with a Nikon AX inverted confocal microscope equipped with 8 laser lines (405 nm/445 nm/488 nm/514 nm/561 nm/594 nm/640 nm/730 nm) and a Galvano scanning system, using 4× air objective (numerical aperture 0.2) or 10× air objective (numerical aperture 0.45) Plan Apo Lambda D objectives. Images were acquired at 1024×1024 resolution. All images were acquired within the following range of parameters: Laser power Zeiss LSM700: 405 nm: 5-20%; 488 nm: 5-30%; 555 nm 10-40%; 635 nm: 30-80%. Gain ranged from 350 to 600. Laser power Nikon AX: 405 nm: 0.5-15%; 445 nm: 0.5-20%; 488 nm: 0.5-20%; 514 nm: 0.5-30%; 561 nm 0.5-30%; 594 nm 0.5-30%; 640 nm: 0.5-50%; 730 nm: 1-60%. Gain ranged from 0.5 to 40. Pixel size was 1.25 µm with a z-step of 10-12 µm when using the 10× objective (Zeiss). Pixel size was 1.14-4.31 µm with a z-step of 5-10 µm when using the 4× and 10× objectives (Nikon). For confocal imaging, iDISCO/SWITCH cleared embryos were mounted in 35 mm glass bottom dishes (In Vitro Scientific, D35201.5N), employing ethyl cinnamate (Sigma, 112372) as imaging solution for iDISCO or EasyIndex (LifeCanvas Technologies, EI-500-1.52) for SWITCH. Images and maximum intensity projections were processed using Zen 2 blue edition software (Carl Zeiss), NIS Elements (Nikon) and Adobe Photoshop 26.11.0. Imaris (Bitplane) software was used to create 3D reconstructions and animations of the imaged embryos.

### Assessment of mouse embryo and embryo-model length

Morphometric measurements were performed using the NIS Elements software (Nikon) or the CellSens Entry 1.18 software (Olympus). Length of the antero-posterior axis was measured for natural embryos between E6.5 to E8.5. The crown-rump length (the longest straight line from the cranial to the caudal end of the body) after removing the yolk sac was measured for embryos at E9.5 or equivalent stages. Length of cultured embryos was compared with CD1(ICR) in utero dissected embryos at matched embryonic stages.

### Longitudinal live imaging of peri-implantation embryo development

Longitudinal bright-field imaging of mouse embryos was performed using an Olympus IX85 inverted microscope equipped with a Hamamatsu ORCA-Fusion BT scientific CMOS (sCMOS) camera. E4.5 blastocysts were transferred into 8-well ibiTreat plastic µ-Plates (ibidi; 3–4 blastocysts per well) containing CMRL-based implantation medium, as described above. Culture plates were maintained on the microscope stage using an integrated Okolab environmental incubation system equipped with an H203 K-Frame stage-top chamber and an electronically controlled gas mixer. Temperature was maintained at 37 °C under an atmosphere containing 5% CO₂ and 21% O₂ throughout imaging. Embryos were maintained under sequential PRI-ExUt implantation culture conditions, with medium composition and serum supplementation adjusted according to the corresponding culture day, as described above. Half of the culture medium was replaced every 24 h throughout the imaging period. Bright-field images were acquired every 15 min for 5 consecutive days using Olympus CellSens Imaging Software. Acquired images were processed and assembled into time-lapse sequences using CellSens software, allowing longitudinal visualization of blastocyst attachment, trophoblast outgrowth and progressive egg-cylinder morphogenesis (Supplementary Video1).

### 10X Genomics single cell RNA-sequencing

For PRI-ExUt embryo characterization, PRI-ExUt embryos at day 10 (2 embryos, pooled, 10000 cells targeted), 11 (1 embryo, 2 samples, 20000 cells targeted) & 12 (1 embryo, 2 samples, 20000 cells targeted) of the protocol, along with E9.5 in utero control embryo (1 embryo, 2 samples, 20000 cells targeted) were selected and harvested for single cell RNA sequencing. In addition, single-cell RNA sequencing was generated from PRI-ExUt embryos at day E4.5+24h, +48h and +72h of the protocol (40-60 embryos each), and from cell cultures of V6.5 ESC line grown for 2 passages on MEFs in either 2i/LIF ESC media or in PrESC media^75^.

All embryos were processed with their extraembryonic compartments without any dissection. All embryos were dissociated with Trypsin-EDTA solution A 0.25% (Biological Industries; 030501B). Trypsin was neutralized with media including 10% FBS and cells were washed and resuspended in 1x PBS with 400 µg/ml BSA. Cell suspension was filtered with a 100 µm cell strainer to remove cell clumps. A percentage of cell viability higher than 90% was determined by trypan blue staining. Cells were diluted at a final concentration of 1000-2000 cells/µL. The same procedure was repeated for the analysis of PrESCs. However, MEF depletion was performed to exclude the MEF cells for the analysis by having the cell suspension in MEF media and 0.2% gelatin coated plates. scRNA-seq libraries were generated using the 10x Genomics Chromium v4 Dual Index system and sequenced using Illumina NovaSeq 6000 platform according to the manufacturer’s instructions.

### Single cell RNA-seq data processing

A total of 38 samples were used for analysis, out of which 10 were generated for this paper while others were taken from the indicated previously published ^68,69,73,75^ papers as detailed in **Supplementary Table S1**. 10X Genomics data analysis was performed using Cell Ranger 9.0.0 software (10X Genomics) for pre-processing of the raw sequencing data. Reads were aligned to 10X mm10-3.0.0 genome with “—include-introns true” flag. One sample (ExUt_E85_1e_d0) was processed with --force-cells=30000 flag, to limit the number of cells and reduce noise. Samples that were analyzed together were aggregated with Cell Ranger aggr software. Some of the downstream analysis was done with R Seurat 5.0.0 as described below: Cells were filtered to include cells that express between 2000 and 8000 genes and a maximum of 10% mitochondrial reads. In E45+24/48/72h samples minimal gene number was adjusted to 500. Sample integration and normalization was done in Seurat by anchoring all individual samples, and log-normalization using a scale-factor of 10,000. The top 2,000 variable genes were identified by the variance stabilizing transformation method and subsequently scaled and centered. Principal Components Analysis was performed for dimensional examination using the ‘elbow’ method. The first 15 dimensions showed majority of data variability. Therefore, UMAP dimensional reduction was performed on the first 15 dimensions in all samples. Clusters were detected using Seurat FindClusters function, with resolution parameter that varies between 0.1 and 0.5, dependent on the dataset. V6.5 ESCs in 2i/LIF or PrESC media samples had an additional filtering step to exclude MEF cells from further analysis.

### Downstream Analysis of scRNA-seq based data

UMAPs, Dot plots, stacked bar plots, scatter-plots analysis, Feature plots and Violin plots analysis were done with the use of the following R packages: Seurat, ggplot2, ggtern, Matrix, stringr, dplyr. Some code parts and code documentation were written with the assistance of Claude (Opus 4.8, Anthropic). Similarity Score (Extended Data Fig 7X is calculated from Seurat Module Score S, with min-max rescaling as following: S-norm = (S - min(S)) / (max(S) - min(S)) × 1.5. The code is available in GitHub: https://github.com/hannalab/PRI_EXUT.

### Software and statistical analysis

Embryos were randomly allocated when placed in the different growth conditions and samples were processed in an arbitrary order, but formal randomization techniques were not used. Samples sizes were not pre-determined based on statistical power calculations but were based on our experience with these assays. For most experiments, the minimum number of independent experiments was 3, with some exceptions as indicated in the figure legends. Other experiments were not randomized. The investigators were not blinded to allocation during experiments and outcome assessment, since there was no relevant scientific reason to do so. All statistical analysis were performed using the GraphPad Prism 11 software (La Joya, California). For embryology experiments, data on graphs indicates mean plus S.E.M. of a minimum of three independent experiments. Kolmogorov-Smirnov test was performed to check normal distribution of data before each statistical test. Significant difference between two samples was evaluated by unpaired two-sided Student’s t-test if data was normally distributed or Mann-Whitney test for non-normally distributed data. *p* < 0.05 was considered as statistically significant.

## Ethics Statement

All animal experiments were performed according to the Animal Protection Guidelines of the Howard Hughes Medical Institute Janelia Research Campus and the Weizmann Institute of Science and were approved by relevant HHMI Janelia IACUC (#24-0256.02), or Weizmann Institute IACUC (#04640724-3, #07901123-1, #03370423-1, #09521222-1). Human blood serum derivations and their use in this study were ethically reviewed and approved by a Rambam Medical Center Helsinki committee (#RMB-0452-15), the UTSW IRB committee with Parkland Site Approval (#STU-112010-047), and the Weizmann Institute for Science IRB and ESCRO committee (#2710-1, #1497-1).

## Declaration of generative AI and AI-assisted technologies in the manuscript preparation process

During the preparation of this work, the authors used Claude for language editing and fine formatting of the text to match journal style. Some code parts and code documentation were written by Noa Novershtern with the assistance of Claude (Opus 4.8, Anthropic). The authors reviewed and edited the output as needed and take full responsibility for the content of the published pre-print.

## Acknowledgements

We thank members of Hanna lab and RenewalBio Ltd. for helpful discussions.

## Data Availability

The scRNA-seq data generated in this study have been deposited in the **NCBI Gene Expression Omnibus (GEO).** The code used for scRNA-seq analysis and visualization is **available in GitHub.**

## Author Contributions

G.G. and A.Y. assisted on establishing the zygote to organogenesis ex utero culture protocol, performed embryology, immunostainings, imaging, and sequencing experiments for natural embryos and generated blastoids and cultured them. D.L., N.Z, I.S. and M.G. assisted in establishing the zygote to organogenesis ex utero culture protocol, performed embryology and immunostainings for natural embryos. L.W., A.Y., and C.R., performed library preparation and sequencing for 10X genomics. F.R., S.E., and J.D.L.O. assisted on embryo culture and immunostainings. I.M., N.K. and R.R.K., collected human cord blood and calibrated human cord serum production. N.N. performed bioinformatic analysis. A.A-C. conceived the idea for this project, established the zygote to organogenesis ex utero culture protocol and gas regulation systems/incubators, designed and conducted most embryology, sequencing and imaging on natural embryos, acquired funding, supervised executions of experiments, adequate data analysis, and wrote the manuscript. J.H.H. conceived the idea for this project, acquired funding, supervised executions of experiments, adequate analysis of data and wrote the manuscript.

